# A Chemical Probe of CARM1 Alters Epigenetic Plasticity against Breast Cancer Cell Invasion

**DOI:** 10.1101/591164

**Authors:** Xiao-Chuan Cai, Tuo Zhang, Eui-jun Kim, Ming Jiang, Ke Wang, Junyi Wang, Shi Chen, Nawei Zhang, Hong Wu, Fengling Li, Carlo C. dela Seña, Hong Zheng, Victor Vivcharuk, Xiang Niu, Weihong Zheng, Jonghan P. Lee, Yuling Chen, Dalia Barsyte, Magda Szewczyk, Taraneh Hajian, Glorymar Ibáñez, Aiping Dong, Ludmila Dombrovsky, Zhenyu Zhang, Haiteng Deng, Jinrong Min, Cheryl H. Arrowsmith, Linas Mazutis, Lei Shi, Masoud Vedadi, Peter J. Brown, Jenny Xiang, Li-Xuan Qin, Wei Xu, Minkui Luo

## Abstract

CARM1 is a cancer-relevant protein arginine methyltransferase that regulates many aspects of transcription. Its pharmacological inhibition is a promising anti-cancer strategy. Here **SKI-73** is presented as a CARM1 chemical probe with pro-drug properties. **SKI-73** can rapidly penetrate cell membranes and then be processed into active inhibitors, which are retained intracellularly with 10-fold enrichment for days. These compounds were characterized for their potency, selectivity, modes of action, and on-target engagement. **SKI-73** recapitulates the effect of CARM1 knockout against breast cancer cell invasion. Single-cell RNA-seq analysis revealed that the **SKI-73**-associated reduction of invasiveness act via altering epigenetic plasticity and suppressing the invasion-prone subpopulation. Interestingly, **SKI-73** and CARM1 knockout alter the epigenetic plasticity with remarkable difference, arguing distinct modes of action between the small-molecule and genetic perturbation. We therefore discovered a CARM1-addiction mechanism of cancer metastasis and developed a chemical probe to target this process.

## Introduction

Numerous biological events are orchestrated epigenetically upon defining cellular fates.(Atlasi and Stunnenberg, 2017; Berdasco and Esteller, 2019) Among key epigenetic regulators are protein methyltransferases (PMTs), which can render downstream signals by modifying specific Arg or Lys residues of their substrates with *S*-adenosyl-L-methionine (SAM) as a methyl donor cofactor.(Luo, 2018) Significant efforts have been made to identify the PMT-dependent epigenetic cues that are dysregulated or addicted under specific disease settings such as cancer.(Berdasco and Esteller, 2019) Many PMTs are implicated as vulnerable targets against cancer malignancy.(Kaniskan et al., 2018; Luo, 2018) The pro-cancerous mechanism of these PMTs can be attributed to their methyltransferase activities via individual or combined effects of upregulating oncogenes, down-regulating tumor suppressors, and maintaining cancer-cell-addicted homeostasis.(Berdasco and Esteller, 2019; Blanc and Richard, 2017) Pharmacological inhibition of these epigenetic events thus presents promising anti-cancer strategies,(Berdasco and Esteller, 2019) as exemplified by the development of the clinical inhibitors of DOT1L,(Bernt et al., 2011; Daigle et al., 2011) EZH2(Kim et al., 2013; Konze et al., 2013; McCabe et al., 2012; Qi et al., 2012; Qi et al., 2017), and PRMT5.(Bonday et al., 2018; Chan-Penebre et al., 2015)

Protein arginine methyltransferases (PRMTs) act on their substrates to yield three different forms of methylated arginine: asymmetric dimethylarginine (ADMA), symmetric dimethylarginine (SDMA), and monomethylarginine (MMA)---the terminal products of Type I, II and III PRMTs, respectively.(Blanc and Richard, 2017; Yang and Bedford, 2013) Among the important Type I PRMTs is CARM1 (PRMT4), which regulates multiple aspects of transcription by methylating diverse targets including RNAPII, SRC3, C/EBPβ, PAX3/7, SOX2/9, RUNX1, Notch1, p300, CBP, p/CIP, Med12, and BAF155.(Blanc and Richard, 2017; Hein et al., 2015; Vu et al., 2013; Wang et al., 2015; Wang et al., 2014; Yang and Bedford, 2013) The physiological function of CARM1 has been linked to differentiation and maturation of embryonic stem cells to immune cells, adipocytes, chondrocytes, myocytes, and lung tissues.(Blanc and Richard, 2017; Yang and Bedford, 2013) The requirement of CARM1 is implicated in multiple cancers with its methyltransferase activity particularly addicted by hematopoietic malignancies and metastatic breast cancer.(Drew et al., 2017; Greenblatt et al., 2018; Nakayama et al., 2018; Wang et al., 2014) Our prior efforts using *in vivo* mouse and *in vitro* cell models uncovered the role of CARM1 in promoting breast cancer metastasis.(Wang et al., 2014) Mechanistically, CARM1 methylates Arg1064 of BAF155 and thus facilitates the recruitment of the BAF155-containing SWI/SNF complex to a specific subset of gene loci essential for breast cancer metastasis. CARM1 thus emerges as a novel anti-cancer target.(Wang et al., 2014)

While this cancer relevance inspired the development of CARM1 inhibitors,(Kaniskan et al., 2018; Scheer et al., 2019) many small-molecule CARM1 inhibitors lack target selectivity or cellular activity (Kaniskan et al., 2018)---two essential criteria of chemical probes.(Frye, 2010) To the best of our knowledge, EZM2302,(Drew et al., 2017; Greenblatt et al., 2018) TP-064(Nakayama et al., 2018) and **SKI-73** (www.thesgc.org/chemical-probes/SKI-73) are the only selective and cell-active CARM1 chemical probes, which were developed by Epizyme, Takeda/SGC(Structural Genomic Consortium), and our team, respectively. EZM2302 and TP-064 were developed through conventional small-molecule scaffolds occupying the substrate-binding pocket of CARM1.(Drew et al., 2017; Greenblatt et al., 2018; Nakayama et al., 2018) The potential utility of EZM2302 and TP-064 is implicated by their selective anti-proliferative effects on hematopoietic cancer cells, in particular multiple myeloma cells.(Drew et al., 2017; Greenblatt et al., 2018; Nakayama et al., 2018) However, definitive molecular mechanisms of the CARM1 addiction in these contexts remain elusive.(Greenblatt et al., 2018)

Here we report the characterization and novel utility of **SKI-73**---a chemical probe of CARM1 with pro-drug properties. **SKI-73** can readily penetrate cell membranes and then be processed into two active CARM1 inhibitors containing 6′-homosinefungin (**HSF**) as their core scaffold.(Scheer et al., 2019; Wu et al., 2016) Notably, the two inhibitors can be accumulated inside cells at remarkable high concentrations and for a prolonged period. The potency, selectivity, modes of action, on-target engagement, and off-target effects of these compounds were characterized with multiple orthogonal assays *in vitro* and under cellular settings. The pharmacological inhibition of CARM1 by **SKI-73** recapitulates the anti-invasion effect of the genetic perturbation of CARM1. In the context of cellular heterogeneity, we developed a cell-cycle-aware algorithm for single-cell RNA-seq (scRNA-seq) analysis and dissected the invasion-prone subset of breast cancer cells that is sensitive to **SKI-73** treatment. Our scRNA-seq analysis provides the unprecedented insight that pharmacological inhibition of CARM1 alters epigenetic plasticity and suppresses invasion by suppressing the most invasive subpopulation of breast cancer cells.

## Results

### Development of 6′-homosinefungin derivatives as potent and selective CARM1 inhibitors

Upon developing cofactor-competitive PMT inhibitors,(Wu et al., 2016; Zheng et al., 2012) we identified 6′-homosinefungin (**HSF**, **1**) for its general high affinity to Type I PRMTs (Fig. 1a,b, S1). As a SAM mimic, **1** binds to the Type I PRMTs---PRMT1, CARM1, PRMT6 and PRMT8--- with IC_50_ of 13~300 nM (Fig. 1a,c, Table S1). Its relative affinity to Type I PRMTs aligns with that of the SAM mimics **SAH** and **SNF** (around 20-fold lower IC_50_ of **1** versus **SAH** and **SNF**, Fig. 1a,c, Table S1). This observation argues that **1** retains the structural features of **SAH** and **SNF** to engage PRMTs and meanwhile leverages its 6′-methyleneamine group for additional interaction. Strikingly, the **HSF** derivative **2a**, which was synthesized *via* the same precursor **3** (Fig. 1b, S1), preferentially binds to CARM1 with IC_50_ = 30 ± 3 nM and > 10-fold selectivity over other 7 human PRMTs and 26 methyltransferases of other classes (Fig. 1c, Table S1). The structural difference between **2a** and **1** (Fig. 1b) suggests that the *N*-benzyl substituent enables **2a** to engage CARM1 *via* a distinct mechanism (see results below). This engagement is expected to be maintained by **5a**, an amide derivative of **2a** prepared from the common precursor **3** and then the intermediate **4** (Fig. 1b, S2). Here **5a** shows an IC_50_ of 43 ± 7 nM against CARM1 and a >10-fold selectivity over the panel of 33 diverse methyltransferases (Fig. 1c, Table S1). In comparison, the negative control compounds **2b** (Bn-SNF)(Zheng et al., 2012) and **5b** (Figure 1b, S3), which differ from **2a** and **5a** only by the 6′-methylene group, poorly inhibit CARM1 (IC_50_ > 25 μM and 1.91 ± 0.03 μM) (Fig. 1c, Table S1). The dramatic increase of the potency of **2a** and **5a** in contrast to **2b** and **5b** supports an essential role of the 6′-methylene moiety on binding CARM1. Distinguished from SAM mimics **SAH**, **SNF** and **1** as nonspecific PMT inhibitors, **2a** and **5a** were developed as potent and selective SAM analog inhibitors of CARM1 (Fig. 1c, Table S1).

**Figure 1.**
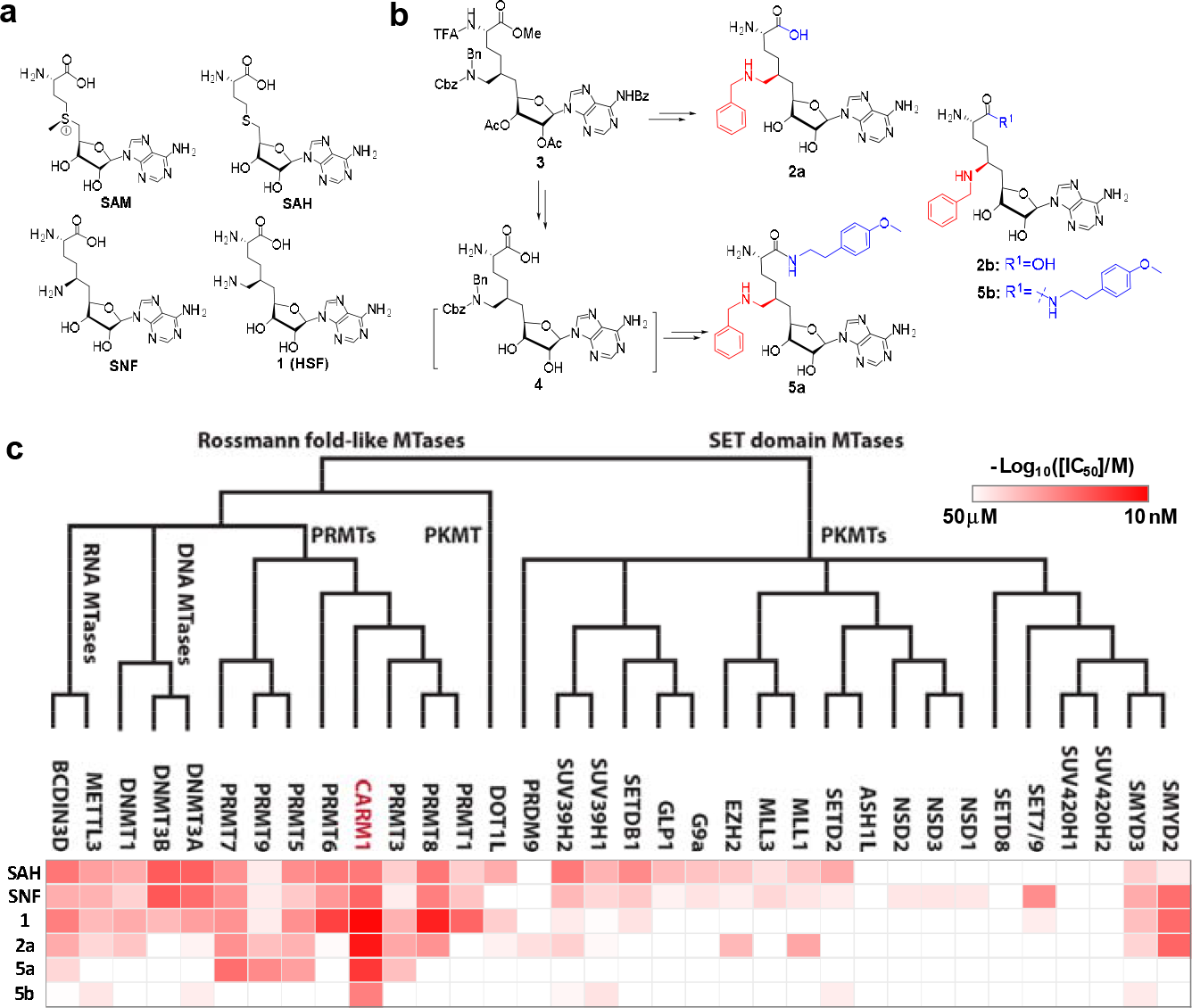
Structures, synthesis and target inhibition of SAM analogs. **a**, Structures of SAM, SAH, sinefungin (**SNF**) and 6′-homosinefungin (**HSF**, **1**). **b**, Structures and synthetic outline of **HSF** derivatives **2a** and **5a**, and their structurally-related control compounds **2b** and **5b**. **c**, IC_50_ heat-map of SAM analogs against 34 methyltransferases. **HSF** derivatives **2a** and **5a** were identified as potent and selective inhibitors of CARM1; **2b** and **5b** as their respective control compounds.

### Modes of interaction of 6′-homosinefungin derivatives as CARM1 inhibitors

With **2a** and **5a** characterized as CARM1 inhibitors, we leveraged orthogonal *in vitro* assays to explore their modes of interaction (Fig. 2a). CARM1 inhibition by **2a** and **5a** was assessed in the presence of various concentrations of SAM cofactor and H3 peptide substrate (Fig. 2b,c). IC_50_ values of **2a** and **5a** showed a linear positive correlation with SAM concentrations, as expected for SAM-competitive inhibitors.(Daigle et al., 2011; Luo, 2018; Zheng et al., 2012) The *K*_d_ values of **2a** and **5a** (*K*_d,**2a**_ = 17 ± 8 nM; *K*_d,**5a**_ = 9 ± 5 nM) were extrapolated from the y-axis intercepts upon fitting the equation IC_50_ = [SAM]×*K*_d_/*K*_m,SAM_ +*K*_d_ (Fig. 2b).(Segel, 1993) *K*_m,SAM_ of 0.21 ± 0.09 μM and 0.28 ± 0.14 μM (an averaged *K*_m,SAM_ = 0.25 μM) for competition with **2a** and **5a** can also be derived through the ratio of the y-axis intercepts to the slopes (Fig. 2b and Supplementary Methods).(Segel, 1993) In contrast, the presence of the H3 peptide substrate had negligible effect on the binding of **2a** and **5a**, indicating their substrate-noncompetitive character (Fig. 2c). The SAM analogs **2a** and **5a** were thus characterized as SAM-competitive, substrate-noncompetitive inhibitors of CARM1.

**Figure 2.**
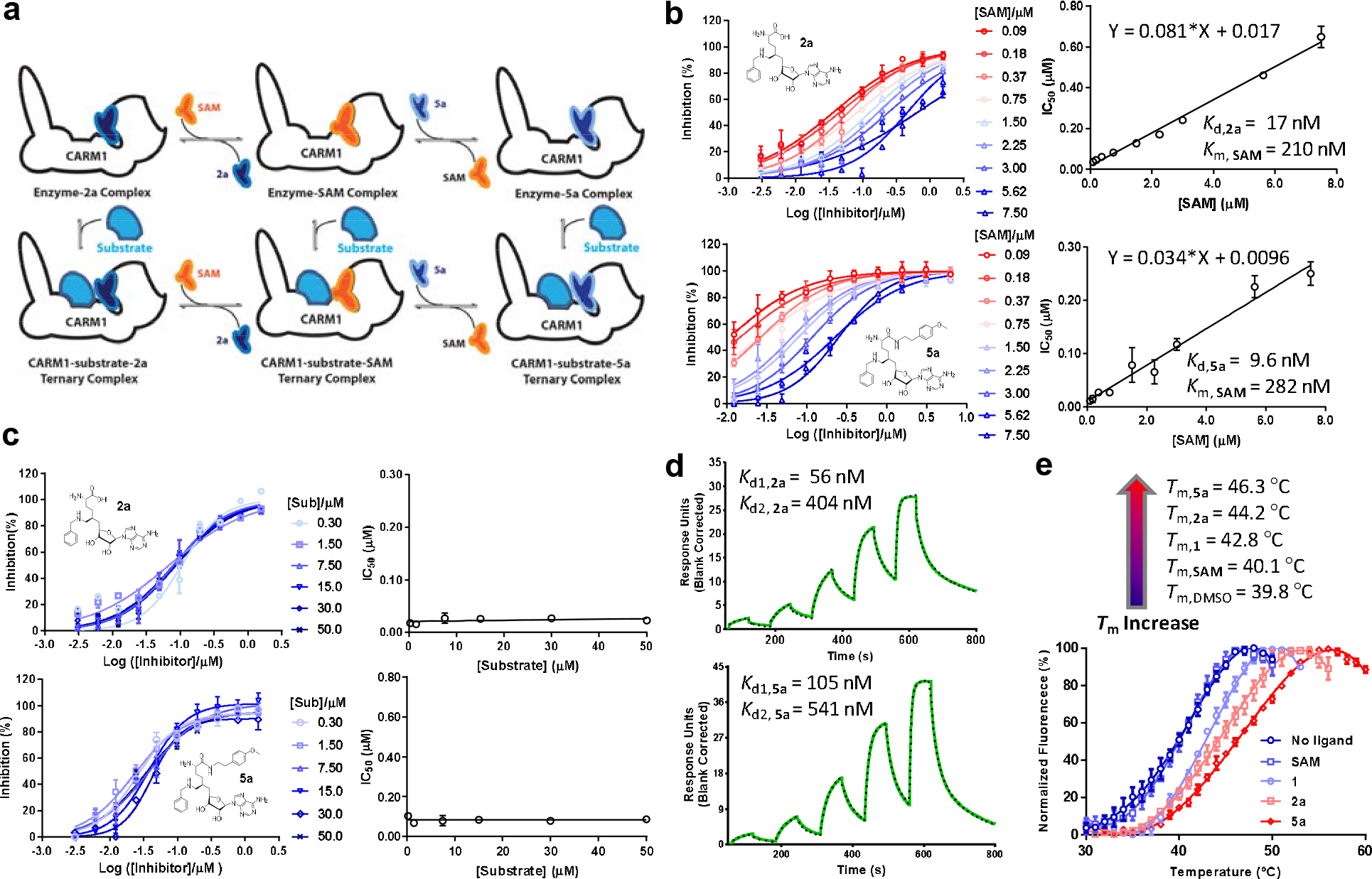
*In vitro* characterization of CARM1 inhibitors 2a and 5a. **a**, Schematic description of CARM1 in complex with SAM, **2a** and **5a** in the absence or presence of a substrate peptide. **b,c**, IC_50_ of **2a** and **5a** in the presence of varied concentrations of SAM and H3 peptide substrate. IC_50_ data were obtained and presented as the mean of replicates ± standard errors. The IC_50_ values of **2a** and **5a** show a linear increase versus the SAM concentration but remain near constant versus the substrate concentration. Given the SAM competitive character, the *K*_d_ values of **2a** and **5a** as well as *K*_d,SAM_ can be obtained according to IC_50_ = [SAM]×*K*_d_/*K*_d,SAM_ +*K*_d._ **d**, SPR assay for the binding of CARM1 by **2a** and **5a**. Processed sensorgrams upon ligand binding (black dots) were fit with the kinetics of a biphasic binding mode (green line) with *K*_d1,**2a**_ = 56 nM (0.06 ± 0.02 μM) and *K*_d2,**2a**_ = 404 nM (0.4 ± 0.1 μM); *K*_d1,**5a**_ = 105 nM (0.10 ± 0.01 μM) and *K*_d2,**5a**_ = 541 nM (0.54 ± 0.07). **e**, Thermal shift assay of CARM1 in the absence or presence of SAM, **1**, **2a**, and **5a**. *T*_m_ values of 39.8 ± 0.2 °C, 40.1 ± 0.5 °C, 42.8 ± 0.3 °C, 44.2 ± 0.6 °C and 46.3 ± 0.3 °C (means of triplicates ± standard derivatives) were obtained for apo-CARM1 and CARM1 complex with 5 μM SAM, **1**, **2a**, and **5a**, respectively.

The CARM1-binding kinetics of **2a** and **5a** were also examined using surface plasmon resonance (SPR) (Fig. 2d). The SPR signal progression of **2a** and **5a** fits with a biphasic rather mono-phasic binding mode with the lower *K*_d1,**2a**_ = 0.06 ± 0.02 μM, *K*_d1,**5a**_ = 0.10 ± 0.01 μM, and the higher *K*_d2,**5b**_ = 0.54 ± 0.07 μM, *K*_d2,**2a**_ = 0.4 ± 0.1 μM, likely due to multi-phase binding kinetics of **2a** and **5a** (Figure 2d). *In vitro* thermal shift assay(Blum et al., 2014) further showed that the binding of **2a** and **5a** increases the melting temperature (*T*_m_) of CARM1 by 4.4 °C and 6.5 °C, respectively (Fig. 2e, *T*_m,**2a**_ = 44.2 ± 0.4 °C and *T*_m,**5a**_ = 46.3 ± 0.3 °C versus *T*_m,DMSO_ = 39.8 ± 0.3 °C as control). In contrast, the binding of SAM and **1** shows much less effects on *T*_m_ of CARM1 (Fig. 2e*, T*_m,SAM_ = 40.1 ± 0.3 °C and *T*_m,**1**_ = 42.8 ± 0.4 °C versus *T*_m,DMSO_ = 39.8 ± 0.3 °C). Therefore, albeit comparable affinity of **1**, **2a** and **5a** to CARM1 (IC_50_ = 13~43 nM, Fig. 1c), their well-separated effects on *T*_m_ suggest that these inhibitors engage CARM1 differentially (see results below). Multiple orthogonal biochemical assays thus verified tight binding of **2a** and **5a** with CARM1.

### Structural rationale of 6′-homosinefungin derivatives as CARM1 inhibitors

To further seek structural rationale of **5a** and **2a** for CARM1 inhibition, we solved the X-ray structure of CARM1 in complex with **5a** and modeled the CARM1 binding of **2a** (Fig. 3, Supplementary Results and Methods). The overall topology of the CARM1-**5a** complex is indistinguishable with a V-shape subunit of CARM1 dimer in complex with **SNF** and **1** (Figure S4, S5 and Table S2-4)---the Rossmann fold of Class I methyltransferases (Fig. 3a).(Luo, 2018) However, **5a** adopts a noncanonical pose with its 6′-*N*-benzyl moiety in a binding pocket that used to be occupied by the α-amino carboxylate moiety of canonical ligands such as **SAH**, **SNF** and **1** (Fig. 3b, S4 and Table S2-4), while the α-amino methoxyphenethyl amide moiety of **5a** protrudes into the substrate-binding pocket.(Boriack-Sjodin et al., 2016; Sack et al., 2011) This noncanonical mode is consistent with the SAM-competitive character of **5a** (Fig. 2b). Under the canonical setting, the guanidinium moiety of Arg168 forms a salt bridge with the carboxylic moiety of canonical ligands (Fig. 3c). In contrast, Arg168 in the CARM1-**5a** complex has to adopt an alternative orientation (two possible configurations), accompanied by an altered conformation of Glu257, to accommodate the 6′-*N*-benzyl moiety of **5a** (Fig. 3c). The α-amino amide moiety of **5a** also engages CARM1 through the combined outcomes of a hydrogen-bond network with Glu266 and His414 and hydrophobic interactions with Phe152 and Tyr261 (Fig. 3d). Interestingly, the overlaid structures of CARM1 in complex with **5a** and a substrate peptide implicate a steric clash and thus a potential competitive-binding mode between **5a** and a CARM1 substrate (Fig. 3e). However, the apparent substrate-noncompetitive character of **5a** (Figure 2c) suggests that this steric clash might be avoided if there is no significant energy penalty for the substrate Arg to adopt alternative conformation(s).

**Figure 3.**
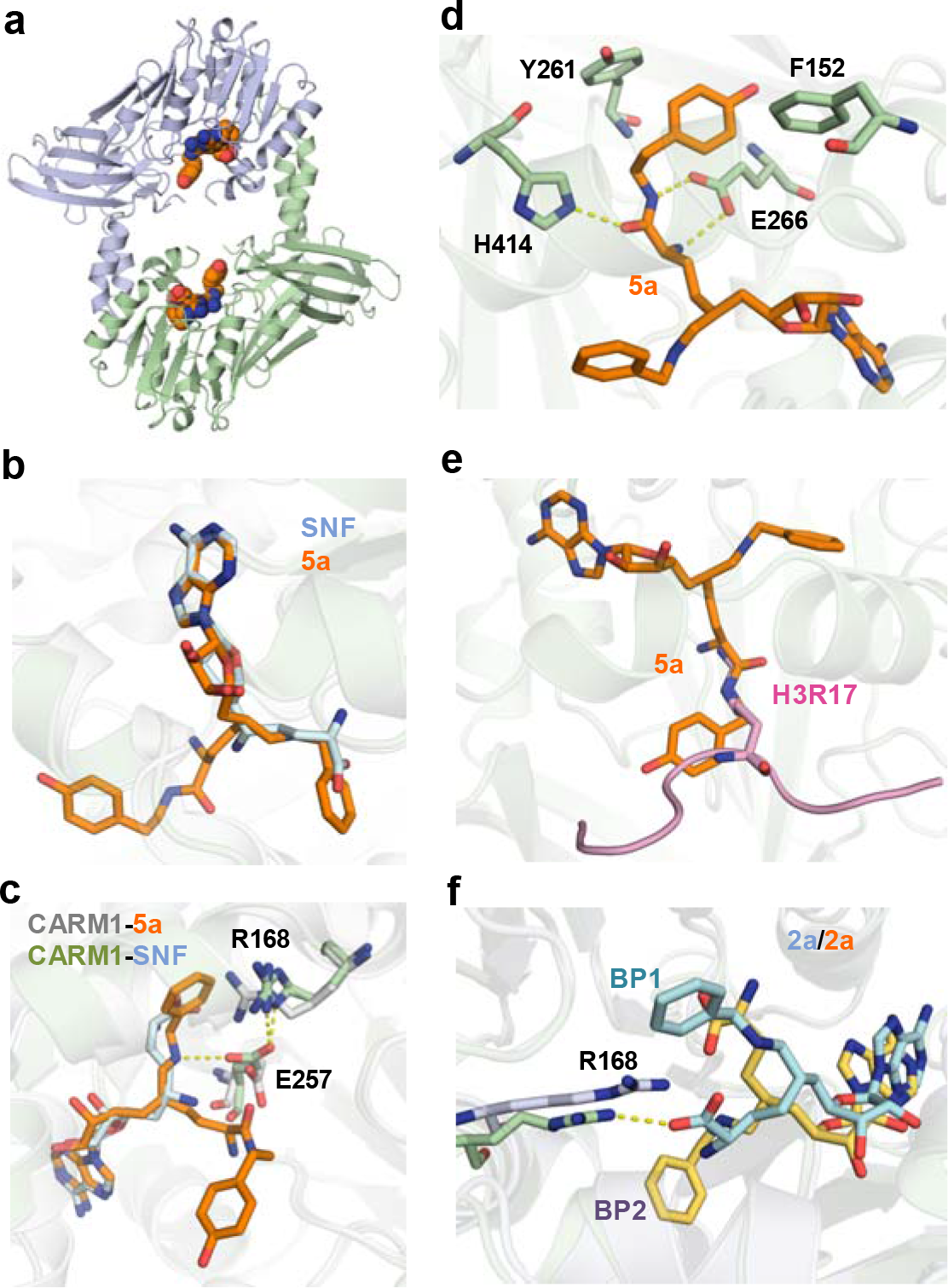
Crystal structure or molecular modeling of CARM1 in complex with 5a and 2a. **a**, Overview of the Rossmann fold in the X-ray structure of CARM1 with **5a**. **b**, Comparison of the binding modes between **5a** (noncanonical) and **SNF** (canonical). The structure of **SNF** was extracted from a CARM1-**SNF**–**H3R17** complex (PDB 5DX0). **c**, Key interactions between CARM1 and ligands in canonical and noncanonical binding modes. The differentiated interactions are highlighted in grey (CARM1) and blue (**SNF**) for the canonical mode; green (CARM1) and orange (**5a**) for the noncanonical mode. **d**, Additional interactions via the α mino amide moiety of **5a. e**, Steric clash between the α amino amide moiety of **5a** and an Arg substrate. The structure of the Arg substrate was extracted from a CARM1**-SNF-H3R17** complex (PDB 5DX0). **f**. Two modeled binding poses (**BP1** and **BP2**) of **2a** upon binding CARM1.

The binding mode of the CARM1-**2a** complex was modeled via molecular docking followed by molecular dynamics (MD) simulation (Supplementary Methods). Here we uncovered two distinct poses of **2a** (Binding Pose 1/2 or **BP1/2**) with the C4'-C5'-C6'-C7' dihedral angle of −50° and −170°, respectively (Fig. 3f). **BP1** was characterized by the direct interaction between the α-amino carboxylate moiety of **2a** with the guanidinium of Arg168, while **BP2** features a titled orientation of Arg168 to accommodate the 6′-*N*-benzyl moiety of **2a** (the Cβ-Cγ-Cδ-Nε dihedral angle χ3 = 180° for **BP1** versus χ3 = −65° of **BP2**) (Fig. 3f, S7). The BP1 and BP2 of **2a** closely resemble those of **1** and **5a**, respectively, in terms of the orientations of Arg168 and the α-amino carboxylate moiety of ligands. When the same modeling protocol was applied to the CARM1-**SNF** complex, only the canonical pose was identified (Fig. S7). Energy calculation indicated that both **BP1** and **BP2** are stable with comparable binding free energies. Interestingly, the side chain configurations of His414 in both **BP1** and **BP2** (the C-Cα-Cβ-Cγ dihedral angle χ_1_ = −48° and −66°) are different from those in the CARM1-**5a** complex and the CARM1-**SNF** complex (the C-Cα-Cβ-Cγ dihedral angle χ_1_ = 81°) (Fig. S7). Collectively, **5a** and **2a**, though structurally related to the SAM analogs **1** and **SNF**, engage CARM1 via distinct modes of interaction.

### A pro-drug-like 6′-homosinefungin derivative as a cell-active CARM1 inhibitor

While the *in vitro* characterization demonstrated the potency and selectivity of **2a** and **5a** against CARM1, we anticipated their poor membrane permeability as observed for structurally-related analogs such as **SAH** and **SNF** (Fig. 1a).(Boriack-Sjodin et al., 2016; Sack et al., 2011) The lack of membrane penetration is likely due to their primary amine moiety, which has pKa of ~ 10 and is fully protonated at a physiological pH of 7.4. Given the essential roles of the 9′-amine moiety of **2a** and **5a** in CARM1 binding (Fig. 3d), we envisioned overcoming the membrane permeability issue via a pro-drug strategy by cloaking this amine moiety with a redox-triggered trimethyl-locked quinone propionate moiety (**TML**, Fig. 4a).(Levine and Raines, 2012) We thus prepared **6a** as well as its control compound **6b** by derivatizing **5a** and **5b** with the **TML** moiety (Fig. S2, S3). To assess the cellular activity of **6a**, we relied on our prior knowledge that CARM1 methylates the Arg1064 of BAF155, a core component of the SWI/SNF chromatin remodeling complex, and CARM1 knockout abolishes this posttranslational modification in MCF-7 cells.(Wang et al., 2014) Treatment of MCF-7 cells with 10 μM of **6a** fully suppressed this methylation mark, whereas treatment with **2a** and **5a** did not affect this mark (Fig. 4b). We thus demonstrated the prodrug-like cellular activity of **6a**.

**Figure 4.**
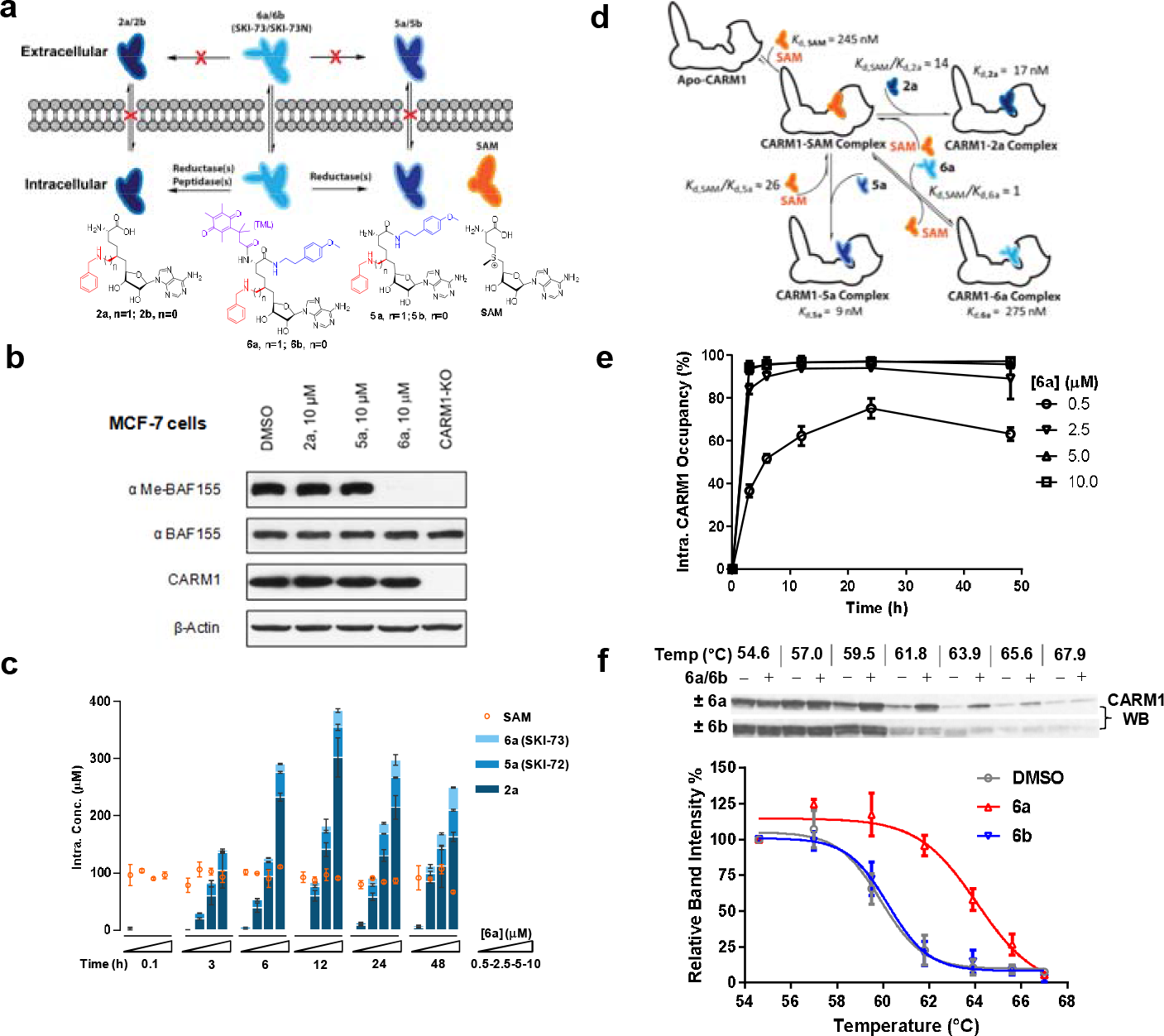
Characterization of cellular activity of 6a as a chemical probe. **a**, Schematic description of extracellular and intracellular fates of **2a**, **5a** and **6a**. Extracellularly, **2a**, **5a** and **6a** are stable; only **6a** can readily penetrate cell membrane. Intracellularly, **6a** can be processed into **5a** and **2a**. Given the poor membrane permeability of **2a** and **5a**, they are accumulated within cells at high concentrations. **b**, CARM1 inhibition of **2a**, **5a** and **6a** in MCF-7 cells with BAF155 methylation as a mark. MCF-7 cells were treated with 10 μM of **2a**, **5a** and **6a** for 48 hours. The ratios between me-BAF155 and BAF155 were quantified as a cellular reporter of CARM1 inhibition. DMSO-treated MCF-7 cells and MCF-7 CARM1-*KO* cells were used as negative and positive controls, respectively. **c**, MS-based quantification of intracellular concentrations of **2a**, **5a**, **6a** and SAM. These compounds were accumulated within MDA-MB-231 cells in a dose- and time-dependent manner. In comparison, the intracellular concentration of SAM remains a constant of 89 ± 16 μM. **d**, Schematic description of intracellular engagement of CARM1 by **6a**, **5a** and **2a** in the presence of the SAM cofactor (eqs. S5-7). **e**, Modeled ligand occupancy of CARM1 with **2a**, **5a** and **6a** as ligands in competition with the SAM cofactor. Percentage of competitive CARM1 occupancy was calculated on the basis of the concentrations of ligands (SAM, **2a**, **5a** and **6a**, Fig. 4c) and their *K*_d_ values (Fig. 2c). **f**, Cellular thermal shift assay (CETSA) of CARM1. Representative western blots of CARM1 in MDA-MB-231 cells upon the treatment of 15 μM **6a** or its negative control **6b** for 48 hours with DMSO treatment as reference. The relative intensity of CARM1 was quantified. The *T*_m_ values were determined at the 50% loss of the relative intensity signals with *T*_m,**6a**_ = 63.9 ± 0.3 °C, *T*_m,**6b**_ = 60.2 ± 0.6 °C and *T*_m,DMSO_ = 59.6 ± 0.2 °C.

### Characterization of 6a (SKI-73) as a chemical probe of CARM1

To further evaluate **6a** as a chemical probe of CARM1, we assessed the efficiency of **6a** to suppress CARM1-dependent invasion of breast cancer cells. Because of the pro-drug character of **6a** and its control compound **6b**, we first developed quantitative LC-MS/MS methods to examine their cellular fates (Supplementary Methods). Upon the treatment of MDA-MB-231 cells with **6a**, we observed its time- and dose-dependent intracellular accumulation (Fig. 4c). While we anticipated the conversion of the pro-drug **6a** into **5a**, a striking finding is that **6a** can also be readily processed into **2a** inside cells (Fig. 4c). Remarkably, > 100 μM of **2a** can be accumulated inside cells for 2 days after 6-h treatment with a single dose of 5~10 μM **6a**. This observation likely reflects a slow efflux and thus effective intracellular retention of **2a** due to its polar α-amino acid zwitterion moiety. Given that cellular CARM1 inhibition is involved with multiple species (**2a**, **5a** and **6a**) in competition with SAM, we modeled the ligand occupancy of cellular CARM1 on the basis of their *K*_d_ values (*K*_d,**2a**_ =17 nM, *K*_d,**5a**_ =9 nM, *K*_d,**6a**_ =0.28 μM and *K*_d,SAM_ ≈*K*_m,SAM_=0.25 μM) and MS-quantified intracellular concentrations (Fig. 4d, eqs. S5-S7, Supplementary Methods). The SAM cofactor, whose intracellular concentration was determined to be 89 ± 16 μM (Fig. 4c,d), is expected to occupy > 99.5% CARM1 with residual < 0.5% as the apo-enzyme under a native setting. With single doses of **6a** of 2.5~10 μM, the combined CARM1 occupancy by **2a**, **5a** and their pro-drug precursor **6a** rapidly reached the plateaus of >95% within 6 h, and was maintained at this level for at least 48 h (Fig. 4e). Notably, the treatment of **6a** as low as 0.5 μM is sufficient to reach 60% target engagement within 10 h and maintain this occupancy for 48 h (Fig. 4e). The time- and dose-dependent progression of the CARM1 occupancy by these ligands thus provides quantitative guidance upon the treatment of MDA-MB-231 cells with **6a**.

With a cellular thermal shift assay (CETSA),(Jafari et al., 2014) we further observed that the treatment of MDA-MB-231 cells with **6a** but not the control compound **6b** increases cellular *T*_m_ and thus thermal stability of CARM1 by 4.3 ± 0.6 °C (Fig. 4f). The distinct effect of **6a** in contrast to **6b** on the cellular *T*_m_ of CARM1 aligns well with the 4.1~6.2 °C difference of *in vitro T*_m_ of CARM1 upon binding **2a** and **5a** versus SAM (Figs. 2e). Here **6b** can penetrate cell membrane and be processed into **5b** and **2b** in a similar manner as **6a** (Figure S11). These observations thus present the cellular evidence of CARM1 engagement of **2a** and **5a**.

To further characterize **6a** as a CARM1 chemical probe, we treated MDA-MB-231 cells with **6a** and examined the Arg1064 methylation of BAF155 and the Arg455/Arg460 methylation of PABP1, two well-characterized cellular methylation marks of CARM1.(Lee and Bedford, 2002; Wang et al., 2014) These methylation marks can be fully suppressed by **6a** in a dose-dependent manner (Fig. 5a). The resultant EC_50_ values of 0.45~0.75 μM (Fig. 5b) are well correlated with the modeled 60% cellular occupancy of CARM1 upon the treatment of 0.5 μM **6a** for 48 h (Fig. 4e). In contrast, the treatment of the negative control compound **6b** showed no effect on these methylation marks (Fig. 5a). We therefore demonstrated the robust use of **6a** (**SKI-73**) as a CARM1 chemical probe and **6b** (**SKI-73N**) as its control compound.

**Figure 5.**
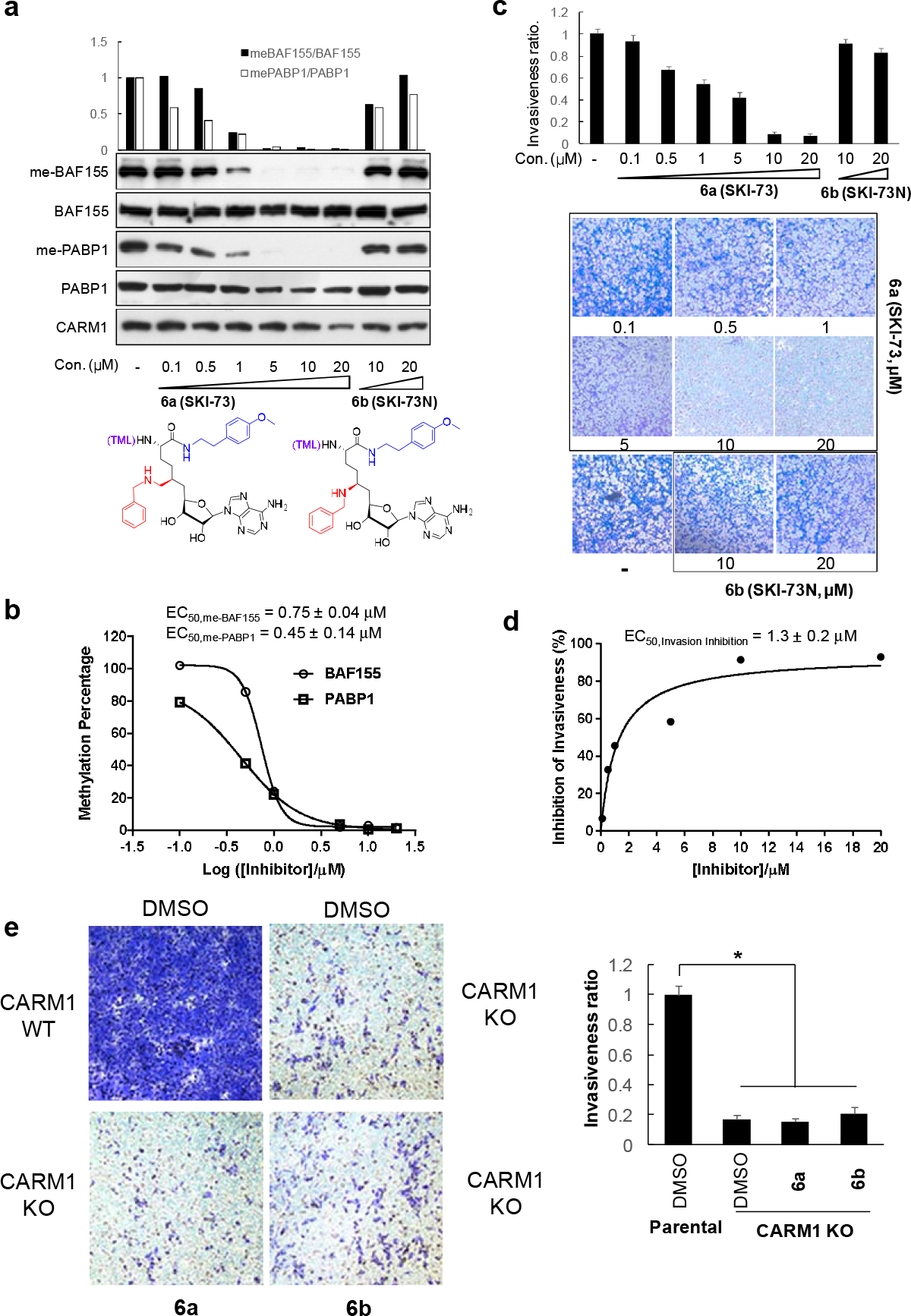
Biological outcomes of CARM1 inhibition by 6a in MDA-MB-231 cells. **a**, Dose-dependent depletion of BAF155 methylation and PABP1 methylation by **6a**. BAF155 methylation and PABP1 methylation, two marks of the CARM1-specific methyltransferase activity, were examined upon the treatment of **6a** and its structural analog **6b** (negative control compound) for 48 hours. Western Blot analysis was then conducted to quantify the relative intensities of the methylated versus total proteins (BAF155 and PABP1, two replicates with representative one shown). **b**, EC_50_ of the methylation depletion of BAF155 and PABP1. The relative intensity of the methylated versus total BAF155 or PABP1 was plotted against log[**6a**] with the resultant EC_50_ upon fitting a standard sigmoid curve using GraphPad Prism. **c**, Inhibition of cell invasion by **6a**. Representative images of trans-well migration of MDA-MB-231 cells were shown upon treatment with various concentrations of **6a** (**SKI-73**) or its control compound **6b** (**SKI-73N**) for 16 hours. Invasive cells were fixed and stained with crystal violet. The invasiveness ratios were determined using the relative cell invasion of the treatment of **6a** or **6b** versus DMSO treatment. **d**, EC_50_ of invasion inhibition by **6a**. The invasiveness ratios were plotted as a function of the concentration of **6a**. EC_50_ of 1.3 ± 0.2 μM was obtained upon fitting a standard sigmoid curve using GraphPad Prism. **e**, Effect of **6a** on cell invasion in combination with CARM1 *KO*. Representative images of trans-well migration of parent and CARM1 *KO* MDA-MB-231cells shown upon treatment with DMSO, **6a** or **6b** for 16 hours. The results were analyzed in a similar manner as described for Figs. 5c,d.

### Inhibition of *in vitro* invasion but not proliferation of breast cancer cells by SKI-73

After demonstrating **6a** (**SKI-73**) as a chemical probe of CARM1, we examined whether chemical inhibition of CARM1 can recapitulate biological outcomes associated with CARM1 knockout (CARM1-*KO*).(Wang et al., 2014) Our prior work showed that CARM1’s methyltransferase activity is required for invasion of MDA-MB-231 cells.(Wang et al., 2014) We thus conducted a matrigel invasion assay with MDA-MB-231 cells in the presence of **6a**. Relative to the control treatment with DMSO, the treatment of **6a** (**SKI-73**) but not its negative control compound **6b** (**SKI-73N**) suppressed the invasion of MDA-MB-231 cells in a dose-dependent manner (EC_50_ = 1.3 μM) (Fig. 5c,d). The treatment with ≥ 10 μM **6a** reached the maximal 80% suppression on the invasion of MDA-MB-231 relative to the DMSO control, which is comparable with the phenotype of CARM1-*KO* (Fig. 5e). Critically, no further inhibition by **6a** on the invasiveness was observed upon its treatment of MDA-MB-231 CARM1-*KO* cells in comparison with the treatment with DMSO or **6b** (Fig. 5e). Notably, the treatment with **6a** and **6b** under the current condition has no apparent impact on the proliferation of parental and CARM1-*KO* MDA-MB-231 cells (Figure S12), consistent with the intact proliferation upon the treatment with other CARM1 chemical probes.(Drew et al., 2017; Greenblatt et al., 2018; Nakayama et al., 2018) These results suggest that **6a** (**SKI-73**) and CARM1 knockout perturb the common, proliferation-independent biological process and then suppresses 80% of the invasiveness of MDA-MB-231 cells. We thus characterized **6a** (**SKI-73**) as a chemical probe to interrogate CARM1-dependent invasion of breast cancer cells.

### scRNA-seq and cell-cycle-aware algorithm reveals CARM1-dependent epigenetic plasticity

Because of the advancement of scRNA-seq technology, stunning subpopulation heterogeneity has been uncovered even for well-defined cellular types.(Tanay and Regev, 2017) In the context of tumor metastasis including its initial step---invasion, epigenetic plasticity is required to offer a small subset of tumor cells to adapt distinct transcriptional cues for neo-properties.(Chatterjee et al., 2017; Flavahan et al., 2017; Wu et al., 2018) To explore the feasibility of dissecting the CARM1-dependent, invasion-prone subset of MDA-MB-231 breast cancer cells, we formulated a cell-cycle-aware algorithm of scRNA-seq analysis and dissected those subpopulations sensitive to CARM1 perturbation (Figure 6a, Supplementary Methods). Here we conducted 10× Genomics droplet-based scRNA-seq of 3,232, 3,583 and 4,099 individual cells (the total of 10,914 cells) exposed to 48-hour treatment with **SKI-73** (**6a**), **SKI-73N** (**6b**) and DMSO, respectively. Guided by Silhouette analysis of each treatment condition as well as their combination for the modularity-based shared-nearest-neighbor(SNN) graph clustering, cell-cycle-associated transcripts were identified as dominant signatures to define subpopulations (Figure S13-30). These signatures naturally exist for proliferative cells and are not expected to be specific for the invasive phenotype. To dissect subpopulation-associated transcriptomic signatures of invasive cells, we included one additional layer for hierarchical clustering by first classifying the individual 10,914 cells into G0/G1, S, and G2/M stages (6,885, 1,520 and 2,509 cells, respectively) (Figure S18, Table S5), and then conducted the unsupervised clustering within each cell-cycle-aware subset (Figure S18, S31-38, S42-45, Table S5). To resolve efficiently the subpopulations associated with the three treatment conditions (**6a**, **6b** and DMSO) without redundant clustering, we developed an entropy analysis method and relied on the Fisher Exact test (Supplementary Methods). The optimal scores of the combined methods were implemented for the modularity-based SNN graph clustering and to determine the numbers of cluster for each subset (Figure S32, S36, S43).(Butler et al., 2018) The cell-cycle-aware algorithm allowed the clustering of the 10,914 cancer cells according to the three cell cycle stages under the three treatment conditions (**6a**, **6b** and DMSO) and resulted in 21, 7 and 6 subpopulations in G0/G1, S, and G2/M phases, respectively (Figure 6b, S33, S37, S44, Table S6-8). Notably, the 48-hour treatment with **SKI-73** (**6a**) or **SKI-73N** (**6b**) had no effect on the cell cycle, as indicated by the comparable cell-cycle distribution patterns between **SKI-73**, **SKI-73N**, and DMSO treatment (Figure S18, Table S5). This result is also consistent with the intact proliferation upon the treatment with **SKI-73** and **SKI-73N** (Figure S12).

**Figure 6.**
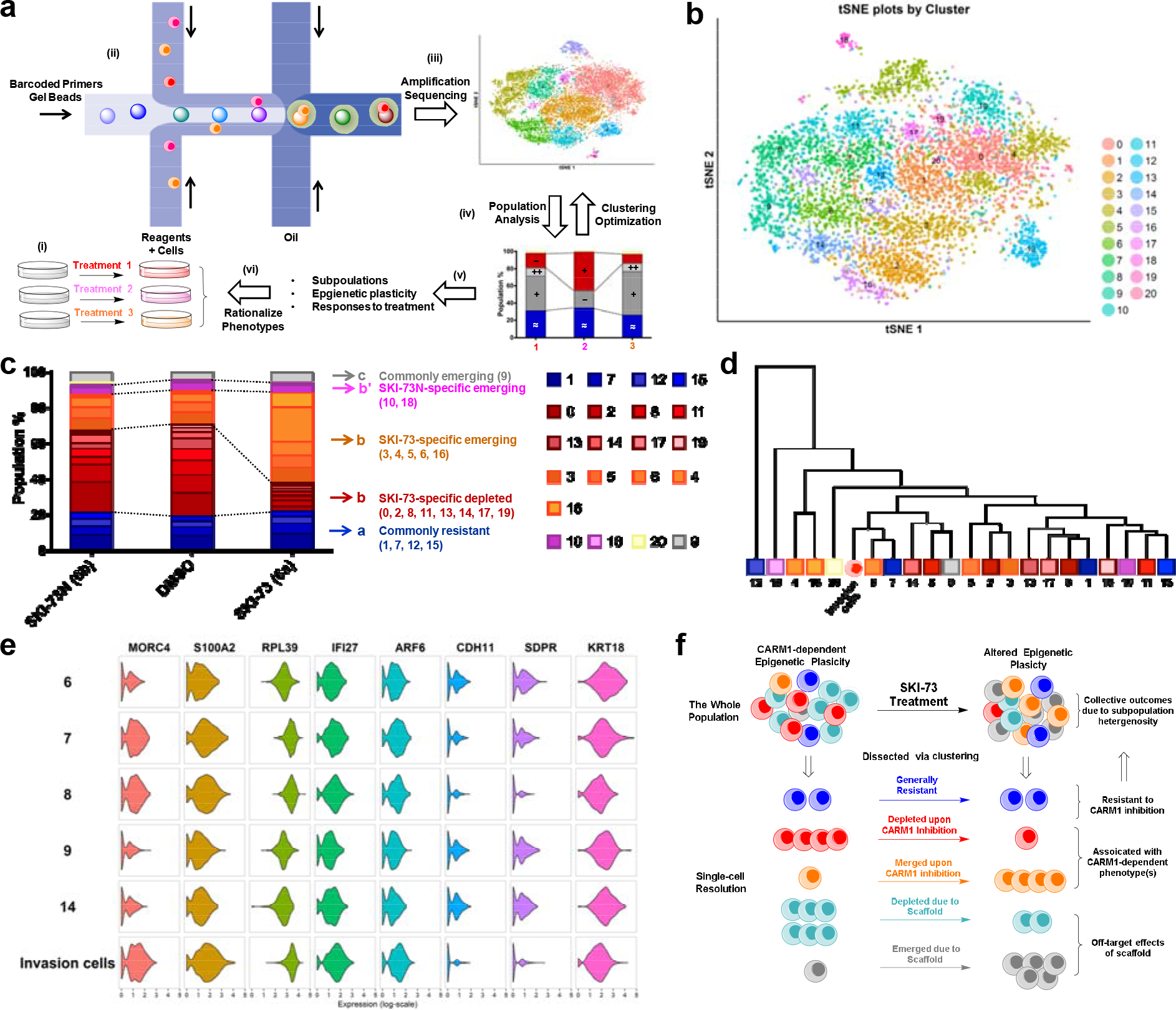
scRNA-seq analysis of MDA-MB-231 cells upon CARM1 perturbation. (a) Schematic description of scRNA-seq analysis algorithms. (b) tSNE plots of the 21 clustered subpopulations of the G0/G1-phase cells treated with **SKI-73N**, DMSO and **SKI-73**. (c) Population analysis of the 21 clustered subpopulations of the G0/G1-phase cells upon the treatment with **SKI-73N** and **SKI-73**. (d) Phylogenic tree of the 21 clustered subpopulations of the G0/G1-phase cells and invasion cells. (e) Violin plots of the representative transcripts of G0/G1-phase invasion cells that distinguish Subpopulation-8 from the closely related subpopulations Subpopulation-6, 7, 9, 14 of G0/G1-phase cells.

### CARM1-associated epigenetic plasticity of breast cancer cells with single-cell resolution

With the 21, 7 and 6 subpopulations clustered into the G0/G1, S, and G2/M stages, respectively, we then conducted population analysis between the three treatment conditions (**SKI-73** and **SKI-73N** versus DMSO) (Figure 6c, S39, S46 and Table S6-8). These subpopulations can be readily classified into five distinct categories according to how the cells respond to **SKI-73** and **SKI-73N** treatment in each cell cycle stage: commonly resistant/emerging/depleted versus differentially depleted/emerging (**SKI-73/SKI-73N-**specific) (Figure 6c, S39, S46 and Table S6-8). Here we are particularly interested in the **SKI-73**-specific depleted subpopulations (0/2/8/11/13/14/17/19 of G0/G1-phase cells and 3 of S-phase cells) as the potential invasion-associated subpopulations, given their sensitivity to **SKI-73** but not its control compound **SKI-73N**. The subpopulations that remain unchanged after the treatment of **SKI-73** and **SKI-73N** (1/7/12/15 of G0/G1-phase cells; 2/4/5 of S-phase cells; 1 of G2/M-phase cells) were defined as the common resistant subset. **SKI-73**-specific emerging subpopulations (3/4/5/6/16 of G0/G1-phase cells; 6 of S-phase cells; 4 of G2/M-phase cells) are expected to be suppressed by CARM1 but emerge upon its inhibition. Other subpopulations are either associated with effects of the small-molecule scaffold of **SKI-73/SKI-73N** (commonly emerging Subpopulation-9 of G0/G1-phase cells, 0/5 of G2/M-phase cells; commonly depleted Subpopulation-2/3 of G2/M-phase cells) or **SKI-73N**-specific effects (differentially depleted Subpopulation-10/18 of G0/G1-phase cells, 0 of S-phase cells; differentially emerging Subpopulation-20 of G0/G1-phase cells). Interestingly, in comparison with **SKI-73** treatment, scRNA-seq analysis of 3,291 CARM1-*KO* cells suggests that CARM1 knockout has more profound effects on the overall landscape of the epigenetic plasticity (Figure S49). Collectively, the chemical probe **SKI-73** alters the epigenetic plasticity of MDA-MB-231 breast cancer cells via the combined effects of **SKI-73**’s molecular scaffold and specific inhibition of CARM1’s methyltransferase activity.

### Identification of CARM1-dependent, invasion-prone subpopulations of breast cancer cells

Given that **SKI-73** has no effect on cell cycle and proliferation of MDA-MB-231 cells under the current treatment dose and duration, we envision that the invasion capability of MDA-MB-231 cells mainly arises from an invasion-prone subset, 80% of which is depleted by **SKI-73** treatment (Figure 5c-e). We thus focused on Subpopulation-0/2/8/11/13/14/17/19 of G0/G1-phase cells and Subpopulation-3 of S-phase cells---in total nine depleted subpopulations specific for **SKI-73** (Figure 6c, S39, S46 and Table S6-8). To identify invasion-prone subpopulation(s) among these candidates, we compared their transcriptional signature(s) with those that freshly invaded through Matrigel within 16 hours. Strikingly, in comparison with the highly heterogenous scRNA-seq signature of the parental MDA-MB-231 cells, the freshly-harvested invasive cells (3,793 cells for scRNA-seq) are relatively homogeneous with their subpopulations mainly determined by the cell-cycle-related transcriptomic signatures (Figure S49-53). Like the cells treated with **DMSO**, **SKI-73** and **SKI-73N**, we classified the freshly-harvested invasive cells into G0/G1, S and G2/M stages (Figure S51, Table S5). Through the correlation analysis between the invasion cells and the subpopulations within each cell-cycle stage (Figure 6d, S40, S41, S47, S48, S54, S55), we readily revealed the subsets whose transcriptional signatures closely relate to those of the invasion cells including Subpopulation-6/7/8/9/14 in G0/G1-phase cells, 0/3 in S-phase cells and 1/2 of G2/M-phase cells (Table S6-11). In the context of population analysis for the nine **SKI-73**-specific depleted subpopulations, Subpopulation-8/14 of G0/G1-phase cells and Subpopulation-3 in S-phase are putative invasion-prone candidates. Subpopulation 8 of G0/G1-phase cells is the most sensitive and the only subpopulation that can be depleted by around 80% with **SKI-73** treatment (Figure 6c). Given the ~80% suppression and ~20% residual invasion capability upon **SKI-73** treatment, we argue that the invasive phenotype of MDA-MB-231 cells predominantly arises from the Subpopulation-8 of G0/G1-phase cells, which only accounts for ~8% of the parental cells in G0/G1 phase or ~5% without cell-cycle awareness. Differential expression analysis further revealed the single-cell transcriptional signatures of metastasis-implicated genes (*e.g.* MORC4, S100A2, RPL39, IFI27, ARF6, CHD11, SDPR and KRT18) that are specific for the G0/G1-phase Subpopulation-8 and invasion cells but not other G0/G1-phase invasion-prone candidates such as Subpopulation-6/7/9/14 (Fig. 6e, S55 and Table S12). The remaining cells of G0/G1-phase Subpopulation-8 after **SKI-73** treatment (Fig. 6c,d) together with others (subpopulation-6/7/9/14 in G0/G1-phase cells, 0/3 in S-phase cells and 1/2 of G2/M-phase cells, Figs. S39, S41 S46 S48) may account for the 20% residual invasion capacity. Collectively, either CARM1 knockout or CARM1 inhibition with **SKI-73** alters the epigenetic plasticity in a proliferation-independent manner, depleting the most invasion-prone subpopulation and thus suppressing the invasive phenotype.

## Discussion

### Chemical probes of CARM1

Based on a novel small-molecule scaffold 6′-homosinefungin (**HSF**), **SKI-73** was developed as a pro-drug-like chemical probe of CARM1 by cloaking the 9′-amine moiety of **5a** with the **TML** moiety. **SKI-73N** was developed as a control compound of **SKI-73**. The inhibitory activity of **SKI-73** against CARM1 was demonstrated by the ability of **SKI-73** but not **SKI-73N** to abolish the cellular methylation marks of CARM1---the Arg1064 methylation of BAF155 and the Arg455/Arg460 methylation of PABP1.(Lee and Bedford, 2002; Wang et al., 2014) While the ready intracellular cleavage of **TML** is expected for the conversion of **SKI-73** and **SKI-73N** into **5a** and **5b**, respectively, it is remarkable that **SKI-73** and **SKI-73N** can also be efficiently processed into **2a** and **2b** inside cells. Here **2a** and **5a** are presented as potent and selective CARM1 inhibitors, while their control compounds **2b** and **5b** poorly interact with CARM1. Competitive assays with SAM cofactor and peptide substrate showed that **2a** and **5a** act on CARM1 in a SAM-competitive and substrate-noncompetitive manner. The SAM-competitive mode is consistent with the ligand-complex structures of CARM1, in which the SAM binding site is occupied by **2a** and **5a**. Strikingly, as revealed by their ligand-CARM1 complex structures, **2a** and **5a** engage CARM1 via noncanonical modes with their 6′-*N*-benzyl moiety in the binding pocket that is otherwise occupied by the α-amino carboxylic moiety of the conventional SAM analogs such as **SAH**, **SNF** and **1**. This observation is consistent with the 4.1~6.5 °C increase in *in vitro* and cellular *T*_m_ of CARM1 upon binding **2a** and **5a** in contrast to the less *T*_m_ changes with SAM as a ligand. The distinct modes of interaction of CARM1 with **2a** and **5a** (Figure 3b,3f) also rationalize the CARM1 selectivity of the two SAM analogs over other methyltransferases including closely related PRMT homologs. Through mathematic modeling using the inputs of the LC-MS/MS-quantified intracellular concentrations and CARM1-binding constants of relevant **HSF** derivatives and SAM cofactor, we concluded that high intracellular concentrations of **5a** and **2a** and thus efficient CARM1 occupancy can be achieved rapidly and maintained for several days with a single low dose of **SKI-73**. The polar α-amino acid zwitterion moiety of **2a** and the polar α-amino moiety of **5a** likely account for their accumulation and long-time retention inside cells.

To the best of our knowledge, EZM2302, TP-064, **SKI-73** (www.thesgc.org/chemical-probes/SKI-73) and their derivatives are the only selective and cell-active CARM1 inhibitors.(Drew et al., 2017; Nakayama et al., 2018) While the potency, selectivity, on-target engagement and potential off-target effects associated with these compounds have been examined *in vitro* and in cellular contexts as chemical probes, EZM2302, TP-064, **SKI-73** are distinct by their molecular scaffolds and modes of interaction with CARM1 (www.thesgc.org/chemical-probes/SKI-73).(Drew et al., 2017; Nakayama et al., 2018) **SKI-73** is a cofactor analog inhibitor embedding a *N*6′-homosinefungin moiety to engage the SAM binding site of CARM1 in a cofactor-competitive, substrate-noncompetitive manner; EZM2302 and TP-064 occupy the substrate-binding pocket of CARM1 in a SAH-uncompetitive or SAM-noncompetitive manner.(Drew et al., 2017; Nakayama et al., 2018) In particular, the prodrug property of **SKI-73** allows its ready cellular uptake, followed by rapid conversion into its active forms inside cells. The prolonged intracellular CARM1 inhibition further distinguishes **SKI-73** from EZM2302 and TP-064.

### Anti-cancer effects and conventional mechanisms associated with pharmacological inhibition of CARM1

With **SKI-73** as a CARM1 chemical probe and **SKI-73N** as a control compound, we showed that pharmacological inhibition of CARM1 with **SKI-73**, but not **SKI-73N**, suppressed 80% invasion capability of MDA-MB-231 cells. In contrast, the pharmacological inhibition of CARM1 with **SKI-73** had no effect on the proliferation of MDA-MB-231 cells. This result is consistent with the lack of anti-proliferation activities of the other two CARM1 chemical probes EZM2302 and TP-064 against breast cancer cell lines.(Drew et al., 2017; Nakayama et al., 2018) The anti-invasion efficiency of **SKI-73** is in a good agreement with the intracellular occupancy and the resulting abolishment of several methylation marks of CARM1 upon the treatment of **SKI-73**. Our prior work showed that the methyltransferase activity of CARM1 is required for breast cancer metastasis.(Wang et al., 2014) Among diverse cellular substrates of CARM1,(Blanc and Richard, 2017) BAF155---a key component of the SWI/SNF chromatin-remodeling complex---is essential for invasion of MDA-MB-231 cells.(Wang et al., 2014) Mechanistically, the CARM1-mediated Arg1064 methylation of BAF155 facilitates the recruitment of the SWI/SNF chromatin-remodeling complex to a specific subset of gene loci.(Wang et al., 2014) Replacement of the native CARM1 with its catalytically dead mutant or an Arg-to-Lys point mutation at the Arg1064 methylation site of BAF155 is sufficient to abolish the invasive capability of breast cancer cells.(Wang et al., 2014) CARM1 inhibition with **SKI-73**, but not its control compound **SKI-73N**, recapitulates anti-invasion phenotype associated with the genetic perturbation of CARM1. More importantly, there is no additive effect upon combining CARM1-*KO* with **SKI-73** treatment, underlying the fact that the two orthogonal approaches target the commonly shared pathway(s) essential for invasion of breast cancer cells. In comparison to **SKI-73**, the CARM1 inhibitors EZM2302 and TP-064 demonstrated the anti-proliferation effects on hematopoietic cancer cells, in particular multiple myeloma.(Drew et al., 2017; Greenblatt et al., 2018; Nakayama et al., 2018) Mechanistically, genetic perturbation of CARM1 in the context of leukemia impairs cell-cycle progression, promotes myeloid differentiation, and ultimately induces apoptosis, likely *via* targeting pathways of proliferation and cell-cycle progression---E2F-, MYC-, and mTOR-regulated processes.(Greenblatt et al., 2018) In comparison, CARM1 inhibition with EZM2302 led to a slightly different phenotype, including reduction of RNA stability, E2F target downregulation, and induction of a p53 response signature featured for senescence.(Greenblatt et al., 2018) Collectively, the effects of CARM1 chemical probes are highly context-dependent with the different uses of **SKI-73** against invasion of breast cancer cells versus TP-064 and EZM2302 against proliferation of hematopoietic cancer cells.

### CARM1-dependent epigenetic plasticity revealed by SKI-73 with single-cell resolution

Given the increased awareness of epigenetic plasticity,(Flavahan et al., 2017) we employed the scRNA-seq approach to examine MDA-MB-231 cells and their responses to chemical and genetic perturbation with CARM1. Because of the lack of the prior reference to define subpopulations of MDA-MB-231 cells, we developed a cell-cycle-aware algorithm to cluster the subpopulations with a resolution to dissect subtle changes upon the treatment of **SKI-73** versus its control compound **SKI-73N** in each cell cycle stage. Guided by Silhouette analysis, the population entropy analysis and the Fisher Exact test, >10,000 MDA-MB-231 breast cancer cells were classified on the basis of their cell cycle stages and then clustered into 34 subpopulations. With further annotation of these subpopulations according to their different responses to the treatment of **SKI-73** versus **SKI-73N**, we readily dissected the subpopulations that were altered in a **SKI-73**-specific (CARM1-dependent) manner and then identified the subsets with the transcriptional signatures that are similar to that of the freshly-isolated invasive cells. Quantitative analysis of **SKI-73**-depleted subpopulations further revealed the most invasion-prone subpopulation, which accounts for only 5% of the total population but at least 80% invasive capability of the parental cells. Collectively, we propose a model that MDA-MB-231 cells consist of the subpopulations with their epigenetic plasticity determined by multiple factors including the CARM1-involved BAF155 methylation.(Wang et al., 2014) **SKI-73** inhibits the methyltransferase activity of CARM1, the Arg1064 methylation of BAF155, and thus the target genes associated with the methylated BAF155. These effects alter the cellular epigenetic landscape by affecting certain subpopulations of MDA-MB-231 cells without apparent effect on cell cycle and proliferation. In the context of the invasion phenotype of MDA-MB-231 cells, the subset of invasion-prone cells is significantly suppressed upon the treatment with **SKI-73**. Essential components to dissect the invasion-prone population in this CARM1-dependent epigenetic plasticity model are the scRNA-seq analysis of sufficient MDA-MB-231 cells (>10,000 cells here), the utility of the freshly isolated invasive cells as the reference, the timing and duration of treatment, and the use of **SKI-73N** and DMSO as controls. Interestingly, although the invasion-prone subpopulation is also abolished in the CARM-*KO* strain, CARM-*KO* reshapes the epigenetic plasticity in a much more profound manner---significantly reducing the subpopulation heterogeneity of MDA-MB-231 cells. The distinct outcomes between the pharmacological and genetic perturbation can be due to their different modes of action---short-term treatment with **SKI-73** versus long-term clonal expansion of CARM1-*KO* cells. The pharmacological inhibition captures the immediate response, while the genetic perturbation reports long-term and potential resistant outcomes. This work thus presents a new paradigm to understand cancer metastasis in the context of epigenetic plasticity and provides guidance to carry out similar analysis in broader contexts---other cell lines, patient derived xenograft samples, and *in vivo* mouse models of breast cancer.

## Online content

Supplementary results, methods, Figures S1-55, Tables S1-13, and references are available at https://doi.org/

## Acknowledgements

The authors thank Christina Leslie for providing suggestion of scRNA-seq analysis; the National Institutes of Health of USA (ML: R01GM096056, R01GM120570), National Cancer Institute (ML: 5P30 CA008748; WX: R01CA236356, R01CA213293), Starr Cancer Consortium (ML), MSKCC Functional Genomics Initiative (ML), the Sloan Kettering Institute (ML), Mr. William H. Goodwin and Mrs. Alice Goodwin Commonwealth Foundation for Cancer Research, and the Experimental Therapeutics Center of Memorial Sloan Kettering Cancer Center (ML), MSKCC Metastasis and Tumor Ecosystems Center (ML), the Tri-Institutional PhD Program in Chemical Biology (SC), NIH (WX), Susan G. Komen Foundation (EJK : PDF17481306), National Cancer Institute of National Institutes of Health of USA (LXQ: CA214845, CA008748), and Special Funding of Beijing Municipal Administration of Hospitals Clinical Medicine Development---YangFan Project (ZZ: ZYLX201713). The Structural Genomics Consortium is a registered charity (no. 1097737) that receives funds from AbbVie; Bayer Pharma AG; Boehringer Ingelheim; Canada Foundation for Innovation; Eshelman Institute for Innovation; Genome Canada; Innovative Medicines Initiative (EU/EFPIA) (ULTRA-DD grant no. 115766); Janssen; Merck KGaA; Darmstadt, Germany; MSD; Novartis Pharma AG; Ontario Ministry of Economic Development and Innovation; Pfizer; São Paulo Research Foundation-FAPESP; Takeda; and the Wellcome Trust. The X-ray structure results of CARM1 are derived from work performed at Argonne National Laboratory, Structural Biology Center (SBC) at the Advanced Photon Source. SBC-CAT is operated by U Chicago Argonne, LLC, for the U.S. Department of Energy, Office of Biological and Environmental Research under contract DE-AC02-06CH11357. These experiments were performed using beamline 08ID-1 at the Canadian Light Source, which is supported by the Canada Foundation for Innovation, Natural Sciences and Engineering Research Council of Canada, the University of Saskatchewan, the Government of Saskatchewan, Western Economic Diversification Canada, the National Research Council Canada, and the Canadian Institutes of Health Research.

## Author Contributions

X.C.C., K.W., W.Z., and J.P.L. developed synthetic strategies and prepared compounds; X.C.C., J.W., C.S., T.H., G.I., M.V., F.L., C.C.S., and T.H. characterized the compounds with in vitro biochemical and biophysical assays; H.W., F.L., C.C.S., H.Z., A.D., and L.D. solved the X-ray crystal structures of CARM1 in complex with ligands; V.V. and L.S. modeled the interaction of CARM1 with ligands; E.J.K., X.C.C., M.J., N.Z., Y.C. D.B. and M.S. examined on-target engagement in a cellular context; S.C. and X.C.C. modeled target occupancy; E.J.K., X.C.C., M.J., and N.Z. conducted invasion assay; X.C.C., M.J., and L.M. performed scRNA-seq; T.Z., M.L., L.X.Q., X.C.C., and X.N. analyzed scRNA-seq data; X.C.C., M.L., W.X., L.X.Q., J.X., P.J.B., M.V., L.S., C.H.A., J.M., H.D. and Z.Z. designed experiments and supervised the projects; all of the authors analyzed the data; X.C.C. and M.L. wrote the manuscript with inputs of other authors.

## Additional information

**PDB codes**: 4IKP for the CARM1-**1** complex and 6D2L for CARM1-**5a** complex

