## Supplemental Information for "A Chemical Probe of CARM1 Alters Epigenetic Plasticity against Breast Cancer Cell Invasion"

**CONTENTS**

**1. Supplementary Results**

**The X-ray structure of the CARM1-1(HSF) complex.**

**2. Supplementary Methods**

2.1. **Synthesis and characterization of 3, 1 (HSF), 2a, 5a, 6a (SKI-73), 2b, 5b and 6b (SKI-73N)**
2.2. **Determination of IC_50_ of SAH, SNF, 1, 2a, 5a, 5b and 6a**
2.3. **Determination of IC_50_ and *K*_d_ values of CARM1 inhibitors and *K*_m,SAM_ in the presence of the varied concentrations of SAM and peptide substrate**
2.4. **Surface Plasmon Resonance (SPR)**
2.5. ***In vitro* thermal shift assay (TSA)**
2.6. **Crystallization of CARM1 in complex with 1 and 5a**
2.7. **Molecular docking and MD simulations of CARM1-2a and CARM1-SNF complexes**
2.8. **LC-MS/MS quantification of intracellular concentrations of 6a, 5a, 2a and SAM**
2.9. **Modeling of the percentages of intracellular apo-CARM1, CARM1 occupied by 6a, 5a, and 2a, and SAM-bond CARM1 versus the total amount of CARM1**
2.10. **Evaluation of methylation marks of CARM1 in breast cancer cells**
2.11. **Cellular thermal shift assay (CETSA)**
2.12. **Matrigel invasion assay**
2.13. **Sample preparation for single cell RNA-seq analysis (scRNA-seq)**
2.14. **Processing and preparation of scRNA-seq data**
2.15. **Subpopulation clustering guided by scRNA-seq**
2.16. **Correlation analysis of subpopulations, heat map analysis, and selection of representative transcripts.**

**3. Supplementary Figures and Tables**.

**1. Supplementary Results**

**The X-ray structure of the CARM1-1(HSF) complex.** Given the tight CARM1 binding by **1**, we solved the X-ray structure of CARM1 in complex with **1** (**HSF**). The overall folding of the CARM1-**1** (PDB: 4IKP) is similar to those in complex with **SNF** and **SAH** (PDB: 2Y1X, 2Y1W) with a V-shape subunit in a dimer of a dimer (Figure S4a).(Sack et al., 2011) However, the CARM1-**1** complex is distinct for multiple configurations of its ligand (Figure S4b) and interactions via its 6′-methyleneamine moiety (Figure S5). In the CARM1-**1** complex, **1** can adopt four alternative configurations (Configuration I in Chains B, D; Configuration II in Chain C; Configuration III, IV in Chain A) accompanied with structural accommodation of the adjacent residues and water hydrogen bonds (Figure S4b, S5 and Tables S2-4). In contrast, only a single configuration of the ligands was observed in **SAH**- or **SNF**-bound CARM1 (Figure S5a,c, PDB: 2Y1X, 2Y1W).(Sack et al., 2011)

Detailed structural comparison of CARM1 in complex with **SNF**, **SAH** and **1** further revealed that **1** maintains common interactions observed in the CARM1-**SNF** and CARM1-**SAH** complexes with several exceptions (Figure S5 and Table S2-4). Most noticeably, the CARM1-**1** complex gains the strong hydrogen bonds via the 6′-methyleneamine moiety of the ligand with (i) the backbone carbonyl of CARM1’s Glu258 with the average distance of 2.6 Å (Configuration I, II in Chains B, C, D and Configuration III in Chain A, Figure S5) or (ii) the side chain of Tyr154 (Configuration IV in Chain A, Figure S5), together with multiple less conserved water hydrogen bonds bridged to Y150, Y154, E258, M260, or E267 (Table S3). In contrast, the 6′-amine of **SNF** in the CARM1-**SNF** complex forms weaker hydrogen bonds with Glu258 of the average distance of around 3.0 Å and far less water hydrogen bonds (Figure S5, Table S2, S3 and PDB: 2Y1W). Comparable interactions are completely absent in the CARM1-**SAH** complex (Figure S5c, d, Table S2-4 and PDB: 2Y1X). The desired hydrogen-bond networks of the 6′-methyleneamine moiety of **1** with CARM1, which are present in the CARM1-**1** complex but absent in the CARM1-**SNF** and CARM1-**SAH** complexes, can rationalize the significant decrease of IC_50_ from **SNF** and **SAH** to **1**.

Another key difference among CARM1-**1**, CARM1-**SNF** and CARM1-**SAH** complexes lies in the region around the carboxylic moiety of these ligands. In Chain A, C of the CARM1-**1** complex, the carboxylic moiety of the ligand forms an ionic bond with Arg169 and a hydrogen bond with Gln160 (Figure S4b, S5a, c and Table S2, S3). Such interactions are absent in CARM1-**SNF** and CARM1-**SAH** complexes (Figure S5c). In contrast, in Chain B, D of the CARM1-**1** complex, the same carboxylic moiety forms the ionic bonds with Arg169 and a water hydrogen bond bridged to Gln160 and Ser196 (Figure S4b, 5a, 5b, 5d and Table S2, S3). To accommodate the latter conformation, the Gln160 residue flips toward the 3′-ribosyl hydroxyl moiety of **1** to form a new hydrogen bond (Figure S4b, S5d, Table S2, S3). Similar interaction patterns can also be found in CARM1-**SNF** and CARM1-**SAH** complexes (Figure 5d).

In regard with rest CARM1-ligand interactions, the CARM1 complexes with **1**, **SAH** and **SNF** are nearly identical except slightly altered water hydrogen bonds (Figure S5 and Table S2-4). Here the α-amino moiety of these ligands forms hydrogen bonds with the carbonyl backbone of Gly193, as well as two water hydrogen bonds bridged toward the regions of CARM1’s Glu191, Val192, Gly193, Cys194, Leu199, Ser257 and Asp258; their 2′,3′-ribosyl hydroxyl groups form two hydrogen bonds with the side chain of CARM1’s Glu215; adenine’s N1 and N6 form the hydrogen bonds with Asn243 and Glu244/Ser272, respectively; the adenine ring of these ligands is buried within a hydrophobic pocket consisting of the side chains of Phe151, Val192, Val214, Ala216, Val243, Met269, the hydrophobic portion of Lys242’s side chain and Ser217’s backbone. In contrast, there are less conserved water hydrogen bonds, such as those involved with the carboxylic and 3′-ribosyl hydroxyl moieties of **1** (H_2_O 753, H_2_O 777, and H_2_O 784 in Chain A of the CARM1-**1** complex, Table S3). In contrast, adenine-N7 in the CARM1-**SNF** and CARM1-**SAH** complexes forms water hydrogen bonds bridged to Ser272, which are absent in the CARM1-**1** complex (Table S3, S4). Collectively, the general high affinity of **1**, **SNF** and **SAH** (Table S2-4) arises from the combined hydrophilic and hydrophobic interactions of these ligands with CARM1. However, in comparison with **SNF** and **SAH**, **1** gains the extra interactions via its 6′-methyleneamine moiety (Figure S5 and Table S2-4). Additionally, **1** adopts the canonical pose with its α-amino carboxylate moiety interacting with Arg168, which is similar to that of **SNF** and **SAH** but different from the noncanonical pose of **2a** and **5a**, upon binding CARM1.

**2. Supplementary Methods**

**2.1 Synthesis and characterization of 3, 1, 2a, 5a, 6a, 2b, 5b and 6b**

**2.1.1** **Abbreviations in chemical structures.** Ac: acetyl; Bn: benzyl; Bz: benzoyl; Cbz: benzyloxycarbonyl; TFA: trifluoroacetyl.

**2.1.2 Abbreviations of chemical reagents.** CbzCl: benzyl chloroformate; DCM: dichloromethane; DMF: *N, N*-dimethylformamide; HATU: 1-[bis(dimethylamino)methylene]-1H-1,2,3-triazolo[4,5-b] pyridinium 3-oxid hexafluorophosphate; TFA: trifluoroacetic acid; THF: tetrahydrofuran; TML-NHS: *N*-hydroxysuccinimidyl ester 3-methyl-3-(2,4,5-trimethyl-3,6-dioxocyclo-hexa-1,4-dienyl)-butanoic acid; SAH: *S*-adenosyl homocysteine.

**2.1.3 General experimental information.** Reagents for chemical reactions were purchased from Sigma-Aldrich without purification unless mentioned otherwise. Anhydrous solvents were prepared from solvent purification system (PURE SOLV^TM^, Innovative Technology, Inc.). Chemical reactions were carried out under argon atmosphere at the temperatures displayed by thermocouple or at ambient temperature (22 °C) unless described otherwise. The phrase ″concentrated″ in the synthetic method session refers to the reaction workup to remove volatile solvents by a rotary evaporator attached with a diaphragm pump (15-20 Torr) and then by a high vacuum pump (<1 Torr). Chromatography purification was carried out with silica gel from Dynamic Adsorbents, Inc. (neutral, 32-63 µm). NMR spectra were recorded on Bruker AVIII 600MHz spectrometers and reported with chemical shifts (ppm), multiplicities (s = singlet, d = doublet, t = triplet, q = quartet, p = pentet, m = multiplet, and br = broad), integration and coupling constants (*J* in Hz). Chemical shifts were recorded with residual proton peaks of deuterated solvents as references (residual ^1^H of DMSO, 2.50 ppm; CD_3_OD, 3.31 ppm; D_2_O, 4.80 ppm; ^13^C of DMSO, 39.52 ppm; CD_3_OD, 49.00 ppm). ^1^H-NMR spectra were recorded at 24.0 °C or at 70.0 °C. ^1^H-NMR spectra at 70.0 °C were recorded in DMSO-*d_6_* to facilitate the equilibrium between rotamers. ^13^C-NMR spectra were recorded at 24 °C. Mass spectra for compound characterization were collected by Waters Acuity SQD LC-MS with the mode of electron spray ionization (ESI). The final concentrations of the stock solutions of **3**, **1**, **2a**, **5a**, **5b**, SAH and nonradioactive SAM were determined on the basis of their UV absorption at 260 nm (ε_260_ = 15,400 L.mol^-1^.cm^-1^) with Nanodrop 1000 Spectrophotometer (Thermo Scientific). The final concentrations of **6a** and **6b** were determined with ^1^H-NMR in CD_3_OD containing 1.0 mM SAH as an internal reference for the first time and with Nanodrop 1000 Spectrophotometer (Thermo Scientific) on the basis of their UV absorption at 267 nm (ε_267_ = 19,300 L.mol^-1^.cm^-1^) thereafter. CD_3_-SAM was prepared as described previously.(Linscott et al., 2016)

**Synthesis of 3.** Compound **3** was prepared as previously reported.(Wu et al., 2016) Briefly, into an oven-dried flask was added *N*^6^-benzoyladenine (44 mg, 0.18 mmol), hexamethyldisilazane (3 mL) and pyridine (1 mL). The suspension was heated at 115 °C to afford a clear solution and stirred for another 3 h. The mixture was concentrated under reduced pressure. The residual was dried via azeotrope with toluene (3×5 mL) and over high vacuum. To the solid crude of bis-sily*-N*^6^-benzoyladenine was added a solution of triacetate derivative (0.037 mmol) in 15 mL 1,2-dichloroethane. The resultant suspension was treated with TMSOTf (33 µL, 0.18 mmol) and heated at 50 °C for 2 h. The mixture was quenched with 10 mL saturated NaHCO_3_, followed by extraction with 3×20 mL CH_2_Cl_2_. The combined organic phases were washed with brine, dried with anhydrous Na_2_SO_4,_ and evaporated to afford 56 mg crude of **3**. Flash silica gel chromatography with MeOH/DCM = 30:1 afforded 30 mg of **3** as a white solid (92% yield).

**3**: ^1^H-NMR (600 MHz, DMSO-*d_6_*, 84 °C): δ 10.76(s, 1H), 9.39(d, 1H, *J* = 6.4), 8.71(s, 1H), 8.59(s, 1H), 8.04(d, 2H, *J* = 7.6), 7.63(t, 1H, *J* = 7.4), 7.54(t, 2H, *J* = 7.6), 7.22-7.31(m, 8H), 7.12-7.13(m, 2H), 6.23(d, 1H, *J* = 5.6), 6.04(t, 1H, *J* = 5.6), 5.41(t, 1H, *J* = 5.6), 5.10(d, 2H, *J* = 5.5), 4.45(d, 1H, *J* = 15.6), 4.39(d, 1H, *J* = 15.6), 4.24-4.28(m, 1H), 4.15-4.18(m, 1H), 3.66(s, 3H), 3.23-3.26(m, 1H), 3.13-3.16(m, 1H), 2.10(s, 3H), 2.04(s, 3H), 1.92-1.95(m, 1H), 1.81-1.88(m, 2H), 1.75-1.80(m, 1H), 1.64-1.67(m, 1H), 1.31-1.39(m, 2H); ^13^C-NMR (150 MHz, DMSO-*d_6_* *rotamers*): δ 170.92, 169.60, 169.51, 169.40, 165.67, 156.59(q, *J* = 36.4), 155.87, 151.85, 151.80, 150.71, 143.88, 137.85, 136.77, 133.23, 132.57, 128.54, 128.52, 128.43, 128.32, 128.29, 127.79, 127.51, 127.39, 127.28, 127.12, 126.90, 126.01, 115.77(q, *J* = 286.1), 85.85, 85.75, 79.17, 73.17, 71.89, 66.52, 66.38, 54.94, 52.82, 34.25, 32.81, 32.41, 30.72, 27.48, 26.70, 20.42, 20.38, 20.25; MS(ESI) m/z: 940 ([M+Na]^+^ ; HRMS: calculated for C_45_H_47_N_7_O_11_F_3_ ([M+H] ^+^)918.3286, found 918.3311.

**Synthesis of 1 and 2a from 3 (Figure S1).** To a solution of **3** (45 mg, 0.050 mmol) in 15 mL CF_3_CH_2_OH was added Pd/C (22 mg, 10 wt. %, wet support, Degussa type). This mixture was stirred under H_2_ (1 atm) at ambient temperature (22 °C) for 36 h.(Bailey et al., 2008) The reaction mixture was filtered through a short pad of Celite, followed by washing with 150 mL of MeOH. The combined filtrate was concentrated to afford around 1:1 crude mixture of **S1** and **S2** (Figure S1). This crude material was dissolved in 2 mL of MeOH and then mixed with 1 mL of 0.2 M LiOH. The resultant mixture was stirred at ambient temperature (22 °C) for 40 h, neutralized with 0.2 M HCl to reach pH = 7, and concentrated under reduced pressure to afford the crude product mixture of **1** and **2a**. The crude products were then subject to a preparative reversed-phase HPLC (XBridge™ Prep C18 5μm OBD™ 19×150 mm) with 5−95% gradient of CH_3_CN in aqueous trifluoroacetic acid (TFA, vol. 0.1%) in 24 min with a flow rate of 10 mL/min to afford 6.5 mg of **1** (27% yield over two steps) and 7.5 mg of **2a** (31% yield over two steps) as white solids.

**1**: ^1^H-NMR (600 MHz, D_2_O): δ 8.36(s, 1H), 8.35(s, 1H), 6.01(d, 1H, *J* = 3.7), 4.65 (dd, 1H, *J* = 3.7, 5.4), 4.36(t, 1H, *J =* 5.9), 4.22-4.19 (m, 1H), 4.03-3.99 (m, 1H), 3.82(t, 1H, *J* = 6.0), 3.05 (dd, 1H, *J* = 5.4 12.8), 2.96 (dd, 1H, *J* = 6.9, 12.8), 1.97-1.84 (m, 5H), 1.49-1.55(m, 2H); ^13^C-NMR (150 MHz, D_2_O): δ 173.33, 150.08, 148.17, 144.67, 142.76, 119.02, 115.28, 88.81, 81.18, 73.35, 53.79, 41.94, 33.65, 33.12, 26.88, 25.84; HRMS: calculated for C_16_H_26_N_7_O_5_ ([M+H]^+^) 396.1995, found: 396.1982.

**2a**: ^1^H-NMR (600 MHz, CD_3_OD): δ 8.16(s, 1H), 8.09(s, 1H), 7.25(d, 1H, *J* = 7.4), 7.20(t, 2H, *J* = 7.4), 7.12(t, 2H, *J* = 7.4), 5.84(d, 1H, *J* = 4.0), 4.49(dd, 1H, *J* = 5.4 , 4.0), 4.08 (t, 1H, *J =* 5.7 ), 4.04(d, 1H, *J* = 13.0), 3.98-4.00(m, 1H), 3.95(d, 1H, *J* = 13.0), 3.82(t, 1H, *J* = 7.0), 2.94-3.00(m, 1H), 1.92-2.03(m, 2H), 1.83-1.91(m, 2H), 1.76-1.80(m, 1H), 1.46-1.56(m, 2H); ^13^C-NMR (150 MHz, CD_3_OD): δ 171.94, 155.85, 151.31, 150.18, 142.68, 132.10, 130.90 (2×^13^C), 130.83, 130.37 (2×^13^C), 121.19, 91.65, 82.07, 75.36, 74.88, 54.07, 53.01, 52.07, 34.69, 34.32, 28.72, 27.90. HRMS: calculated for C_23_H_32_N_7_O_5_ ([M+H]^+^) 486.2465, found:486.2464.

**Synthesis of 5a and 6a (SKI-73) from 3 (Figure S2).** To a solution of **3** (91 mg, 0.10 mmol) in 20 mL MeOH was added 10 mL of 0.2 M LiOH. The resultant mixture was stirred at ambient temperature (22 °C) for 40 h. The reaction mixture was then neutralized with 0.2 M HCl to reach pH = 7.0 and concentrated under reduced pressure to afford **4** without further purification. The crude product **4** was then dissolved in 10 mL of THF and then mixed with 3 mL of saturated aqueous NaHCO_3_ and CbzCl (12 μL, 0.10 mmol) at 0 °C. This mixture was stirred at 0 °C for 3 h, quenched with 0.2 M HCl to reach pH = 7.0 and concentrated under reduced pressure. The resultant solid was washed with 100 mL THF, filtered and concentrated to afford 65 mg of the crude product **S3** without further purification. To a solution of **S3** in 8 mL DMF were sequentially added HATU (114 mg, 0.3 mmol), 2,3,5-collidine (39 μL, 0.3 mmol) and 4-methoxyphenethylamine (44 μL, 0.3 mmol). (Han and Kim, 2004) The resultant mixture was stirred at ambient temperature (22 °C) under argon until the starting material **S3** was fully consumed as monitored by LC-MS. The reaction was then quenched with 3 mL of saturated aqueous NH_4_Cl, followed by extraction with 3×30 mL DCM. The combined organic layers were washed with brine, dried with anhydrous Na_2_SO_4_, filtered, and evaporated to give the crude solid product **S4**. This crude product was purified by a flash silica gel chromatography (v/v 1:12, MeOH/DCM) to give 55 mg of **S4** as a white solid (62% yield over three steps).

**S4**: ^1^H-NMR (600 MHz, DMSO-*d_6_*, 70 °C): δ 8.28 (s, 1), 8.24 (s, 1), 7.53 (br, 1), 7.35-7.20 (m, 13), 7.11 (d, 2, *J* = 7.02), 7.07 (d, 2, *J* = 8.52), 6.81 (d, 2, *J* = 8.52), 5.86 (d, 1, *J* = 5.03), 5.10 (d, 1, *J* = 12.9), 5.08 (d, 1, *J* = 12.9), 5.03 (s, 2), 4.60 (t, 1, *J* = 5.03), 4.47 (d, 1, *J* = 16.2), 4.38 (d, 1, *J* = 16.2), 3.98 (t, 1, *J* = 5.03), 3.93-3.87 (m, 2), 3.69 (s, 3), 3.53-3.50 (m, 2), 3.16-3.10 (m, 2), 2.63 (t, 2, *J* = 7.14), 1.96-1.85 (m, 1), 1.71-1.58 (m, 3), 1.50-1.41 (m, 1), 1.39-1.32 (m, 1), 1.28-1.20 (m, 1). ^13^C NMR (150 MHz , DMSO-*d_6_*_,_ *rotamers*): δ 172.05 (15), 157.93, 156.47, 156.41, 156.23, 153.04 , 149.04, 141.88, 138.18, 137.26 (2×C), 131.48, 129.53 (2×C), 128.80 (2×C), 128.72 (2×C), 128.70 (2×C), 128.67, 128.19, 128.13, 128.03, 127.98, 127.68 (2×C), 127.55, 127.17, 119.29, 113.99 (2×C), 88.48, 82.88 (d), 73.86, 73.17, 66.82 (d), 65.80, 55.38, 55.23, 50.40 (2×C) 40.62 (2×C), 35.16, 34.46, 33.72, 33.37, 29.25(d), 28.03 (d). HRMS: calculated for C_48_H_55_N_8_O_9_ ([M+H^+^]) 887.4092, found: 887.4082.

To a solution of **S4** (20 mg, 0.022 mmol) in 20 mL CF_3_CH_2_OH was added Pd/C (10 mg, 10 wt. %, wet support, Degussa type). The resultant mixture was stirred under H_2_ (1 atm) at ambient temperature (22 °C) until the starting material **S4** was fully consumed as monitored by LC-MS. The reaction mixture was filtered through a short pad of Celite, followed by washing with 150 mL of MeOH. The combined filtrate was concentrated under reduced pressure to give the crude product **5a**. This crude material is further subject to a preparative reversed-phase HPLC (XBridge™ Prep C18 5μm OBD™ 19×150 mm) with 5−95% gradient (volume ratio) of CH_3_CN in aqueous trifluoroacetic acid (TFA, vol. 0.1%) in 24 min with a flow rate of 10 mL/min to afford 5.4 mg of **5a** as a white solid (40% yield).

**5a**: ^1^H-NMR (600 MHz, CD_3_OD): δ 8.31 (s, 1), 8.21 (s, 1), 7.36 (t, 1, *J* = 7.32), 7.30 (t, 2, *J* = 7.32), 7.25 (d, 2, *J* = 7.32), 7.09 (d, 2, *J* = 8.58), 6.82 (d, 2, *J* = 8.58), 5.96 (d, 1, *J* = 3.84), 4.56 (dd, 1, *J* = 3.84, 5.28), 4.17 – 4.14 (m, 2), 4.09 – 4.03 (m, 2), 3.76 – 3.72 (m, 1), 3.78 (s, 3), 3.53 – 3.48 (m, 1), 3.40 – 3.34 (m, 1), 3.03 - 2.97 (m, 2), 2.78 - 2.69 (m, 2), 2.09 – 2.05 (m, 1), 2.05 – 1.99 (m, 1), 1.91 – 1.82 (m, 2), 1.74 - 1.68 (m, 1), 1.48 – 1.43 (m, 1), 1.41 – 1.35 (m, 1). ^13^C NMR (150 MHz , CD_3_OD): δ 169.72, 159.80, 154.63, 149.98, 149.65, 142.95, 132.08, 132.05, 130.81 (2×C), 130.78(2×C), 130.69, 130.22 (2×C), 121.00, 115.03 (2×C), 91.51, 82.20, 75.30, 74.90, 57.63, 55.74, 54.32, 52.88, 42.17, 36.65, 34.59, 34.37, 29.72, 27.64. HRMS: calculated for C_32_H_43_N_8_O_5_ ([M+H^+^]) 619.3356, found: 619.3326.

To a solution of **5a** (6 mg, 0.01 mmol) in 1 mL anhydrous DMF was added Et_3_N (12 μL, 0.05 mmol) and *N*-hydroxysuccinimidyl ester 3-methyl-3-(2,4,5-trimethyl-3,6-dioxocyclo-hexa-1,4-dienyl) butanoic acid (TML-NHS ester) (3.5 mg, 0.01 mmol) at 0 °C.(Rohde et al., 2006) The resultant mixture was stirred under argon for 8 h, quenched with 2 mL of saturated aqueous NH_4_Cl solution and then concentrated under reduced pressure. The resultant product **6a** was purified a preparative reversed-phase HPLC (XBridge™ Prep C18 5μm OBD™ 19×150 mm) with 5−95% gradient (volume ratio) of CH_3_CN in aqueous trifluoroacetic acid (TFA, vol. 0.1%) in 24 min with a flow rate of 10 mL/min to afford 2.8 mg of **6a** as a yellow solid (33% yield).

**6a**: ^1^H-NMR (600 MHz, CD_3_OD): δ 8.26 (s, 1), 8.19 (s, 1), 7.38-7.31 (m, 3), 7.26-7.24 (m, 2),  7.08 (d, 2, *J* = 8.40), 6.82 (d, 2, *J* = 8.40),  5.93 (d, 1, *J* = 4.50), 4.62 (t, 1, *J* = 4.50), 4.15 - 4.05 (m, 5), 3.76 (s, 3), 3.25 – 3.20 (m, 2), 3.05 (dd, 1, *J* = 6.00, 6.90),  2.99 (dd, 1, *J* = 6.00, 6.90),  2.90 (d, 1, *J* = 14.8), 2.79 (d, 1, *J* = 14.8), 2.67 (t, 2, *J* = 7.20), 2.08 (s, 3), 2.07- 2.00 (m, 1), 2.08 – 1.98 (m, 2), 1.94 (s, 3), 1.91 (s, 3), 1.92 - 1.82 (m, 1), 1.78 - 1.72 (m, 1), 1.41 (s, 3), 1.39 (s, 3), 1.38 – 1.29 (m, 2). ^13^C NMR (150 MHz , CD_3_OD): δ 192.12, 188.69, 174.46, 173.71, 159.76, 155.11, 154.3, 149.97, 149.24, 144.93, 143.15, 138.81, 138.71, 132.23, 132.09, 130.87 (2×C), 130.83 (2×C), 130.76, 130.27 (2×C), 120.99, 114.97 (2×C), 91.30, 82.60, 75.32, 74.89, 55.70, 54.22, 52.86, 51.80, 49.58, 42.09, 39.56, 35.52, 35.35, 33.99, 29.58, 29.27, 29.21, 28.70, 14.42, 12.93, 12.04. HRMS: calculated for C_46_H_59_N_8_O_8_ ([M+H^+^]) 851.4456, found: 851.4431.

**Synthesis of 5b and 6b (SKI-73N) from S5 (Figure S3).** Compound **S5** was prepared as described previously.(Zheng et al., 2012) Into a solution of **S5** (94 mg, 0.10 mmol) in 20 mL MeOH solution was added 10 mL of 0.2 M LiOH. The resultant mixture was stirred at ambient temperature (22 °C) for 20 h. The mixture was then neutralized with 0.2 M HCl to reach pH = 7 and concentrated under reduced pressure to afford the crude product **S6** without further purification. To a solution of **S6** in 8 mL anhydrous DMF was sequentially added HATU (114 mg, 0.3 mmol), 2,3,5-collidine (39 μL, 0.3 mmol) and 4-methoxyphenethylamine (44 μL, 0.3 mmol). The resultant mixture was stirred at ambient temperature (22 °C) under argon until the starting material **S6** was fully consumed as monitored by LC-MS. The reaction was then quenched with 3 mL of saturated aqueous NH_4_Cl, followed by the extraction with 3×30 mL DCM. The combined organic layers were washed with brine, dried with anhydrous Na_2_SO_4_, filtered, and evaporated to give the crude product **S7**. This crude product was purified by a flash silica gel chromatography (v/v 1:12, MeOH/DCM) to afford 61 mg of **S7** as a white solid (71% yield).

**S7**: ^1^H-NMR (600 MHz, DMSO-*d_6_*, 70 °C): δ 8.29 (s, 1), 8.24 (s, 1), 7.50-7.47 (br, 1), 7.37-7.33 (m, 2), 7.28-7.20 (m, 12), 7.19-7.17 (m, 1), 7.07 (d, 2, *J* = 8.58), 6.81 (d, 2, *J* = 8.58), 5.84 (d, 1, *J* = 5.04), 5.07 (s, 2), 5.03 (d, 1, *J* = 12.7), 5.00 (d, 1, *J* = 12.7), 4.61 (t, 1, *J* = 5.04), 4.42 (d, 1, *J* = 15.8), 4.32 (d, 1, *J* = 15.8), 3.94 (t, 2, *J* = 5.04), 3.84 - 3.81 (m, 2), 3.70 (s), 3.25 – 3.20 (m, 2), 2.60 (t, 2, *J* = 7.14), 2.09 - 2.01 (m, 1), 1.85 - 1.82 (m, 1), 1.67 – 1.58 (m, 1), 1.56 - 1.46 (m, 2), 1.42 - 1.33 (m, 1). ^13^C NMR (150 MHz , DMSO- *d_6_, rotamers*): δ 171.45 (d), 157.57, 156.55, 155.81, 155.77, 152.22 (d), 148.55, 148.60, 141.95, 139.11, 137.01, 136.69 (d), 131.16, 129.53 (2×C), 128.30 (2×C), 128.22 (2×C), 128.18 (2×C), 128.08 (2×C), 127.76, 127.64 (2×C), 127.26, 127.05, 126.96, 126.71, 119.10, 113.62 (2×C), 88.06, 81.56 (d), 73.52 (2×C), 73.09 (2×C), 66.47 (d), 65.35, 54.87, 54.84, 54.65, 40.38, 36.66 (d), 34.17, 29.47, 29.33. HRMS: calculated for C_47_H_53_N_8_O_9_ ([M+H^+^]) 873.3936, found: 873.3928.

To a solution of **S7** (20 mg, 0.022 mmol) in 20 mL CF_3_CH_2_OH was added Pd/C (10 mg, 10 wt. %, wet support, Degussa type). This mixture was stirred under H_2_ (1 atm) at ambient temperature (22 °C) until the starting material **S7** was fully consumed as monitored by LC-MS. The reaction mixture was then filtered through a short pad of Celite, followed by washing with 150 mL of MeOH. The combined filtrate was concentrated under reduced pressure and purified with a preparative reversed-phase HPLC (XBridge™ Prep C18 5μm OBD™ 19×150 mm) with 5−95% gradient (volume ratio) of CH_3_CN in aqueous trifluoroacetic acid (TFA, vol. 0.1%) in 24 min with a flow rate of 10 mL/min to afford 5.4 mg of **5b** as a white solid (40% yield).

**5b**: ^1^H-NMR (600 MHz, CD_3_OD): δ 8.28 (s), 8.14 (s), 7.30 (t, 1, *J* = 7.50), 7.21 (t, 2, *J* = 7.50), 7.13 (d, 2, *J* = 8.58), 7.08 (d, 2, *J* = 7.50), 6.83 (d, 2, *J* = 8.58), 5.96 (d, 1, *J* = 3.96, H1), 4.69 (dd, 1, *J* = 3.96, 5.88), 4.40 (t, 1, *J* = 5.88), 4.18 – 4.21 (m, 1), 4.10 (d, 1, *J* = 12.84), 4.05 (d, 1, *J* = 12.84), 3.85 – 3.83(m, 1), 3.72 (s, 3), 3.47 – 3.43 (m, 3), 2.79 – 2.74 (m, 2), 2.36 (ddd, 1, *J* = 4.32, 9.42, 15.1), 2.25 (ddd, 1, *J* = 3.00, 6.00, 15.1), 1.95 – 1.89 (m, 2), 1.85 – 1.80 (m, 2). ^13^C NMR (150 MHz , CD_3_OD): δ 169.29, 159.87, 156.45, 152.48, 150.06, 142.33, 132.00, 130.74 (2×C), 130.58, 130.44 (2×C), 130.10 (3×C), 121.06, 115.05 (2×C), 91.72, 80.78, 74.63, 74.12, 56.94, 55.68, 53.93, 49.50, 42.37, 35.47, 32.25, 28.71, 26.46. HRMS: calculated for C_31_H_41_N_8_O_5_ ([M+H^+^]) 605.3200, found: 605.3180.

To a solution of **5b** (6 mg, 0.01 mmol) in 1 mL anhydrous DMF was added Et_3_N (12 μL, 0.05 mmol) and *N*-hydroxysuccinimidyl ester 3-methyl-3-(2,4,5-trimethyl-3,6-dioxocyclo-hexa-1,4-dienyl) butanoic acid (TML-NHS ester) (3.5 mg, 0.01 mmol) at 0 °C. The resultant mixture was stirred under argon for 8 h, quenched with 2 mL of saturated aqueous NH_4_Cl solution and then concentrated under reduced pressure. The resultant crude product was purified a preparative reversed-phase HPLC (XBridge™ Prep C18 5μm OBD™ 19×150 mm) with 5-95% gradient (volume ratio) of CH_3_CN in aqueous trifluoroacetic acid (TFA, vol. 0.1%) in 24 min with a flow rate of 10 mL/min to afford 3.0 mg of **6b** (**SKI-73N**) as a yellow solid (35% yield).

**6b (SKI-73N)**: ^1^H-NMR (600 MHz, CD_3_OD): δ 8.31 (s, 1), 8.18 (s, 1), 7.32 (t, 1, *J* = 7.38), 7.24 (t, 2, *J* = 7.38), 7.11 – 7.07 (m, 4), 6.82 (d, 2, *J* = 8.40), 5.96 (d, 1, *J* = 3.78), 4.69 (dd, 1, *J* = 3.78, 5.94), 4.35 (t, 1, *J* = 5.94), 4.19 – 4.16 (m, 2), 4.13 (d, 1, *J* = 12.9), 4.04 (d, 1, *J* = 12.9), 3.74 (s, 3), 3.44 – 3.40 (m, 1), 3.37 – 3.30 (m, 2), 2.92 (d, 1, *J* = 14.9), 2.78 (d, 1, *J* = 14.9), 2.68 (t, 2, *J* = 7.08), 2.38 – 2.32 (m, 1), 2.25 - 2.20 (m, 1), 2.08 (s, 3), 1.94 (s, 3), 1.91 (s, 3), 1.82 - 1.78 (m, 2), 1.74 – 1.70 (m, 1), 1.57 – 1.54 (m, 1), 1.40 (s, 6). ^13^C NMR (150 MHz , CD_3_OD): δ 192.14, 188.66, 174.51, 173.17, 159.79, 155.66, 155.10, 151.29, 150.04, 144.95, 142.82, 138.79, 138.73, 132.22, 130.79 (3×C), 130.61, 130.43 (2×C), 130.13 (2×C), 121.07, 114.98 (2×C), 91.65, 81.14, 74.84, 74.24, 57.62, 55.68, 53.77, 49.50 (2×C), 42.18, 39.57, 35.56, 32.85, 29.29, 29.17, 29.10, 27.76, 14.42, 12.94, 12.04. HRMS: calculated for C_45_H_57_N_8_O_8_ ([M+H^+^]) 837.4299, found: 837.4316.

**2.2 Determination of IC_50_ of SAH, SNF, 1, 2a, 5a, 5b and 6a.**

The IC_50_ assays of **SAH**, **SNF**, **1**, **2a**, **5a** and **5b** against a collection of 34 methyltransferases were performed as previously described except for PRMT9.(Bromberg et al., 2017; Li et al., 2016a; Li et al., 2016b) The IC_50_ values were obtained by fitting inhibition% against the concentrations of the inhibitor to a sigmoid curve.

Briefly, the effects of these compounds on G9a, GLP1, SUV39H1, SUV39H2, SUV420H1, SUV420H2, SETD2, SETD8, SETDB1, SETD7/9, a trimeric complex of MLL1, a pentameric complex of MLL3, a trimeric complex of EZH2 (PRC2), PRMT1, PRMT3, CARM1, a PRMT5/MEP50 complex, PRMT6, PRMT7, PRMT8, PRMT9, PRDM9, SMYD2, SMYD3, DNMT1, BCDIN3D and METTL3-METTL14 were assessed by monitoring the incorporation of the tritium-labeled methyl group of [^3^H-Me]-SAM into substrates using scintillation proximity assay (SPA). Here a 10-µl reaction containing [^3^H-Me]-SAM and a substrate at their concentrations close to the apparent *K*_m_ values for each enzyme was prepared. The reaction was quenched by adding 10 µl of 7.5 M guanidine hydrochloride. Into this mixture was added 180 µL of 20 mM Tris buffer (pH 8.0). This mixture was then transferred to a 96-well FlashPlate followed by incubation for 1 h. The counts per minute (CPM) was measured with a TopCount plate reader. The CPM readouts in the absence of these compounds or these enzymes were defined as 100% activity and background (0%), respectively. For BCDIN3D and METTL3-METTL14, biotinylated RNA strands were used as substrates, For DNMT1, the substrate was a double-stranded DNA with the forward strand biotinylated. For PRMT9, the assay buffer contains 10 nM PRMT9 (aa 1-845), 20 µM SAM, 80 nM biotinylated-SAP145 peptide substrate, 20 mM Tris-HCl (pH=7.5), 5 mM DTT, and 0.01% Triton X-100. Purification of PRMT9 will be reported elsewhere.

A filter-based assay was applied for DOT1L, NSD1, NSD2, NSD3, ASH1L, DNMT3A/3L, and DNMT3B/3L. Here a reaction mixture of 10 µL was incubated at 23 °C for 1 h, followed by adding 50 µL of 10% trichloroacetic acid (TCA) to quench the reaction. The resulting mixture was then transferred to filter plates (Millipore, Billerica, MA, USA), followed by centrifugation at 2,000 r.p.m (Allegra X-15R; Beckman Coulter, Brea, CA, USA) for 2 min, washing twice with 10% TCA and once with 180 µl ethanol. After centrifugation and drying, 100 µL MicroScint-O (Perkin Elmer) was added into each well and the plates were centrifuged to remove the liquid. A 70-μL volume of MicroScint-O was added and the CPM was measured with a TopCount plate reader.

A radiometric filter paper assay described previously(Ibanez et al., 2012; Zheng et al., 2012) was also performed as the alternative assay to determine IC_50_ values of **2a**, **5a** and **6a** against CARM1. Briefly, 20 µL enzymatic reactions (triplicate of each data point) were carried out in the assay solution containing 50 mM HEPES-HCl (pH = 8.0), 0.005% Tween-20 (v/v), 0.0005% BSA, 1 mM TCEP, 25 nM CARM1, 1.5 μM H3 peptide (aa 1-40), 0.75 μM [^3^H-Me]-SAM (PerkinElmer, 5-15Ci/mmol in 9:1 v/v of sulfuric acid and ethanol), and the varied concentrations of inhibitors. For the signal readout of no inhibition (high readout), the same reaction (triplicate) was carried out except with inhibitors replaced with DMSO. For the signal readout of 100% inhibition (background readout), the same reaction (triplicate) was carried out except with inhibitors replaced with DMSO and with the omission of 25 nM CARM1. Here a 10 µL solution containing 50 nM CARM1 and an examined inhibitor at 2× concentrations was incubated at 22 °C for 30 min. To the 10 µL mixture of CARM1 and an inhibitor was added 10 µL of the assay buffer containing 3.0 μM H3 peptide (aa 1-40) and 1.5 μM [^3^H-Me]-SAM. The resultant reaction was carried out at ambient temperature (22 °C) for 6 h, which is in the linear range of the increase of signal readouts. The reactions were then quenched by spotting the reaction mixture onto P81 ion-exchange cellulose chromatography paper (Fisher) to immobilize the peptide. The filter paper was washed 3 times with 50 mM NaHCO_3_/Na_2_CO_3_ buffer (pH = 9.2) to remove the unreacted [^3^H-Me]-SAM. The immobilized peptide containing [^3^H-Me] was quantified by a scintillation counter (Tri-Carb 2810TR, PerkinElmer). To determine the IC_50_ value of an inhibitor against CARM1, inhibition% at specific concentrations was calculated as [(CPM of high readout – CPM readout in the presence of the inhibitor)/(CPM of high readout – CPM of background readout)]×100%. The IC_50_ values were obtained by fitting inhibition% against the concentrations of the inhibitor to a sigmoid curve with GraphPad Prism.

**2.3 Determination of IC_50_ and *K*_d_ values of CARM1 inhibitors and *K*_m,SAM_ in the presence of the varied concentrations of SAM and peptide substrate**. SAM- and substrate-dependent IC_50_ values were determined with the radiometric filter paper assay as described above except in the presence of the varied concentrations of the SAM cofactor and the peptide substrate. The IC_50_ values of **2a** and **5a** were obtained in the presence of 0.09, 0.18, 0.37, 0.75, 1.50, 2.25, 3.00, 5.62, and 7.50 µM [^3^H-Me]-SAM or 0.3, 1.5, 7.5, 15, 30, and 50 µM H3 peptide (aa 1-40). The resultant IC_50_ values were plotted as a function of the concentrations of SAM and the peptide substrate using GraphPad Prism Software. Given the SAM-competitive character of **2a** and **5a**, their SAM-dependent IC_50_ values were then fitted according to eq. S1. Here [SAM] is the concentration of SAM, *K*_m,SAM_ is the Michaelis-Menten constant, and *K*_d_ is the dissociation constant of the CARM1 inhibitor **2a** or **5a**. *K*_d_, *K*_d_/*K*_m,SAM_ and *K*_m,SAM_ can therefore be obtained through the intercept of the y-axis, the slop, and their ratio, respectively, upon fitting eq. S1. Given that **6a** showed a much higher IC_50_ (1.1 ± 0.1 µM, Figure S10), the *K*_d_ value of **6a** (*K*_d,_**_6a_** = 0.32 µM) was directly obtained from the IC_50_ value, the *K*_m,SAM_ derived above, and the assayed SAM concentration according to eq. S1.

IC_50_ = [SAM]×*K*_d_/*K*_m,SAM_+*K*_d_	eq. S1

**2.4 Surface Plasmon Resonance (SPR)**.

Full-length CARM1 was used for SPR assay. DNA fragment encoding the full-length CARM1 was cloned into pFB-N-flag-LIC donor plasmid. The resulting plasmid was transformed into DH10Bac Competent *E. coli* cells (Invitrogen) and a recombinant Bacmid DNA was purified, followed by a recombinant baculovirus generation in Sf9 insect cells. Sf9 cells grown in HyQ^®^ SFX insect serum-free medium (ThermoScientific) were infected with 10 mL of P3 viral stocks per 1 L of suspension cell culture and incubated at 27°C using a platform shaker set at 150 revolutions per minute. The cells were collected when viability dropped to 70-80% (~72 hours after infection). Harvested cells were re-suspended in PBS, 1X protease inhibitor cocktail (100X protease inhibitor stock in 70% ethanol containing 0.25 mg/ml Aprotinin, 0.25 mg/ml Leupeptin, 0.25 mg/ml Pepstatin A and 0.25 mg/ml E-64) and 2X Roche complete EDTA-free protease inhibitor cocktail tablet. The cells were lysed chemically by rotating 30 min with NP40 (final concentration of 0.6%) and 50 U/mL Benzonase nuclease (Sigma), 2 mM 2-mercaptoethanol and 10% Glycerol followed by sonication at frequency of 7 (10′′ on/10′′ off) for 2 min (Sonicator 3000, Misoni). The crude extract was clarified by high-speed centrifugation (60 min at 36,000 ×g at 4˚C) by Beckman Coulter centrifuge. The recombinant protein was purified by incubating the cleared lysate with anti-FLAG M2 affinity agarose gel (Sigma, Cat # A2220) and then rotating for 3 hours, followed by washing with 10 CV TBS (50 mM Tris-HCl, 150 mM NaCl, pH 7.4) containing 2 mM 2-mercaptoethanol, 1X protease inhibitor cocktail (100X protease inhibitor stock in 70% ethanol containing 0.25 mg/ml Aprotinin, 0.25 mg/ml Leupeptin, 0.25 mg/ml Pepstatin A and 0.25 mg/ml E-64) and 1X Roche complete EDTA-free protease inhibitor cocktail tablet. The recombinant protein was eluted by competitive elution with a solution containing 100 µg/mL FLAG peptide (Sigma, Catalog # F4799) in 20 mM Tris pH:7.4, 150 mM NaCl, 5% glycerol, 3 mM 2-Mercaptoethanol. Quality of CARM1 (>95%) was determined by SDS–PAGE. The protein was then concentrated, flash frozen with liquid nitrogen, stored at −80 °C for future use.

SPR analysis was performed using a Biacore™ T200 (GE Health Sciences Inc.) at 25 °C. Approximately 5,500 response units of CARM1 (amino acids 1-608) were amino-coupled onto a CM5 chip in one flow cell according to the protocol of the manufacturer. Another flow cell was left empty for reference subtraction. SPR analysis was conducted in the HBS-EP buffer (20 mM HEPES pH 7.4, 150 mM NaCl, 3 mM EDTA, 0.05% Tween-20) containing 2% (v/v) DMSO. The stock solutions of five concentrations of compound **2a** (24.7, 74.1, 222, 667 and 2,000 nM) and four concentrations of **5a** (6.2, 18.5, 55.6 and 167 nM) were prepared by serial dilution. Binding kinetic experiments were performed with single cycle kinetics with the contact time of 60 s and off time of 300 s in a flow rate of 30 µL/min. To facilitate complete dissociation of compound for the next cycle, a regeneration step (300 s, 40 µL/min of buffer), a period of stabilization (120 s) and two blank cycles were included between each cycle. Kinetic curve fittings were carried out with a 1:1 binding model or a heterogeneous ligand model using Biacore T200 Evaluation software (GE Health Sciences Inc.).

**2.5 *In vitro* thermal shift assay (TSA)**. TSA was performed as described previously(Blum et al., 2014; Niesen et al., 2007) to examine the melting temperature (*T*_m_) of CARM1 in the presence or absence of ligands. For each measurement (triplicate of each data point), the assay solution containing 50 mM HEPES-HCl (pH = 8.0), Tween-20 0.005% (v/v), 1 mM TCEP, 0.5 μM CARM1, and 5 μM ligand (SAM, **1**, **2a** or **5a**) was mixed with 5×SYPRO Orange Protein Gel Stain stock (Sigma Aldrich) in a 96-well PCR plate. The mixture was equilibrated at 25 °C in dark for 5 min and then loaded onto Bio-Rad CFX96 Real-Time PCR Detection System. The fluorescence readouts were recorded upon increasing the heating temperature from 25 °C to 100 °C at a rate of 0.2 °C/s. The raw data of the fluorescence readouts *versus* the temperatures were exported with CFX software and processed as the percentage of the fluorescent signal normalized between the lowest readout of 0% and the highest readout of 100% within the 25 ~100 °C region. The melting curves were plotted as the normalized fluorescence (%) versus the heating temperature and fit with a sigmoid curve with GraphPad Prism. The *T*_m_ corresponds to the temperature with the 50% relative fluorescent signal in the sigmoidal curve.

**2.6 Crystallization of CARM1 in complex with 1 and 5a.** A DNA fragment encoding the methyltransferase domain of human CARM1 (residues 140–480) was cloned into a baculovirus expression vector pFBOH-MHL (http://www.thesgc.org/sites/default/files/toronto_vectors/pFBOH-MHL.pdf). The protein was expressed in Sf9 cells as an N-terminal 6×His tag fusion protein and purified by a metal chelating affinity chromatography (TALON resin, Clontech, Mountain View, CA, USA) followed by size-exclusion chromatography (Superdex 200, GE Healthcare). Pooled fractions containing CARM1 were subjected to the treatment of tobacco etch virus to remove the 6×His tag. The protein was purified to homogeneity by ion-exchange chromatography. Purified CARM1 (6.5 mg/mL) was crystallized with the sitting drop vapor diffusion method at 20 °C.

For the CARM1-**1** complex, CARM1 was mixed with **1** at the 1:5 molar ratio (protein:ligand) and crystallized with the sitting drop vapor diffusion method at 20 °C by mixing 1 µL of the protein solution with 1 µL of the reservoir solution containing 20% PEG3350 and 0.2 M diammonium tartrate. X-ray diffraction data for the CARM1-**1** complex were collected at 100 K at beam line 23ID-B of Advanced Photon Source (APS), Argonne National Laboratory. Data sets were processed using the HKL-2000 suite.(Otwinowski and Minor, 1997) The structure of the CARM1-**1** complex was solved by molecular replacement using MOLREP(Otwinowski and Minor, 1997)with the PDB entry 2V74 as the search template. REFMAC(Emsley and Cowtan, 2004; Murshudov et al., 1997) was used for the structure refinement. Graphics program COOTS4 was used for model building and visualization. Crystal diffraction data and refinement statistics for the structure are displayed in Table S13. To further confirm the electronic densities of the ligand, the total omission electron density map was calculated using SFCHECK from CCP4suite and contoured at 1.0 s (Figure S6).(Emsley and Cowtan, 2004; Murshudov et al., 1997)

For the CARM1-**5a** complex, CARM1 was crystallized with the sitting drop vapor diffusion method at 20 °C by mixing 1 µL of protein solution with 1 µL of the reservoir solution containing 25% PEGG3350, 0.1M ammonium sulfate and 0.1 M Hepes (pH = 7.5). The compound **5a** (0.2 μL of 10 μM in DMSO) was added to the drops with apo crystals and incubated overnight. X-ray diffraction data for the CARM1-**5a** complex were collected at 100 K on a Rigaku FR-E superbright X-ray generator. Data were processed using the HKL-3000 suite.(Minor et al., 2006) The structure was isomorphous to PDB entry 4IKP, which was used as a starting model. REFMAC(Murshudov et al., 2011) was used for structure refinement. Geometric restraints for compound refinement were prepared with GRADE v.1.102 developed at Global Phasing Ltd. (Cambridge, UK). The COOT graphics program(Emsley et al., 2010) was used for model building and visualization, and MOLPROBITY(Williams et al., 2018) was used for structure validation. To further confirm the electronic densities of the ligand, the total omission electron density map was calculated using SFCHECK from CCP4suite and contoured at 1.0 s (Figure S6).(Emsley and Cowtan, 2004; Murshudov et al., 1997)

**2.7 Molecular docking and molecular dynamics simulations of CARM1-2a and CARM1-SNF complexes.** The ligands were docked into the binding site of CARM1 using the induced-fit docking (IFD) protocol(Sherman et al., 2006)implemented in the Schrodinger suite (release 2016-4). The poses for **SNF** and **2a** were selected according to the IFD scores. Specifically, the results identified two distinct poses for **2a** with similar scores but only one pose for **SNF**. The poses were then further relaxed by all-atom, explicit solvent molecular dynamics (MD) simulations. Herein CARM1 models in complex with the two ligands were placed into explicit water boxes. Simple point charge (SPC) water model(Berendsen et al., 1981) was used to solvate the system, charges were neutralized, and 0.15 M NaCl was added. The total system size was ∼50,000 atoms. Desmond MD systems (D. E. Shaw Research, New York, NY) with OPLS3 force field(Harder et al., 2016) were used. The system was initially minimized and equilibrated with restraints on the ligand heavy atoms and protein backbone atoms, followed by production runs with all atoms unrestrained. The isothermal-isobaric ensemble was used with constant temperature (310 K) maintained with Langevin dynamics and 1 atm constant pressure achieved with the hybrid Nose-Hoover Langevin piston method(Feller et al., 1995) on a flexible periodic cell. For each CARM1-ligand complex, a 600 ns trajectory was collected.

**2.8 LC-MS/MS quantification of intracellular concentrations of 6a, 5a, 2a and SAM**

**2.8.1 Sample preparation for LC-MS/MS analysis.** To measure the intracellular concentrations of **6a**, **5a**, **2a**, and SAM, a LC-MS/MS quantification method was developed with modification of what was reported previously.(Wang et al., 2014b) Briefly, 0.2×10^6^ MDA-MB-231 cells were incubated with varied concentrations of **6a** (0.5, 2.5, 5.0, and 10.0 µM) for several different periods of time (0.1, 3, 6, 12, 24, and 48 h). The treated cells (triplicate for each data point) were harvested and centrifuged at 4 °C at 94 g for 5 min. The cell pellets were resuspended and then washed with 4×1 mL of 4 °C PBS to remove extracellular **6a**. The 4×1 mL washing is sufficient to remove extracellular **6a**, as evidenced by the LC-MS/MS analysis that the residual **6a** in each washing buffer gradually decreased until the full loss of its MS signal after the 4 times of washing. The washed cell pellets were then treated with 40 µL MeOH (with 0.1% TFA, v/v) containing 0.125 µM **6b**, 0.125 µM **5b**, 16 µM **2b**, and 2 µM CD_3_-SAM(Linscott et al., 2016) as the LC-MS/MS internal standards. The mixture was lysed by 150 W sonication at 0 °C for 20 min. The resultant cell lysis was centrifuged with 21,130 g at 4 °C for 20 min. The 30 µL aliquant of the MeOH extraction of each sample was collected into a 96 well plate (5042-1386, Agilent) and stored at −20 °C until LC-MS/MS analysis.

**2.8.2 LC-MS/MS conditions.** Liquid chromatography-tandem mass spectrometry analysis was performed with a 6410 triple-quad LC-MS/MS system (Agilent Technologies) in electrospray ionization (ESI) mode, equipped with an Agilent Zorbax Eclipse XDB-C18 column (2.1 × 50 mm, 3.5 µM). The samples were eluted with a 5-95% gradient (v/v) of CH_3_CN in aqueous formic acid (HCOOH, 0.1%, v/v) in 7 min with a flow rate of 0.4 mL/min. The 96-well sample plate obtained above was maintained in the 4 °C chamber of the LC-MS/MS prior to analysis. The 7 µL MeOH extraction of each sample was injected into the LC-MS/MS and the MS signals were collected via the multiple-reaction monitoring (MRM) mode.

**2.8.3 Working curves of 6a, 5a, 2a, and SAM with 6b, 5b, 2b, and CD_3_-SAM as internal standards (Figure S8, S9).** Standard working curves to quantify **6a**, **5a**, **2a**, and SAM were generated by plotting a linear function (eq. S2) with the x-axial as the ratio of the mass peak areas (*P*_A_*/P*_IS_) and the y-axial as the ratio of the concentration (*C*_A_/*C*_IS_) between each analyte (*A*) and the structurally related internal standard (*IS*). To obtain the values of “*p*” and “*q*” in eq. S2, the 0.1% TFA MeOH (v/v) samples containing the varied concentrations of an analyte (**6a**, **5a**, **2a** or SAM) and the fixed concentrations of the mixture of the internal standards (0.125 µM **6b**, 0.125 µM **5b**, 16 µM **2b**, and 2 µM CD_3_-SAM) were subject to LC-MS/MS analysis. For **6a**, **5a**, and **2a**, three working curves for each analyte (the total of 9 for the three analytes) were generated to cover the concentration range of 3.9 nM~18.0 µM of the analyte (*C*_A,_**_6a_**, *C*_A,_**_5a_** and *C*_A,_**_2a_**) with the structurally related **6b**, **5b** and **2b** (*C*_IS,_**_6b_**, *C*_IS,_**_5b_** and *C*_IS,_**_2b_**) as internal standards. For SAM, one working curve was generated to cover the concentration range of 0.28~18 µM of SAM (*C*_A,SAM_) with CD_3_-SAM as the internal standard (*C*_IS,CD3-SAM_).

Y = *p* × X + *q*	eq. S2

**2.8.4 Quantification of intracellular concentrations of 6a, 5a, 2a, and SAM.** With the standard working curves generated above, the concentration (*C*_A_) of each analyte (**6a**, **5a**, **2a**, and **SAM**) in the MeOH extraction of cell lysates was obtained through the ratio (*P*_A_/*P*_IS_) of the mass peak areas of each analyte (*P*_A_) versus each internal standard (*P*_IS_), and the concentration of the internal standard (*C*_IS_) according to eq. S2. For **6a**, **5a**, and **2a** under each assay condition, three similar *C*_A_ values (*C*_A-_**_6b_***, C*_A-_**_5b_***,* and *C*_A-_**_2b_**) were obtained with **6b**, **5b** and **2b** as the internal standards, respectively. An average concentration of the analyte (*C̅_A_*) was obtained on the basis of the three concentrations weighted by the mass peak areas of the three internal standards (*P*_IS,_**_6b_**_,_ *P*_IS,_**_5b_** and *P*_IS,_**_2b_**) according to eq. S3:

$\text{C̅}\text{A} = \frac{\text{C}\text{A-}\text{6b}\text{ }\times\text{P}\text{IS-}\text{6b}}{\text{P}\text{IS-}\text{6b}+\text{P}\text{IS-}\text{5b}+\text{P}\text{IS-}\text{2b}\text{ }}+\frac{\text{C}\text{A-}\text{5b}\text{ }\times\text{P}\text{IS-}\text{5b}}{\text{P}\text{IS-}\text{6b}+\text{P}\text{IS-}\text{5b}+\text{P}\text{IS-}\text{2b}\text{ }}+ \frac{\text{C}\text{A-}\text{2b}\text{ }\times\text{P}\text{IS-}\text{2b}}{\text{P}\text{IS-}\text{6b}+\text{P}\text{IS-}\text{5b}+\text{P}\text{IS-}\text{2b}\text{ }}$ eq. S3

On the basis of the *C̅*_A_ values of **6a**, **5a**, and **2a** (*C̅*_A,_**_6a_**, *C̅*_A,_**_5a_** and *C̅*_A,_**_2a_**) in the MeOH extraction of cell lysates, the intracellular concentrations of the analyte **6a**, **5a**, and **2a** (*C*_intra,_**_6a_** *C*_intra,_**_5a_** and *C*_intra,_**_2a_**) were calculated according to eq. S4. Here *C̅*_A,_**_analyte_** is the weighted average of the three concentrations in the MeOH extraction, *N* is the cell number, and *V* is the mean volume of cells (µL). The mean volume of MDA-MB-231 cells is 1.3×10^-6^ µL/cell as reported previously.(Coulter et al., 2012) The cell number (N) was determined with a hemocytometer. The concentration of SAM (*C*_A,SAM_) in the MeOH extraction was obtained according to eq. S2, in which “X” is the ratio (*P*_A,SAM_/*P*_IS,CD3-SAM_) of the mass peak areas of SAM (*P*_A,SAM_) versus the internal standard CD_3_-SAM (*P*_IS,CD3-SAM_) and “Y” is the ratio (*C*_A,SAM_/*C*_IS,CD3-SAM_) of the concentrations of SAM versus CD_3_-SAM in the MeOH extraction. Given the identical LC-MS properties of SAM and CD_3_-SAM, *C*_A,SAM_ was obtained solely based on the working curve with CD_3_-SAM as the internal standard. The intracellular concentration of SAM (*C*_intra,SAM_) was then calculated using eq. S4, in which *N* is the cell number and *V* is the mean volume of MDA-MB-231 cells (1.3×10^-6^ µL/cell).

$\text{C}\text{intra,}\text{analyte}\text{ }=\frac{\text{C̅}\text{A,}\text{analyte}\text{ }\times40 \mu L}{N \times V}$ eq. S4

**2.8.5 Quantification of intracellular concentrations of 6b, 5b, 2b and SAM*.*** Similar experimental procedures were carried out to quantify the intracellular concentrations of **6b**, **5b**, **2b** and SAM except that **6b**, **5b**, and **2b** were the analytes and their structurally related **6a**, **5a** and **2a** were used as the internal standards with their *C*_IS_ values of 0.125 µM, 0.125 µM, and 0.0125 µM, respectively. For each analyte (**6b**, **5b**, and **2b**), three working curves (Figure S9) were generated to cover the concentration range of 3.9 nM~18.0 µM of the analyte (*C*_A,_**_6b_**, *C*_A,_**_5b_** and *C*_A,_**_2b_**) with **6a**, **5a** and **2a** (*C*_IS-_**_6a_**, *C*_IS-_**_5a_**, and *C*_IS-_**_2a_**) as the LC-MS/MS internal standards, respectively.

**2.9 Calculation of the percentages of intracellular apo-CARM1, CARM1 occupied by 6a, 5a, and 2a, and SAM-bond CARM1 versus the total amount of CARM1*.*** On the basis of the intracellular concentrations (*C*_intra,_ _analyte_) of **6a**, **5a**, **2a,** and SAM quantified by the LC-MS/MS experiment described above, the percentages of intracellular apo-CARM1, CARM1 occupied by **6a**, **5a**, and **2a**, SAM-bound CARM1 versus the total amount of intracellular CARM1 were calculated using eqs. S5-7, respectively.

$\text{Occupancy(Apo)\%=}\frac{\left[ \text{E} \right]}{\left[ \text{E} \right]\text{+}\left[ \text{E-SAM} \right]\text{+}\sum_{\text{i=1}}^{n} \text{[E-I}\text{i}\text{]}}\text{×100\% = }\frac{\text{1}}{\text{1+}\frac{\text{C}\text{intra, SAM}}{\text{K}\text{m,SAM}}\text{+}\sum_{\text{i}}^{\text{n}} \frac{\text{C}\text{intra, }\text{Ii}}{\text{K}\text{d,}\text{Ii}}}\text{×100\%}$	eq. S5

$\text{Occupancy(I)\%=}\frac{\sum_{\text{i=1}}^{\text{n}} \text{[E-I}\text{i}\text{]}}{\left[ \text{E} \right]\text{+}\left[ \text{E-SAM} \right]\text{+}\sum_{\text{i=1}}^{n} \text{[E-I}\text{i}\text{]}}\text{×100\% = }\frac{\sum_{\text{i}}^{\text{n}} \frac{\text{C}\text{intra, }\text{Ii}}{\text{K}\text{d,Ii}}}{\text{1+}\frac{\text{C}\text{intra, SAM}}{\text{K}\text{m,SAM}}\text{+}\sum_{\text{i}}^{\text{n}} \frac{\text{C}\text{intra, }\text{Ii}}{\text{K}\text{d,}\text{Ii}}}\text{×100\%}$	eq. S6

$\text{Occupancy(SAM)\%=}\frac{\left[ \text{E-SAM} \right]}{\left[ \text{E} \right]\text{+}\left[ \text{E-SAM} \right]\text{+}\sum_{\text{i=1}}^{n} \text{[E-I}\text{i}\text{]}}\text{×100\% = }\frac{\frac{\text{C}\text{intra, SAM}}{\text{K}\text{m,SAM}}}{\text{1+}\frac{\text{C}\text{intra, SAM}}{\text{K}\text{m,SAM}}\text{+}\sum_{\text{i}}^{\text{n}} \frac{\text{C}\text{intra, }\text{Ii}}{\text{K}\text{d,}\text{Ii}}}\text{×100\% }$	eq. S7

Here [E], [E-SAM] and [E-**I_i_**] are the intracellular concentrations of apo-CAMR1, the CARM1-SAM complex and the CARM1 occupied by the inhibitors; *C*_intra,SAM_ and *C*_intra,_**_Ii_** are the intracellular concentrations of SAM and individual CARM1 inhibitors, respectively; *K*_m,SAM_ is the Michaelis-Menton constant of SAM to form the CARM1-SAM complex, which is approximate to *K*_d,SAM_ (*K*_m,SAM_ ≈ *K*_d,SAM_ = 245 nM); *K*_d,_**_Ii_** is the dissociation constant of these CARM1 inhibitors. In the case of the cellular treatment with **6a**, “n” is equal to 3; I_1_, I_2_ and I_3_ stand for **2a**, **5a** and **6a**, respectively; *K*_d,_**_I1_**, *K*_d,_**_I2_** and *K*_d,_**_I3_** stand for *K*_d,_**_2a_** = 17 nM, *K*_d,_**_5a_** = 9 nM and *K*_d,_**_6a_** = 275 nM.

**2.10 Evaluation of methylation marks of CARM1 in breast cancer cells.** MDA-MB-231 cells and MCF-7 parental and CARM1 KO(Wang et al., 2014a) cells were maintained in DMEM (Gibco, Gaithersburg, MD) media containing 10% FBS (Gibco). These cells were then treated with compounds or DMSO for 48 h. The resultant cells were washed by phosphate-buffered saline (PBS) for 2 times and the samples were sonicated in ice-cold RIPA buffer (Thermo, Waltham, MA). The lysates were centrifuged (15,000 g) at 4 °C for 15 min. The supernatants were kept with the total protein amount determined with Bradford protein assay (BioRad, Hercules, CA). After quantification, 50 μg of protein from each sample was loaded onto 6% SDS-PAGE and transferred onto nitrocellulose membrane (PALL, Port Washington, NY). For the Arg1064 methylation mark of BAF155, the blots were blocked in 5% non-fat milk for 1 h and incubated with anti-me-BAF155 (Cancer Cell, 2014), anti-BAF155 (1:1000; Santa Cruz Biotechnology, Dallas, TX), anti-CARM1 (1:1000; Genemed Synthesis, San Antonio, TX), and anti-β-Actin (1:20000; Sigma-Aldrich, St. Louis, MO) overnight at 4 °C. After three times washing in Tris-buffered saline with Tween 20 (TBST), the blots were incubated with HRP-conjugated secondary antibody (1:3000; Jackson ImmunoResearch, West Grove, PA). After washing of blots with TBST, the membranes were developed using SuperSignal West Pico ECL solution (Thermo). For the Arg455/Arg460 methylation mark of PABP1, anti-me-PABP1 (1:1000) and anti-PABP1 (1:1000) antibodies (Genemed Synthesis, San Antonio, TX) were used instead. Here the antibodies against CARM1, PABP1, and me-PABP1 were custom generated by Genemed Synthesis (San Antonio, TX).(Zeng et al., 2013) The density of the protein bands was quantified using ImageJ software (NIH, Bethesda, MD). The EC_50_ values were obtained by fitting methylation percentage (%) of BAF155 or PABP1 against the concentrations of the inhibitor using a sigmoidal equation with GraphPad Prism Software.

**2.11 Cellular thermal shift assay (CETSA).** CETSA was performed as described previously(Jafari et al., 2014) to examine the intracellular engagement of **6a** or **6b** with CARM1. Briefly, 2.0×10^6^ MDA-MB-231 cells were incubated with 15 μM **6a** or **6b** and DMSO for 48 h. The harvested cells were re-suspended in PBS buffer and divided into eight aliquots (30 µL/aliquot), and the heat-shocked at various temperature (49.1, 54.6, 57.0, 59.5, 61.8, 63.9, 65.6, and 67.0 °C) for 3 min with a Bio-Rad CFX96 Real-Time PCR Detection instrument (the temperature gradient of 49~67 °C). The heat-shocked cells were then lysed with the freeze-thaw method with a liquid nitrogen bath followed by a 25 °C water bath for five cycles. The cell lysate was centrifuged at 4 °C at 18,000 g for 20 min. The resultant supernatant containing the soluble protein fraction was collected and loaded on SDS-PAGE gel (20 µL). Western blotting of CARM1 was performed with anti-CARM1 antibody (Cell Signaling Technology, C31G9). For each sample, the band intensity of CARM1 was quantified with ImageJ. The band intensity of CARM1 was normalized to the band intensity at 49.1°C (the lowest heat-shock temperature). Melting curves were obtained by plotting the normalized band intensity versus the heat-shock temperatures and fit with a Boltzmann sigmoidal equation in GraphPad Prism. The melting temperatures (*T*_m_) of **6a**, **6b**, or DMSO were obtained as the heat-shock temperatures that correspond to the 50% normalized band intensity in the fitted sigmoidal curves.

**2.12 Cell invasion and proliferation assay.** MDA-MB-231 parental and CARM1 *KO* cells(Wang et al., 2014a) were maintained in DMEM (Gibco, Gaithersburg, MD) media containing 10% FBS (Gibco). Cell invasion assays were performed using 8.0 µm pore size Transwell inserts (Greiner Bio-One, Kremsmünster, Austria). MDA-MB-231 parental and CARM1 *KO* (Cancer Cell, 2014) cells were harvested with trypsin/EDTA and washed twice with serum-free DMEM (Gibco). 2×10^5^ cells in 0.2 mL serum-free DMEM (Gibco) were seeded to the upper chamber, which was pre-coated with a thin layer of 40 uL of 2 mg/mL Matrigel (Corning, NY, USA) for 2 h incubation at 37 °C incubator. To the lower chamber was added 0.6 mL DMEM containing 10% FBS (GIBCO) and compounds or DMSO. After 16 h in 37 °C incubator, the cells on the inner side of the upper chamber together with the Matrigel layer were removed using cotton tips. The residual invasive cells in the outer side of the upper chamber were fixed in 3.7 % formaldehyde (a weight percentage) at ambient temperature (22 °C) for 2 min, 100% methanol for 20 min and stained for 15 min with a solution containing 1% crystal violet and 2% ethanol in 100 mM borate buffer (pH 9.0). The number of invasive cells was counted under microscope by taking five independent fields. Relative cell invasion was determined by the number of the invasive cells normalized to the total number of the cells adhering to 0.8 µm transwell filters. The EC_50_ was obtained with GraphPad Prism Software upon fitting eq. S8, in which “Inhibition%” is the percentage of the inhibition of invasiveness, “Maximal Inhibition%” is the percentage of the maximal inhibition of invasiveness, [Inhibitor] is the concentration of the inhibitor.

$\text{Inhibition\%=}\frac{\text{Maximal Inhibition\%×[Inhibitor]}}{\left[ \text{Inhibitor} \right]\text{ + EC}_{\text{50}}}$	eq. S8

To examine proliferation of MDA-MB-231 cells, 5000 cells of parental and CARM1 *KO* cells were seeded at a 96-well plate and incubated in 37 °C for overnight. These cells were treated with various doses (0.0001~10 μM) of **SKI-73** or **SKI-73N** in DMSO and incubated for 72 hours. MTT assay was then performed to examine viability with DMSO-treated cells as the control. The relative viability of compound-treated cells versus DMSO-treated parent cells were plotted against the concentrations of **SKI-73** and **SKI-73N**.

**12.13 Sample preparation for single cell RNA-seq analysis (scRNA-seq)**

**2.13.1** Cell preparation for scRNA-seq. 10× Genomics droplet-based scRNA-seq was implemented to characterize five types of MDA-MB-231 cells: MDA-MB-231 cells treated with **DMSO**, **SKI-73** and **SKI-73N**, MDA-MB-231 cells that freshly invaded through Matrigel, and CARM1-*KO* MDA-MB-231 cells:

For the first three samples, MDA-MB-231 cells were treated with DMSO, 10 μM **SKI-73**, and 10 μM **SKI-73N** for 48 hours. The resulting cells were trypsinized at 37 °C for 3 min. The harvested cells were washed twice with 1×PBS (phosphate-buffered saline) containing 0.04 % (volume%) bovine serum albumin (BSA), gently dispersed to dissociate cells, and then filtered through a cell strainer (100 μm Nylon mesh, Fisherbrand) to obtain single cell suspensions for scRNA-seq.

For the CARM1-*KO* sample, the cells were trypsinized at 37 °C for 3 min, washed twice with 1×PBS containing 0.04 % BSA, and filtered through a 100 μm Nylon-mesh cell strainer (Fisherbrand) to obtain single-cell suspensions for scRNA-seq.

To collect the cells that freshly invaded through Matrigel, the conditions described above for cell invasion assay were applied to allow approximately 5% of the 1×10^7^ seeded MDA-MB-231 cells to invade through Matrigel. Briefly, 10 Transwell inserts with a diameter of 24 mm and a pore size of 8.0 μm were pre-coated with a thin layer of 54 μL of 2 mg/mL Matrigel (Corning). Into the upper chamber of each Matrigel-coated Transwell insert were seeded 1×10^6^ MDA-MB-231 cells in 1.0 mL serum-free DMEM (Gibco). Into the lower chambers was added 2.0 ml DMEM containing 10% FBS (Gibco). After 16-hour incubation, the cells on the inner side of the upper chamber together with the Matrigel layer were removed using cotton tips. The cells that freshly invaded---those attached on the outer side of the upper chamber, were subject to 3-min trypsin digestion at 37 °C for detachment. The resulting cells were washed twice with 1×PBS containing 0.04 % BSA, gently dispersed to dissociate cells, and filtered through a 100 μm Nylon-mesh cell strainer (Fisherbrand) to obtain single-cell suspensions for scRNA-seq.

**2.13.2 Cell barcoding, library preparation and sequencing.** The scRNA-seq libraries were prepared following the user guide manual (CG00052 Rev E) provided by the 10× Genomics and Chromium™ Single Cell 3' Reagent Kit (v2). Briefly, samples containing approximately 8,700 cells (93–97% viability) were encapsulated in microfluidic droplets at a dilution of 66-70 cells/µL, which resulted in 4,369-5,457 recovered single-cells per sample with a multiplet rate ~3.9%. The resultant emulsion droplets were then broken and barcoded-cDNA was purified with DynaBeads, followed by 12-cycles of PCR-amplification---98 °C for 180 s, 12×(98 °C for 15 s, 67 °C for 20 s, 72 °C for 60 s), and 72 °C for 60 s. The 50 ng of PCR-amplified barcoded-cDNA was fragmented with the reagents provided in the kit and purified by SPRI beads with an averaged fragment size of 600 bp. The DNA library was then ligated to the sequencing adapter followed by indexing PCR---98 °C for 45 s; 12× (98 °C for 20 s, 54 °C for 30 s, 72 °C for 20 s), and 72 °C for 60 s. The resulting DNA library was double-size purified (0.6-0.8×) with SPRI beads and sequenced on Illumina NovaSeq platform (R1 – 26 cycles, i7 – 8 cycles, R2 – 96 cycles) resulting in 70-79 million reads per sample with average reads per single-cell being 8,075-10,342 and average reads per transcript 1.11-1.15.

**2.14 Processing, transformation, filtering and dimensionality reduction of scRNA-seq data.** The fastq files containing the transcriptome and barcoding metadata were demultiplexed using the SEquence Quality Control (SEQC) pipeline (http:github.com/ambrosejcarr/seqc.git) resulting in around 8000 UMIs per one cell. The table of UMI counts was used as the input and Seurat package v.2.3.4(Butler et al., 2018) was applied for scRNA-seq analysis. Here the raw UMI counts were normalized per cell by dividing the total number of UMIs in each individual cell, multiplying by a scale factor of 10,000 and transforming into natural logarithm values. Cells with 1,000~5,000 genes and < 20% mitochondrial RNA transcripts were kept for further analysis. Dimensionality reduction was carried out by selecting a set of highly variable genes on the basis of the average expression and dispersion per gene. The set of genes was used for principle component analysis (PCA). Top principle components were then chosen for cell clustering analysis and *t*-SNE projection. Regression was performed to remove cell-cell variation in gene expression driven by the UMI number, mitochondrial gene content and ribosomal gene content using “ScaleData” function in Seurat package.(Nestorowa et al., 2016) Clusters of cells were identified by a shared nearest neighbor (SNN) modularity optimization based clustering algorithm that was included in Seurat package v.2.3.4.(Butler et al., 2018)

**2.15 Subpopulation clustering guided by scRNA-seq**

**2.15.1 Cell cycle awareness.** To assign cells with cell cycle stages---G0/G1, S and G2/M, individual cells were scored on the basis of their expression of G2/M-phase and S-phase markers(Nestorowa et al., 2016) by comparing the average expression of these markers with that of a random set of background genes.(Tirosh et al., 2016) Cells with positive higher S-phase or G2/M-phase scores were assigned as S-phase or G2/M-phase cells, respectively. Cells with negative S-phase and G2/M-phase scores were assigned as non-S/G2/M-phase cells and annotated as G0/G1-phase cells. The whole cell population as well as its subpopulations can thus be classified into the three groups according to their cell-cycle scores.

**2.15.2 Determination of the number of clusters.** Three algorithms---Silhouette analysis, the entropy scoring, and Fisher’s Exact Test---were applied to collectively determine the number of clusters.

**Silhouette analysis.** Silhouette analysis was conducted with no awareness of cell origins (DMSO-, **SKI-73**- or **SKI-73N**-treated cells) and was calculated based on distances defined as Euclidean distance between any pair of cells on the two-dimensional *t-*SNE projection.(de Amorim and Hennig, 2015; Rousseeuw, 1987)

**The entropy scoring**. We developed an entropy-based scoring method to evaluate the efficiency of clustering subpopulations with the three cell origins (DMSO-, **SKI-73**-, and **SKI-73N**-treated). Entropy Score is defined with the range of 0~1 by eq. S9, which was derived on the basis of the double-weighted sum of cell-origin-based Shannon entropy across clustered subpopulations. Herein “*i*” is the series number of a clustered subpopulation starting from zero; “*n*” is the largest series number of clustered subpopulations; the total number of subpopulation is “*n+1*” (*i* = 0~n); “*f_i_*” corresponds to the fraction of the subpopulation “*i*” in the total cell population ($0<f_{i}\leq1; \sum_{i=0}^{n} f_{i}=1$); “*j*” represents one of the three cell origins (*j* =1, 2 or 3*)*; “*d_j,i_*” (“*d_DMSO,i_*”, “*d_SKI-73N,i_*” and “*d_SKI-73,i_*”) is the fractional distribution of the cells with the “*j*” origin (DMSO, **SKI-73**, and **SKI-73N**) within the “*i*” subpopulation ($0\leq d_{j,i}\leq1; \sum_{j=1}^{3} d_{j,i}=1$); “*f_j_*” is the fraction of the cells with the “*j*” origin within the total population ($0<f_{j}\leq1; \sum_{j=1}^{3} f_{j}=1$); “$-\sum_{j=1}^{3} (f_{j}\times\log_{e} {(f}_{j}))$” is the theoretical maximum of $(-\sum_{i=0}^{n} {(f}_{i}\times(\sum_{j=1}^{3} {(d}_{j,i}\times\log_{e} {(d}_{j,i}))))$---all the cells in a single cluster. A smaller Entropy Score indicates that the corresponding method allows cell subpopulations to be clustered with higher resolution for the DMSO-, **SKI-73**-, and **SKI-73N**-treated cell origins. The minimal Entropy Score of zero indicates that subpopulations can be fully resolved for the three treatment conditions.

Entropy Score = $(-\sum_{i=0}^{n} {(f}_{i}\times(\sum_{j=1}^{3} {(d}_{j,i}\times\log_{e} {(d}_{j,i}))))/(-\sum_{j=1}^{3} (f_{j}\times\log_{e} {(f}_{j}))$	eq. S9

**Fisher’s Exact Test.** Fisher’s Exact Test was implemented to evaluate the agreement of the clusters with the three cell origins (DMSO-, **SKI-73**- or **SKI-73N**-treated cells) using R package “fisher.test”: <http://mathworld.wolfram.com/FishersExactTest.html>.(Mehta and Patel, 1983) Because of significant computation cost of Fisher’s Exact Test, the cell population of each Fisher’s Exact Test was down-sampled to 150 cells and this process was repeated for 100 times to cover a majority of the cell population. The p-value of Fisher’s Exact Test was then computed by Monte Carlo simulation. Means and standard errors of p-values were calculated and reported as the outputs of Fisher’s Exact Test.

Three algorithmic scoring systems were used over a range of the resolution parameter that sets for the corresponding “granularity” of clustering, with higher values indicating a greater number of clusters. Here Silhouette analysis was applied to determine the number of clusters in an unsupervised manner---without awareness of the three cell origins (DMSO-, **SKI-73**- or **SKI-73N**-treated cells). The entropy scoring and Fisher’s Exact Test were implemented to evaluate the biological meaning of the clustering---using the minimal cluster number to maximally resolve the cells between the three treatment conditions. Given the awareness of the three cell origins, the minimal number of clusters with the maximal resolution of cell origin guided by the entropy scoring and Fisher’s Exact Test is expected within the 1~3-fold range of the optimized number of clusters guided by Silhouette analysis. Fisher’s Exact Test was used as the primary scoring method to determine the efficiency of clustering, given its higher resolution.

**2.16 Correlation analysis of subpopulations, heat map analysis, and selection of representative transcripts.**

**2.16.1 Population analysis of the three treatment conditions.** “*d_j,i_*” (“*d_DMSO,i_*”, “*d_SKI-73N,i_*” and “*d_SKI-73,i_*” in eq. 9 is defined as the fraction of the cells with the “*j*” origin (*j* =1, 2 or 3 for the treatment with DMSO, **SKI-73N** and **SKI-73**, respectively) within the “*i*” subpopulation (*i* = 0~n for the clustered subpopulation). “*d_i,total_*” is defined as the fraction of the cells of the “*i*” subpopulation within the total cell population in each cell-cycle stage. “*d_(j,i),total_* = *d_j,i_* × *d_i,total_*” represents the fraction of the cells with the “*j*” origin (DMSO, **SKI-73**, and **SKI-73N**) and the “*i*” subpopulation within the total cell population in each cell-cycle stage. For a specific subpopulation “*i*”, there are three “*d_(j,i),total_*” values---*d_(DMSO,i),total_*, *d_(SKI-73N,i),total_* and *d_(SKI-73,i),total_*---for the treatment with DMSO, **SKI-73N** and **SKI-73**, respectively. The alteration of subpopulations is defined as the ratios of *d_(SKI-73N,i),total_*/*d_(DMSO,i),total_* or *d_(SKI-73,i),total_*/*d_(DMSO,i),total_* fall out of the range of 1.0 ± 0.2. Population analysis was conducted by classifying the **SKI-73N**/**SKI-73**-treated subpopulations as the following five categories: commonly resistant (0.8 < “*d_(SKI-73N,i),total_*/*d_(DMSO,i),total_*” and “*d_(SKI-73,i),total_*/*d_(DMSO,i),total_*” < 1.2), commonly emerging (“*d_(SKI-73N,i),total_*/*d_(DMSO,i),total_*” and “*d_(SKI-73,i),total_*/*d_(DMSO,i),total_*” ≥ 1.2), commonly depleted (“*d_(SKI-73N,i),total_*/*d_(DMSO,i),total_*” and “*d_(SKI-73,i),total_*/*d_(DMSO,i),total_*” ≤ 0.8; ⏐“*d_(SKI-73N,i),total_*/*d_(DMSO,i),total_*”−“*d_(SKI-73,i),total_*/*d_(DMSO,i),total_*”⏐ < 0.15), differentially emerging (either “*d_(SKI-73N,i),total_*/*d_(DMSO,i),total_*” or “*d_(SKI-73,i),total_*/*d_(DMSO,i),total_*” > 1.2), and differentially depleted (“*d_(SKI-73N,i),total_*/*d_(DMSO,i),total_*” or “*d_(SKI-73,i),total_*/*d_(DMSO,i),total_*” < 0.8; ⏐“*d_(SKI-73N,i),total_*/*d_(DMSO,i),total_*”−“*d_(SKI-73,i),total_*/*d_(DMSO,i),total_*”⏐ > 0.15). While additional combinations of differentially altered subpopulations could be possible, only those defined above were found under our treatment conditions.

**2.16.2 Correlation analysis of subpopulations.** In each cell cycle (G0/G1, S and G2/M) of the cells treated with DMSO, **SKI-73** or **SKI-73N** and “invasion cells”, correlation analysis of subpopulations was conducted with “BuildClusterTree” function in Seurat package(<https://rdrr.io/cran/Seurat/man/BuildClusterTree.html>).(Nestorowa et al., 2016) The phylogenetic trees were constructed by averaging gene expressions across all cells in each subpopulation and then calculating distance on the basis of averaged expressions between different subpopulations.

**2.16.3 Differential expression across remotely related subpopulations and the selection of representative transcripts.** Differentially expressed genes were identified by comparing two groups of cells using Wilcox rank sumtest with “FindMarkers” function in Seurat package.(Nestorowa et al., 2016) In particular, the “invasion cells” and its most correlated clusters---revealed in the correlation analysis—0were selected as the “high” group; the remaining remotely related clusters were selected as the “low” group. The differential expression analysis was then performed by comparing cells in the “high” group with the cells in the “low” group. For the G0/G1-phase cells, the “invasion cells” and Subpopulation-6, 7, 8, 9, 14 were selected as the “high” group; Subpopulation-0, 1, 2, 3, 4, 5, 10, 11, 12, 13, 15, 16, 17, 18, 19, 20 were selected as the “low” group. For the G2M-phase cells, the “invasion cells” and Subpopulation-1, 2 were selected as the “high” group; Subpopulation-0, 3, 4, 5 were selected as the “low” group. For the S-phase cells, the “invasion cells” and Subpopulation-0, 3 were selected as the “high” group; Subpopulation-1, 2, 4, 5, 6 were selected as the “low” group. Differentially expressed genes were ranked according to the average log_2_ fold change “avg_logFC” and adjusted p-values “p_val_adj” with the Seurat package. Top up-regulated and down-regulated genes were chosen by setting a cutoff on their “avg_logFC” values (> 0.25 or <−0.25), and curating genes with potential functional relevance to cancer malignancy (30 upregulated and 10 downregulated genes) were then selected as representative genes for generating heat map plots with “DoHeatmap” function in Seurat package.

**2.16.4 Analysis of differential expression across invasion-prone subpopulation candidates and the selection of representative transcripts for Violin plots.** To select candidate genes of the G0/G1-phase cells in violin plots, the “invasion cells” and the cells of Subpopulation 8 were selected as the “high” group and the clusters 6, 7, 9, 14 were selected as the “low” group. Differentially expressed genes were ranked according to the average log_2_ fold change “avg_logFC” and adjusted p-values “p_val_adj”. Top up- and down-regulated genes were chosen by setting their “avg_logFC” values > 0.25 or < −0.25 and curating genes with functionally implicated in cancer malignancy. Heat map plots were generated with the selected gene with “DoHeatmap” function in Seurat package.(Nestorowa et al., 2016) Furthermore, a panel of top 8 genes highlighting similarity between the invasion-prone Subpopulation 8 and “invasion cells” (the top 5 upregulated and top 3 downregulated genes) were selected for generating violin plots.

**Supplementary Figures and Tables**.

Supplementary Figures S1-55.

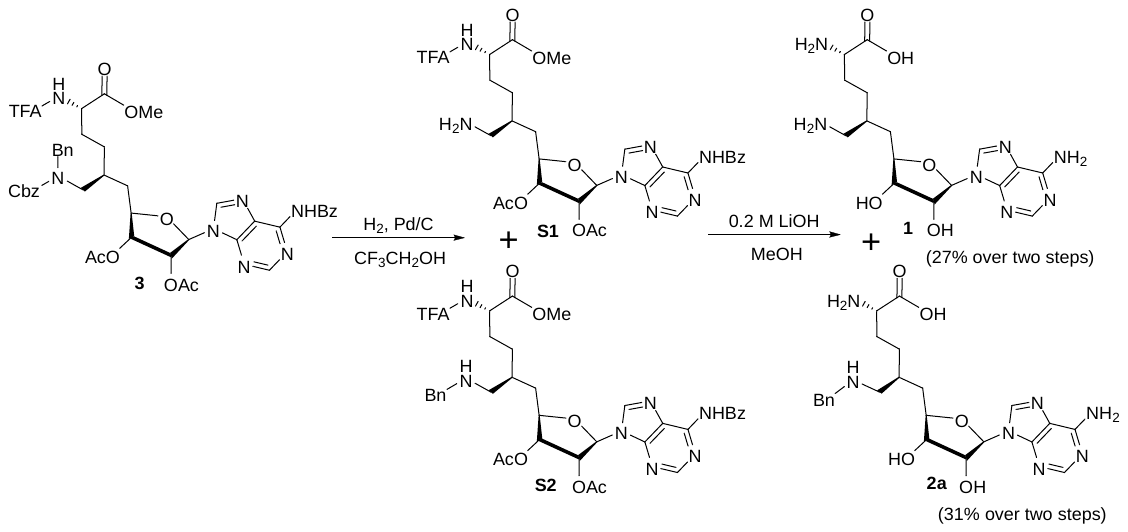

**Figure S1**. Synthetic scheme of **1** and **2a** through the precursor **3**.

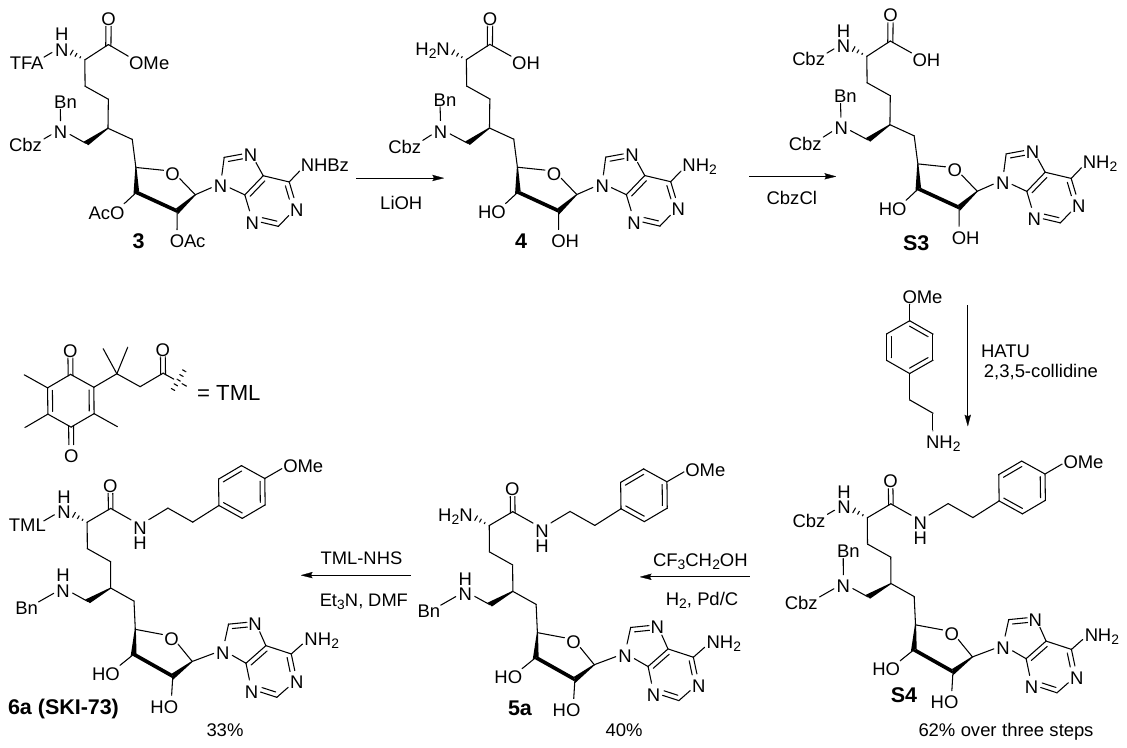

**Figure S2**. Synthetic scheme of **5a** and **6a (SKI-73)** through the precursor **3**.

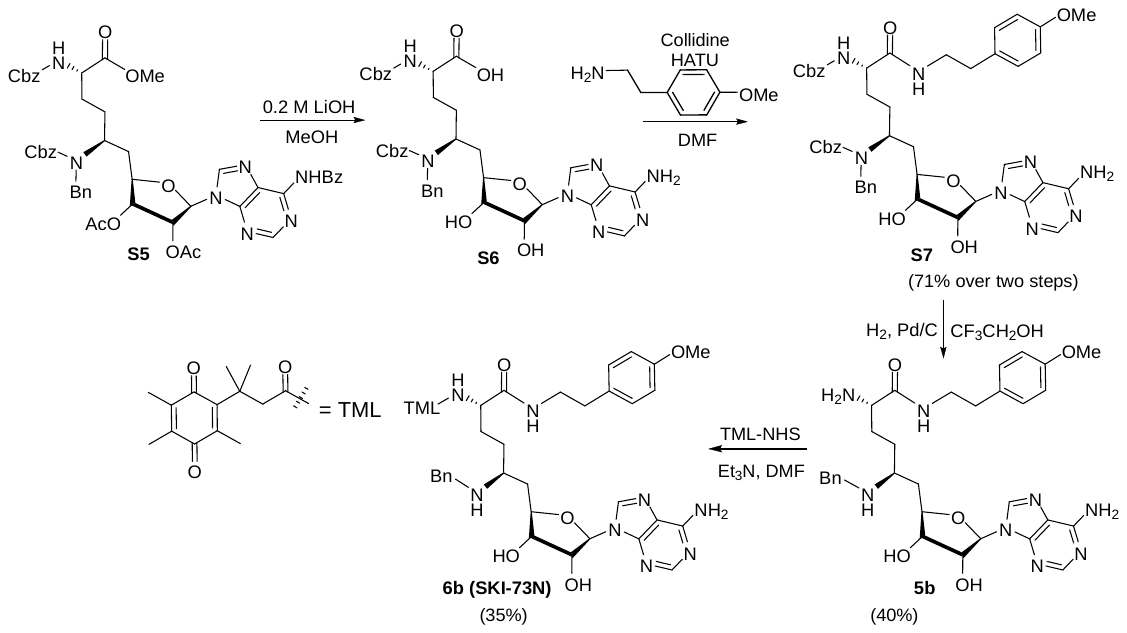

**Figure S3**. Synthetic scheme of **5b** and **6b (SKI-73N)**.

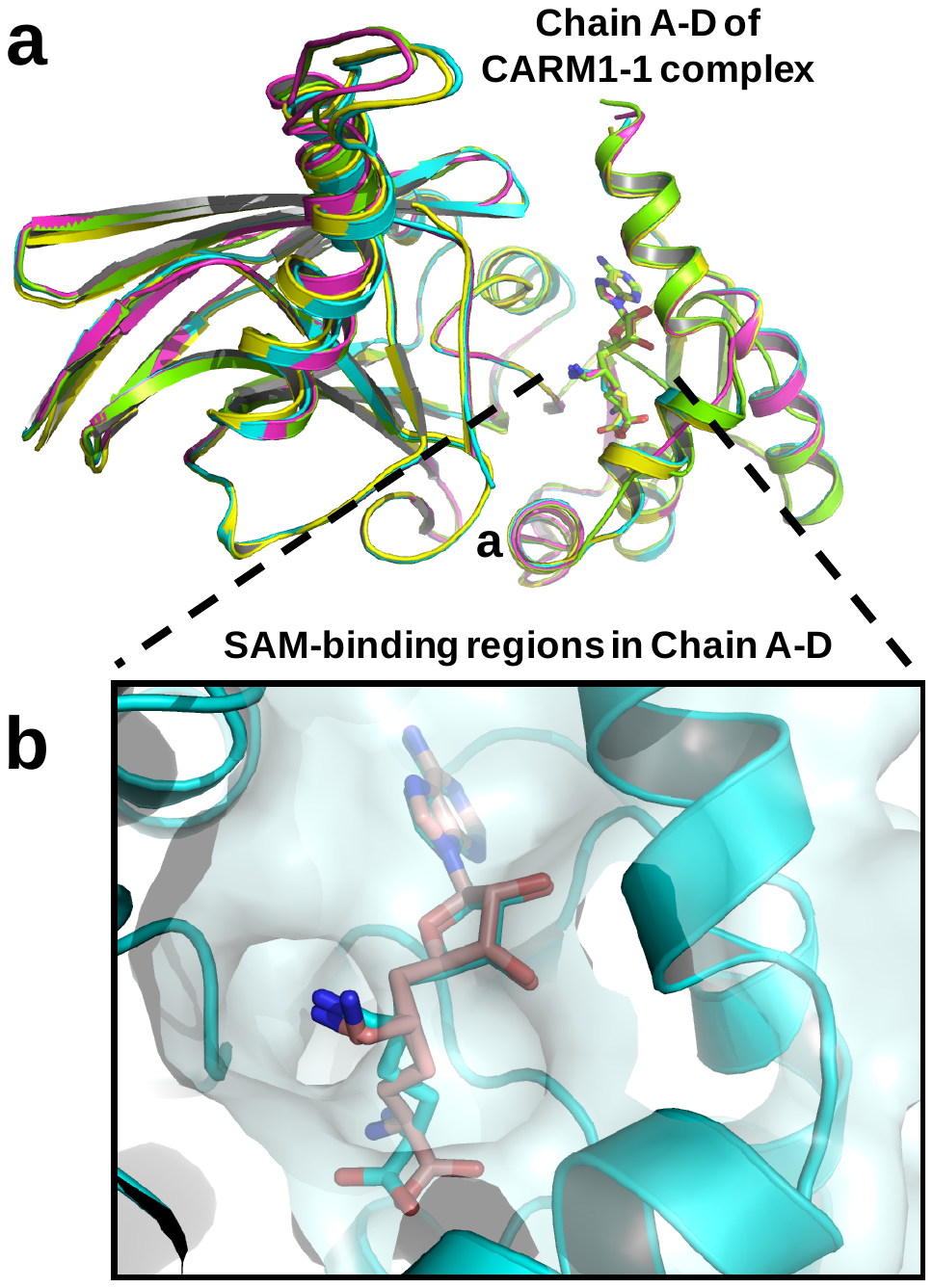

**Figure S4.** Structure of CARM1 in complex with **1** (HSF) in multiple configurations (PDB: 4IKP): (a) Overall structures of the CARM1-**1** complex featured by V-shape subunits in a tetramer (Chain A-D); (b) Multiple configurations of **1** upon occupying SAM-cofactor-binding of CARM1.

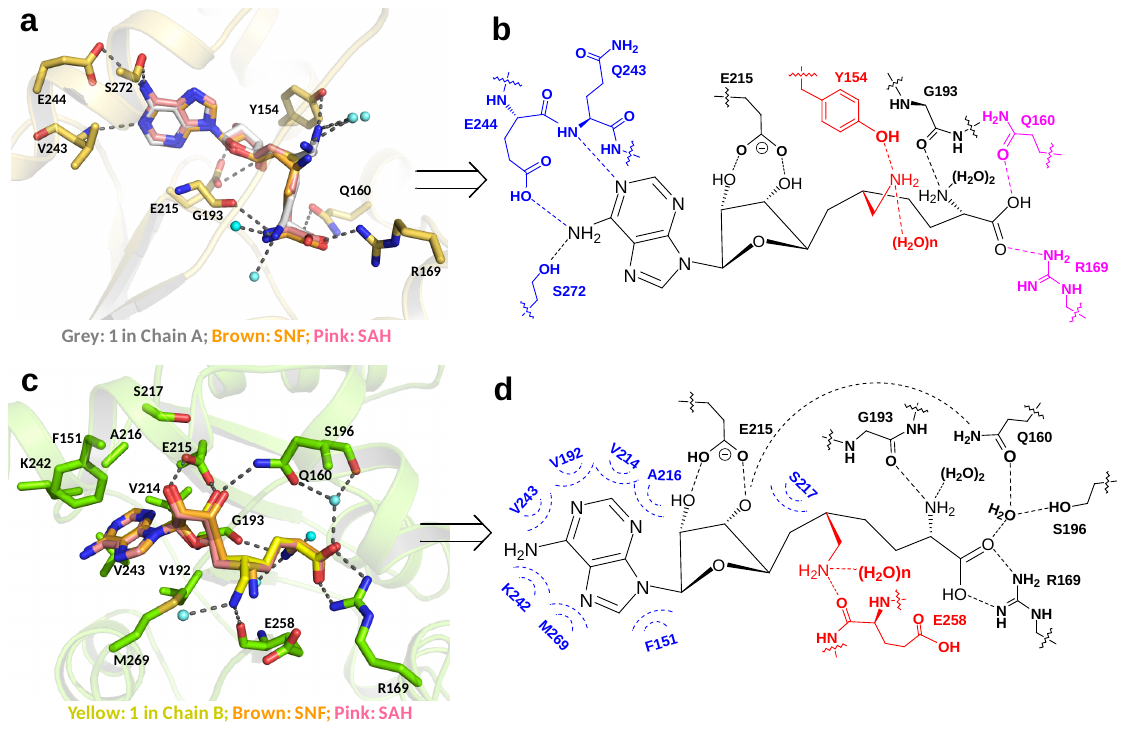

**Figure S5.** Structural comparison of CARM1 in complex with **1** (**HSF**), **SNF** and **SAH**: (a) Representative interaction network of **1** (Configuration III, IV in Chain A) upon binding human CARM1 and its comparison with **SNF** and **SAH**. Here highlighted are the conserved hydrogen bonds with adenine ring, 2′,3′-ribosyl hydroxyl, α-amino groups, and the distinct interaction network in Chain A for carboxylic and 6′-methyleneamine moieties; (b) Simplified version of Figure S5a with highlighted key interactions of **1** including hydrogen bonds with adenine ring (blue), 2′,3′-ribosyl hydroxyl/α-amino groups (black), carboxylic moieties/6′-methyleneamine (pink and red); (c) Representative interaction network of **1** (Configuration I in Chain B) upon binding human CARM1 and its comparison with **SNF** and **SAH**. Here highlighted are the conserved hydrophobic interactions with adenine ring, the conserved hydrogen bond interactions with 2′,3′-ribosyl hydroxyl, α-amino, carboxylic moieties as well as the distinct interaction network of 6′-methyleneamine; (d) Simplified version of Figure S5c with highlighted key interactions of **1** including interactions with adenine ring (blue), 2′,3′-ribosyl hydroxyl/α-amino/carboxylic moieties (black) and 6′-methyleneamine (red). The images of **SNF** and **SAH** were generated on the basis of PDB files 2Y1W and 2Y1X.

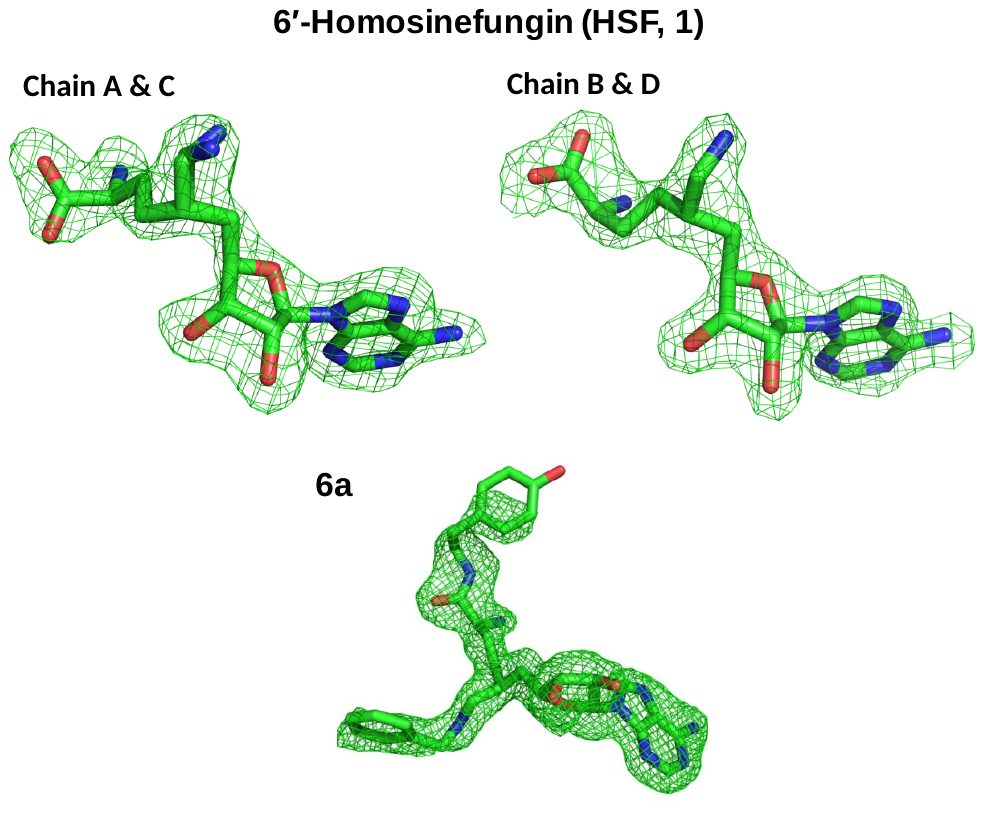

**Figure S6.** Total omission electron density map of 6′-homosinefungin (HSF, **1**) in the CARM1-**1** complex and **5a** in the CARM1-**5a** complex. The total omission electron density map was calculated using SFCHECK as describe in Supplementary Methods. The electron density contoured at 1.0 σ is shown for the ligands.

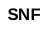

**Figure S7.** Conformational dynamics of CARM1-**2a** and CARM1-**SNF** complexes. Multiple configurations of Arg168, Tyr261 and His414 and their kinetics were revealed through 600 ns MD simulation for CARM1-**2a** and CARM1-**SNF** complexes.

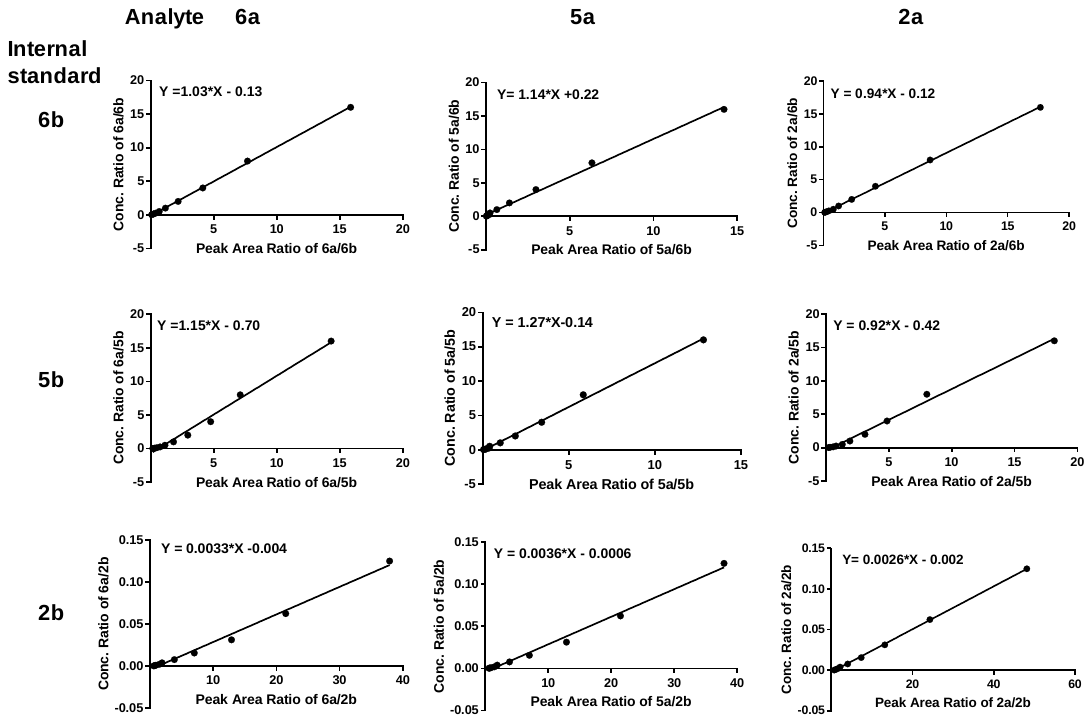

**Figure S8.** LC-MS/MS working curves for quantification of the analytes **6a**, **5a** and **2a** with **6b**, **5b**, and **2b** as internal standards. The concentration ratios show a linear correlation with the mass peak area ratios between the analyte and the internal standards.

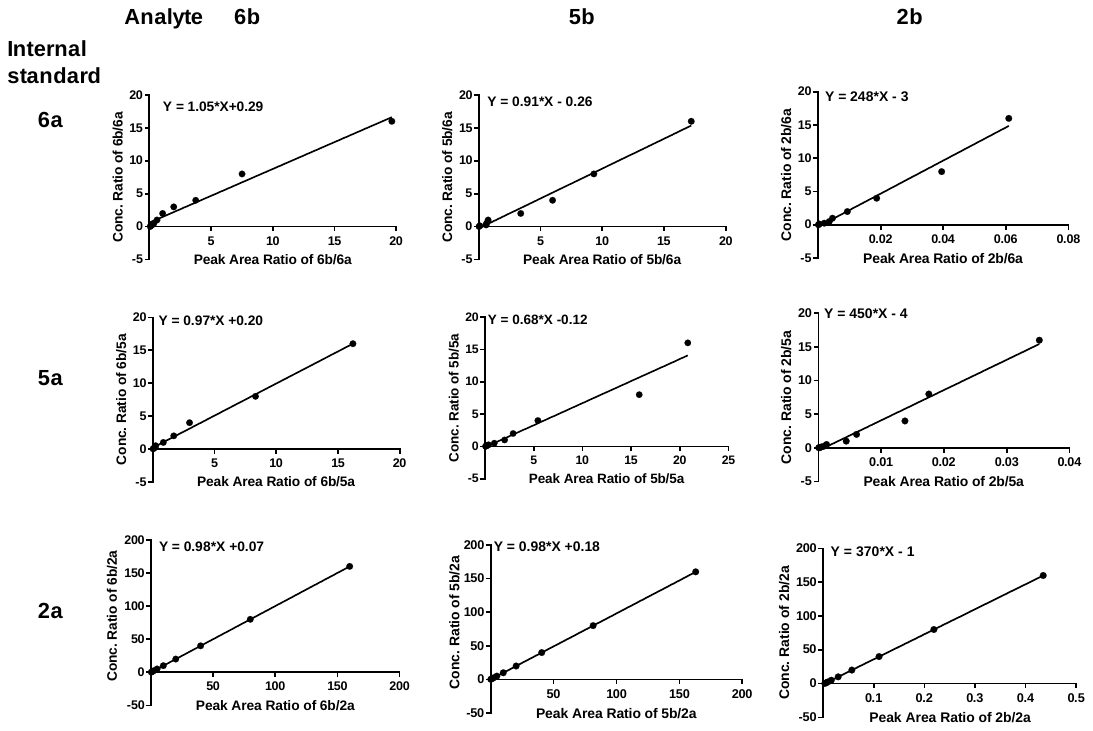

**Figure S9.** LC-MS/MS working curves for quantification of the analytes **6b**, **5b** and **2b** with **6a**, **5a**, and **2a** as internal standards. The concentration ratios show a linear correlation with the mass peak area ratios between the analyte and the internal standards.

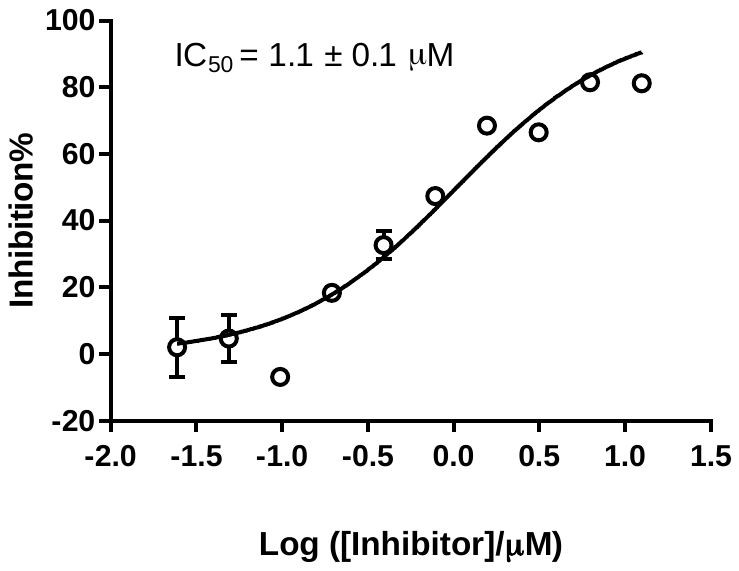

**Figure S10.** IC_50_ of **6a** against CARM1. The IC_50_ value was obtained by fitting inhibition% against the concentration of the inhibitor to a sigmoid curve with GraphPad Prism. Each data point represents the mean of two replicates and s.d.

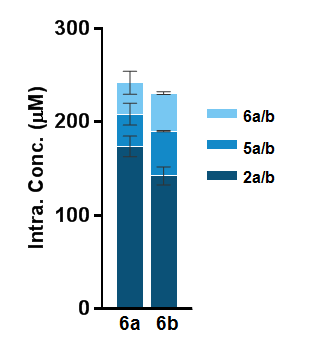

**Figure S11.** Cellular uptake and intracellular fate of **6b** and **6b**. MDA-MB-231 cells were incubated with 10 μM **6b** and **6a** for 24 hours. LC-MS/MS was used to quantify the intracellular concentrations of (**6b**, **5b**, **2b**) and (**6a**, **5a**, **2a**), respectively (n = 3, mean values ± s.d.). (**6b**, **5b**, **2b**) were accumulated inside MDA-MB-231 cells with the efficiency comparable to that of (**6a**, **5a**, **2a**).

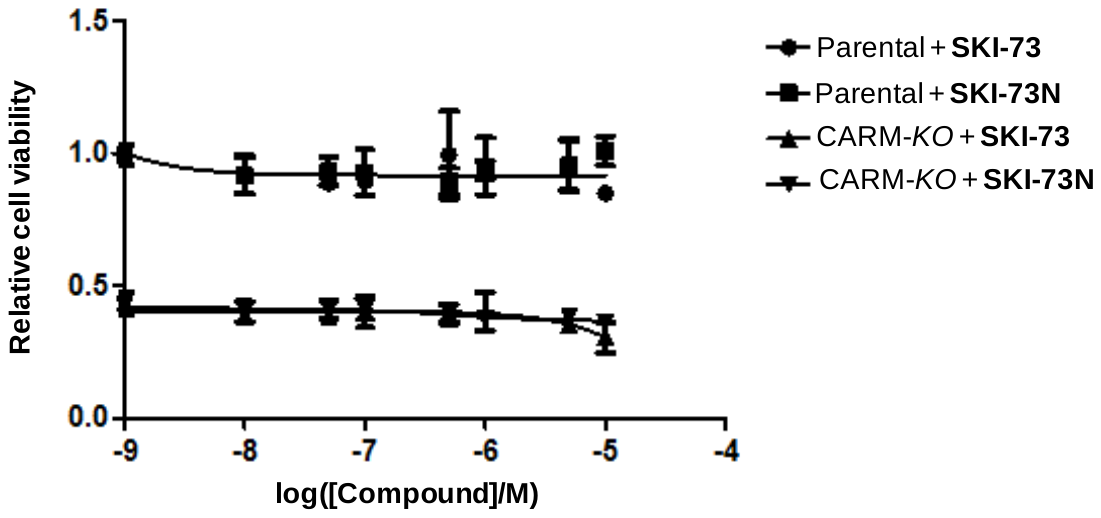

**Figure S12.** Viability of parental and CARM1-*KO* MDA-MB-231 cells upon the treatment of **SKI-73** and its control compound **SKI-73N**. The cells were treated with 0.0001~10 μM of **SKI-73** or **SKI-73N** for 72 hours. MTT assay was then performed to examine their relative viability with DMSO-treated parental cells as the reference.

**Figure S13.** Quality control of the cells subject for scRNA-seq analysis. MDA-MB-231 cells treated with **SKI-73N**, DMSO and **SKI-73**, invasion cells---MDA-MB-231 cells that freshly invaded through Matrigel, and CARM1-*KO* MDA-MB-231 cells were collected for 10× Genomics scRNA-seq. The threshold with 1,000~5,000 genes and < 20% mitochondrial RNA transcripts for further analysis were used to select the cells with their UMIs (top), the number of genes per cell (middle) and the fraction of mitochondria genes (bottom) shown.

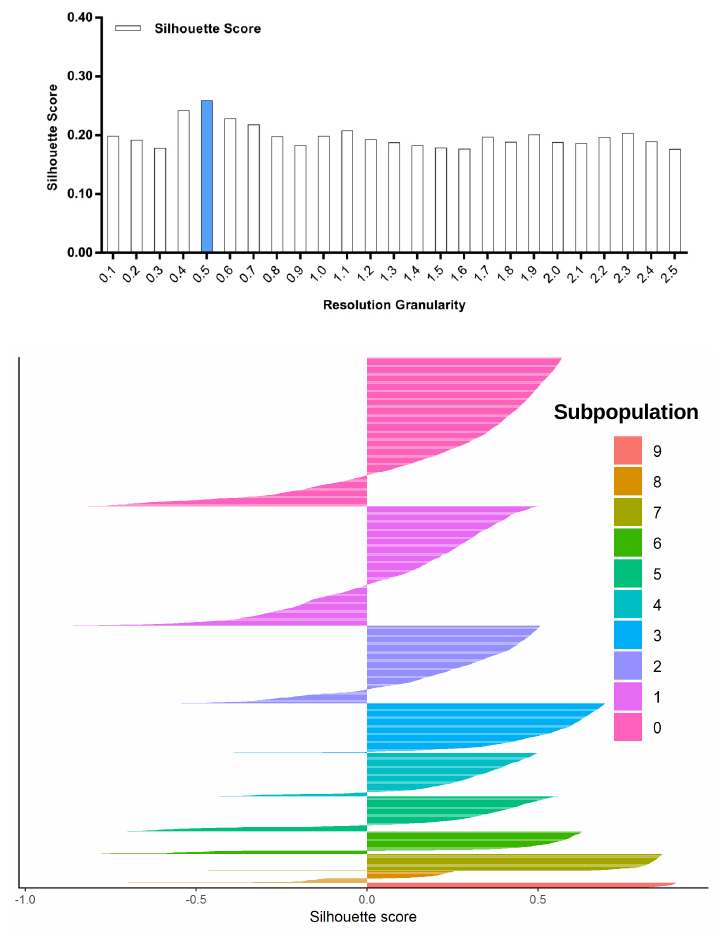

**Figure S14.** Silhouette analysis of the combined cell population treated with DMSO, **SKI-73** and **SKI-73N** guided by the resolution granularity. The resolution granularity of 0.5 for the highest Silhouette score was used for clustering and led to ten subpopulations of the total cell population.

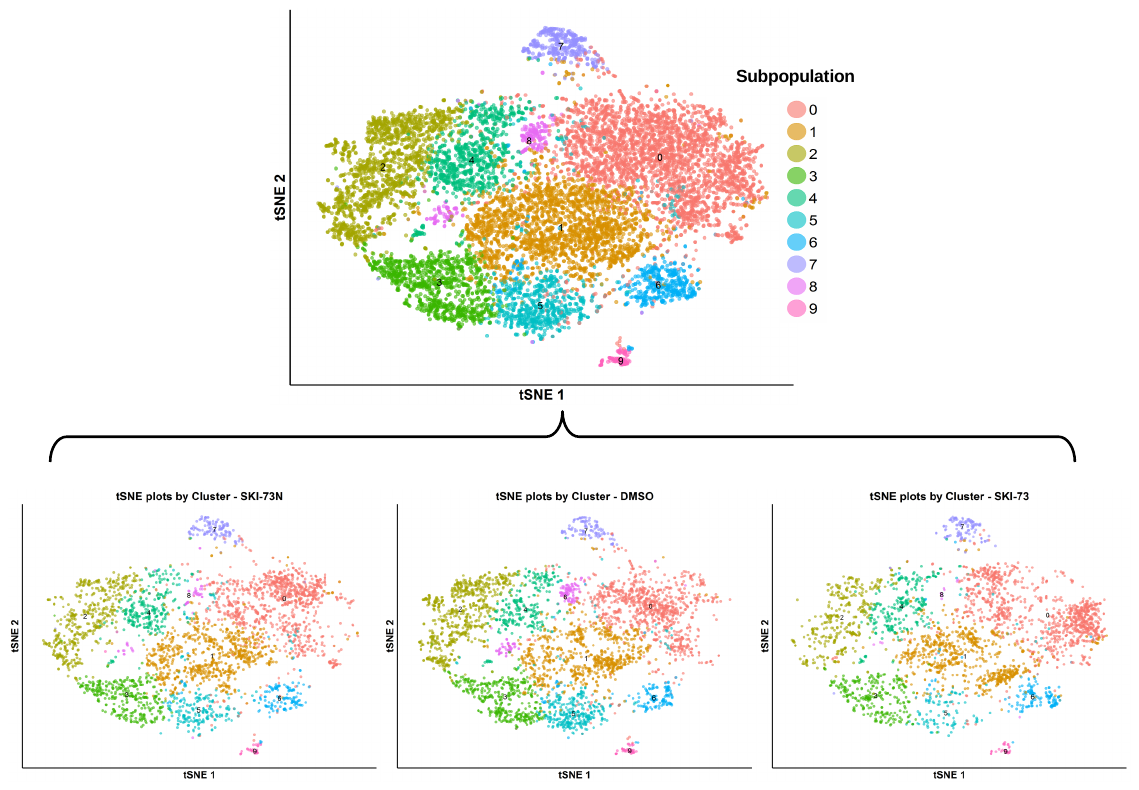

**Figure S15.** tSNE plot of the subpopulations of the combined cells treated with DMSO, **SKI-73** and **SKI-73N**. The combined cell population treated with DMSO, **SKI-73** and **SKI-73N** (the top panel) was clustered into ten subpopulations with the resolution granularity of 0.5. The tSNE plot of the ten clustered subpopulations was split according to their three treatment origins---**SKI-73N**, DMSO and **SKI-73**.

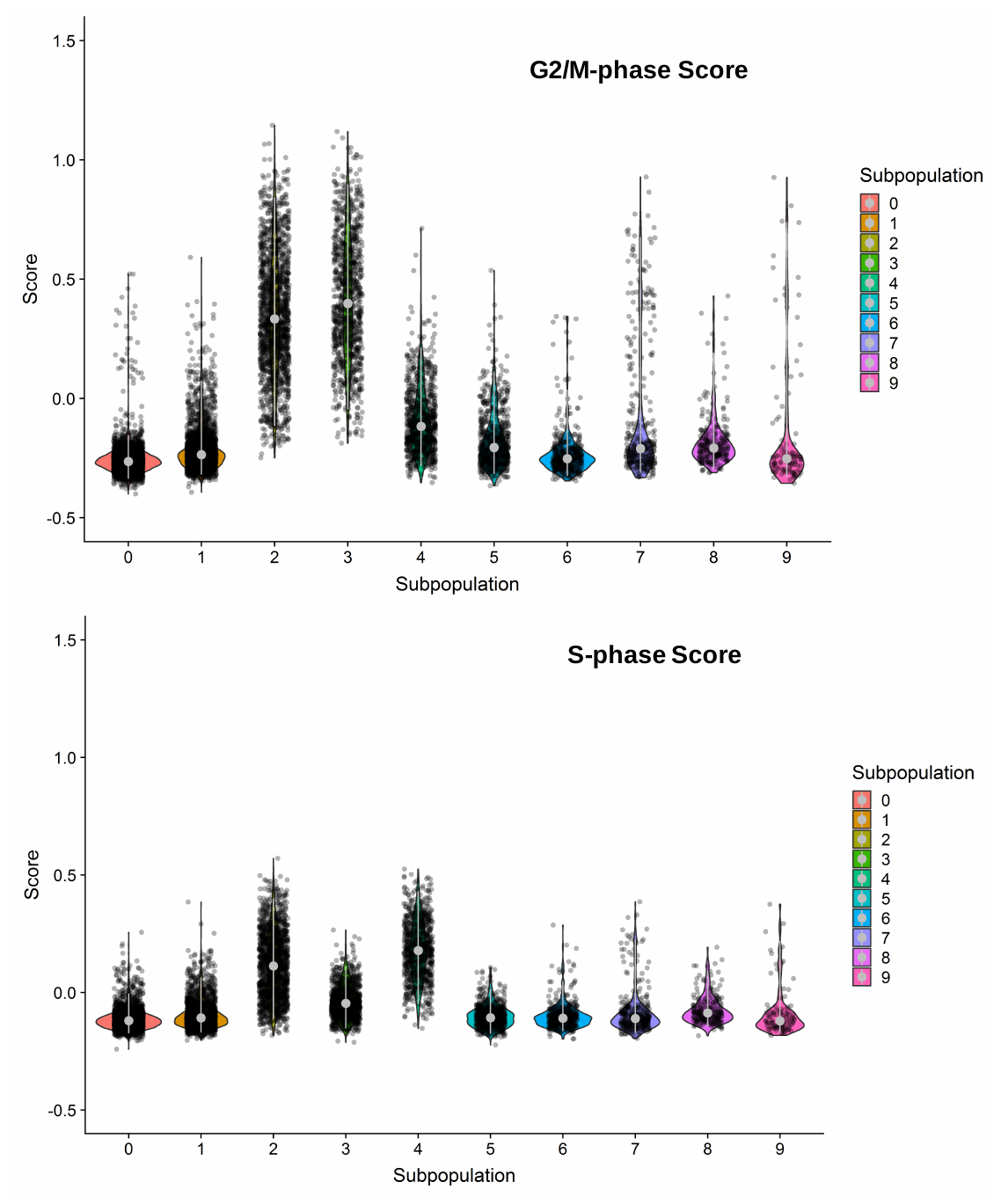

**Figure S16.** G2/M- and S-phase scores of the ten clusters of the combined cell population treated with DMSO, **SKI-73** and **SKI-73N**. Subpopulations 2, 3 and 4 were dominated by G2/M- or S-phase scores.

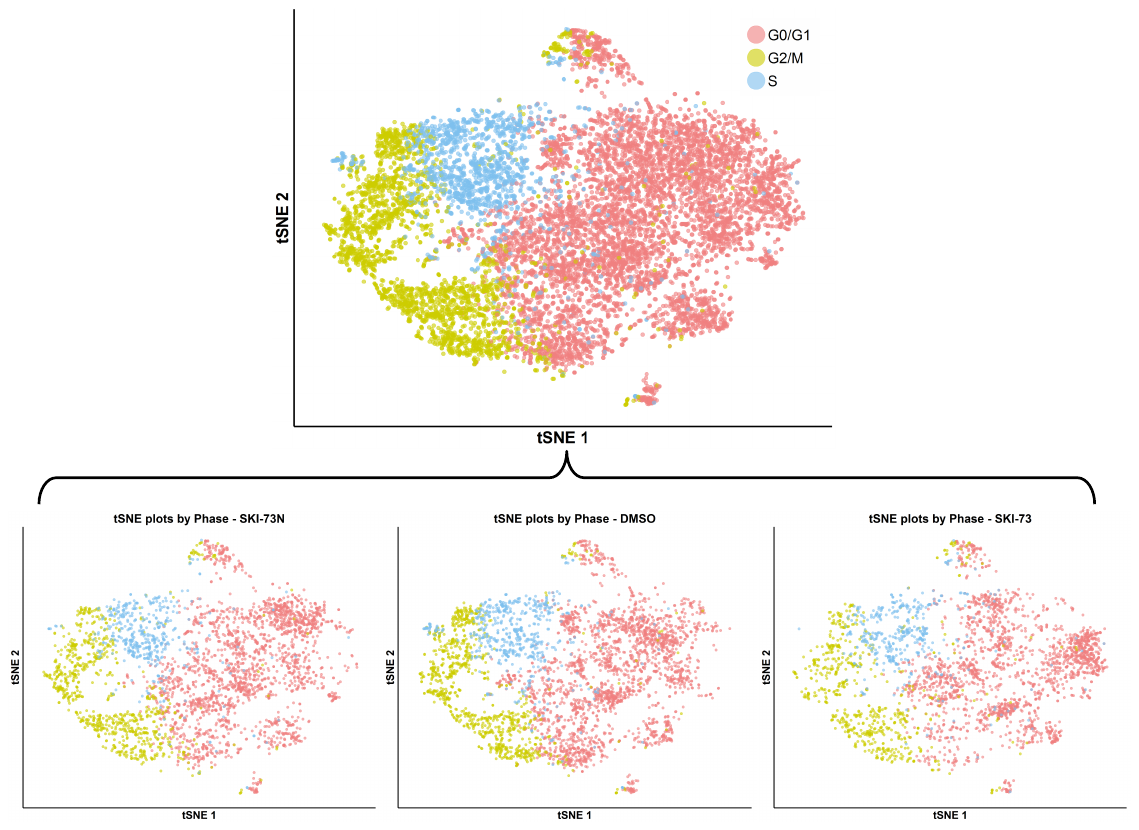

**Figure S17.** tSNE plot with cell-cycle awareness for the combined cell population treated with DMSO, **SKI-73** and **SKI-73N**. The total cell population treated with DMSO, **SKI-73** and **SKI-73N** (the top panel) were classified according to the cell cycle scores of individual cells as described in Supplementary Methods. The tSNE plot of the cell-cycle-aware population was split according to their three treatment origins---**SKI-73N**, DMSO and **SKI-73**.

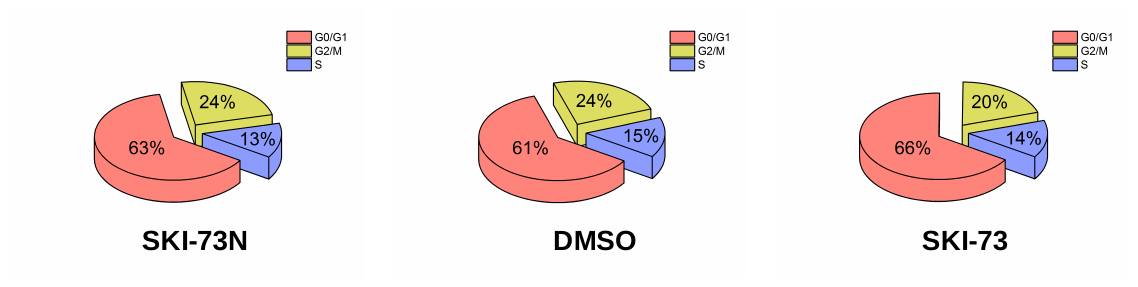

**Figure S18.** Assignment of cell cycle stages of the cells treated with **SKI-73N**, DMSO, and **SKI-73**. Individual cells were classified according to their highest cell-cycle scores for G2/M or S phases as described in Supplementary Methods. Those that cannot be scored for G2/M and S phases were assigned as G0/G1-phase cells. The 2-day treatment with **SKI-73** and **SKI-73N** has a negligible effect on cell cycle.

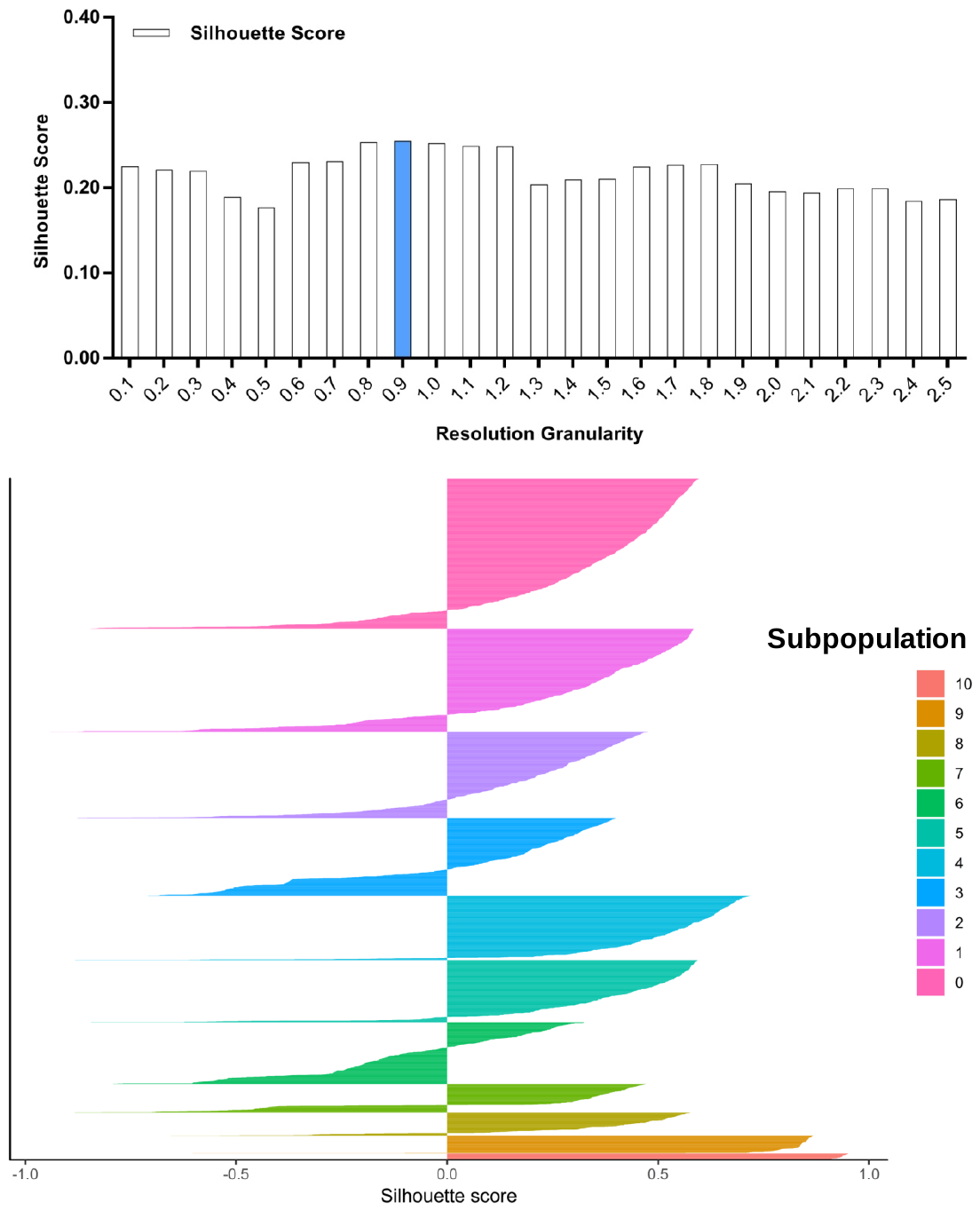

**Figure S19.** Silhouette analysis of the DMSO-treated cells guided by the resolution granularity. The resolution granularity of 0.9 for the highest Silhouette score was used for clustering and led to 6 subpopulations of the DMSO-treated cells.

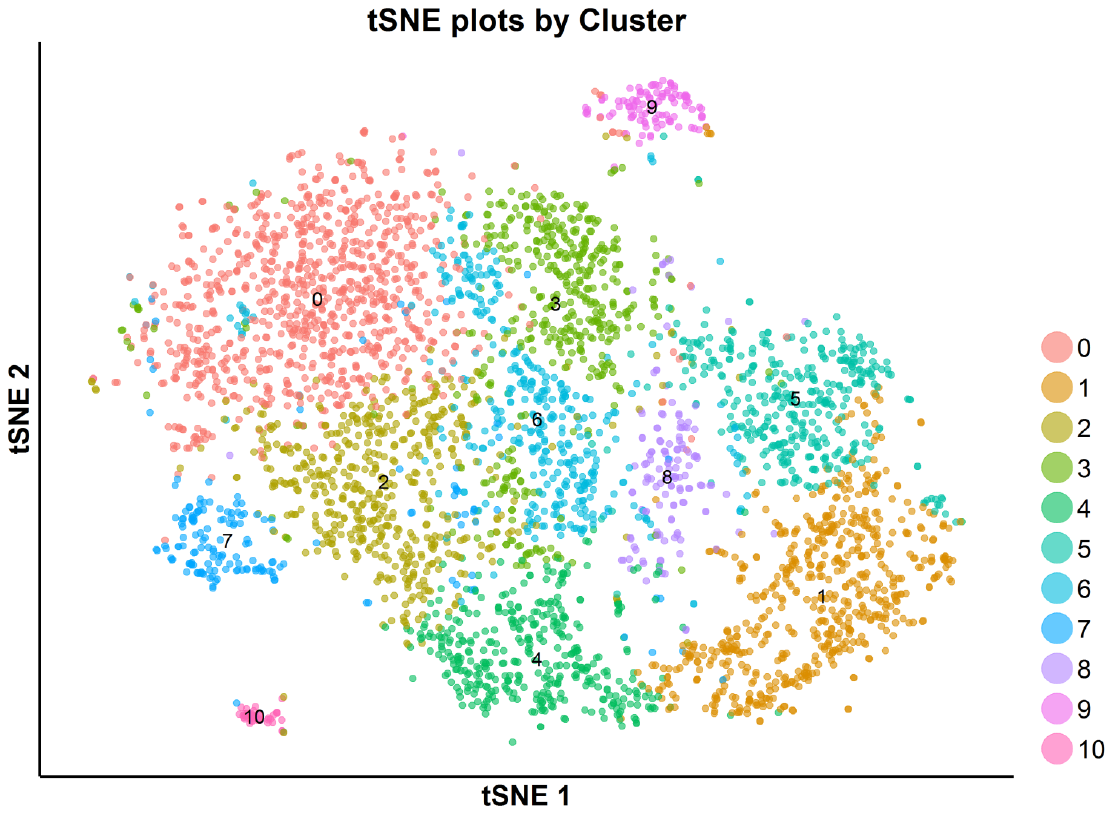

**Figure S20.** tSNE plot of the subpopulations of the DMSO-treated cells. The DMSO-treated cells were clustered into 11 subpopulations with the resolution granularity of 0.9.

**Figure S21.** G2/M- and S-phase scores of the 11 clustered subpopulations of the DMSO-treated cells. Subpopulations 1, 4 and 5 were dominated by G2/M- or S-phase scores.

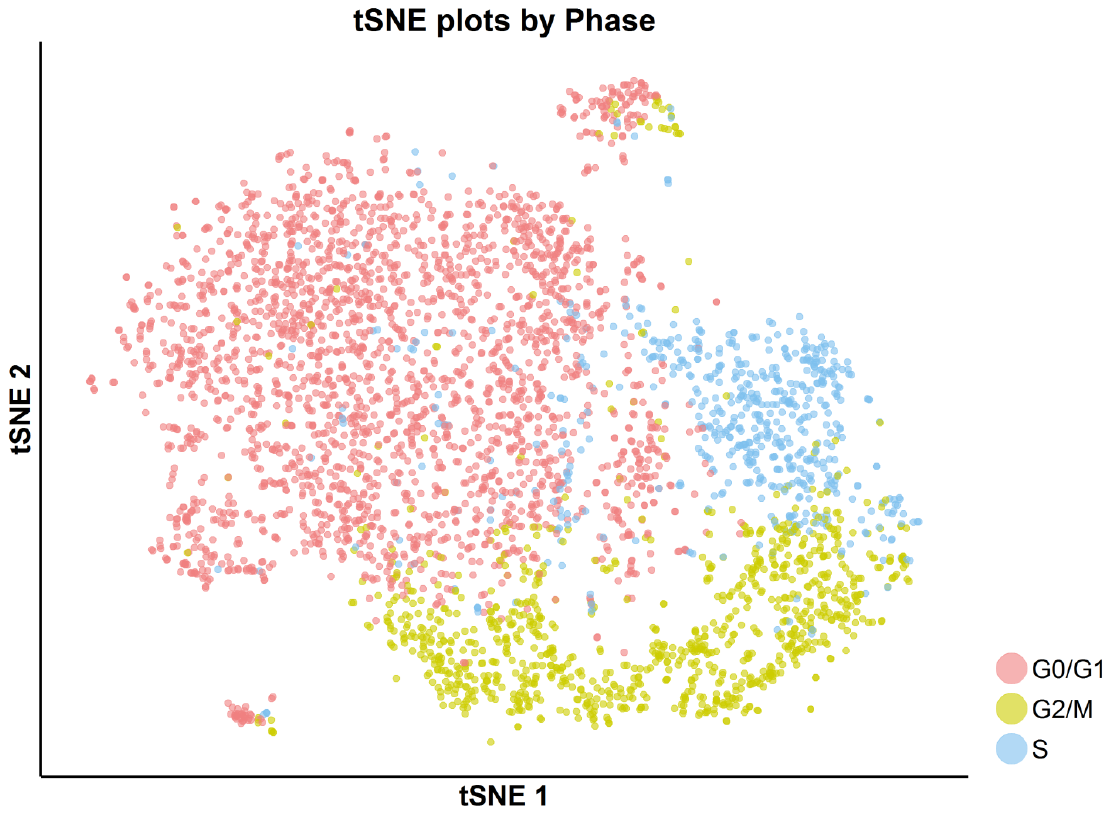

**Figure S22.** tSNE plot with cell-cycle awareness for the DMSO-treated cells. Individual cells were classified according to their cell cycle scores as described in Supplementary Methods.

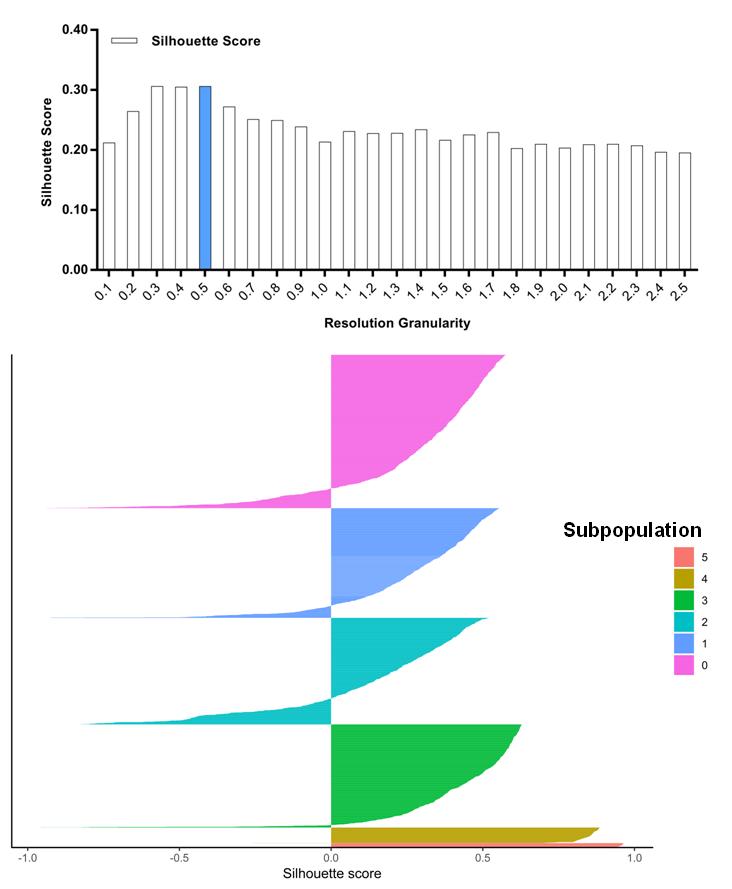

**Figure S23.** Silhouette analysis of the **SKI-73N-**treated cells guided by the resolution granularity. The resolution granularity of 0.5 for the highest Silhouette score was used for clustering and led to 6 subpopulations of the **SKI-73N**-treated cells.

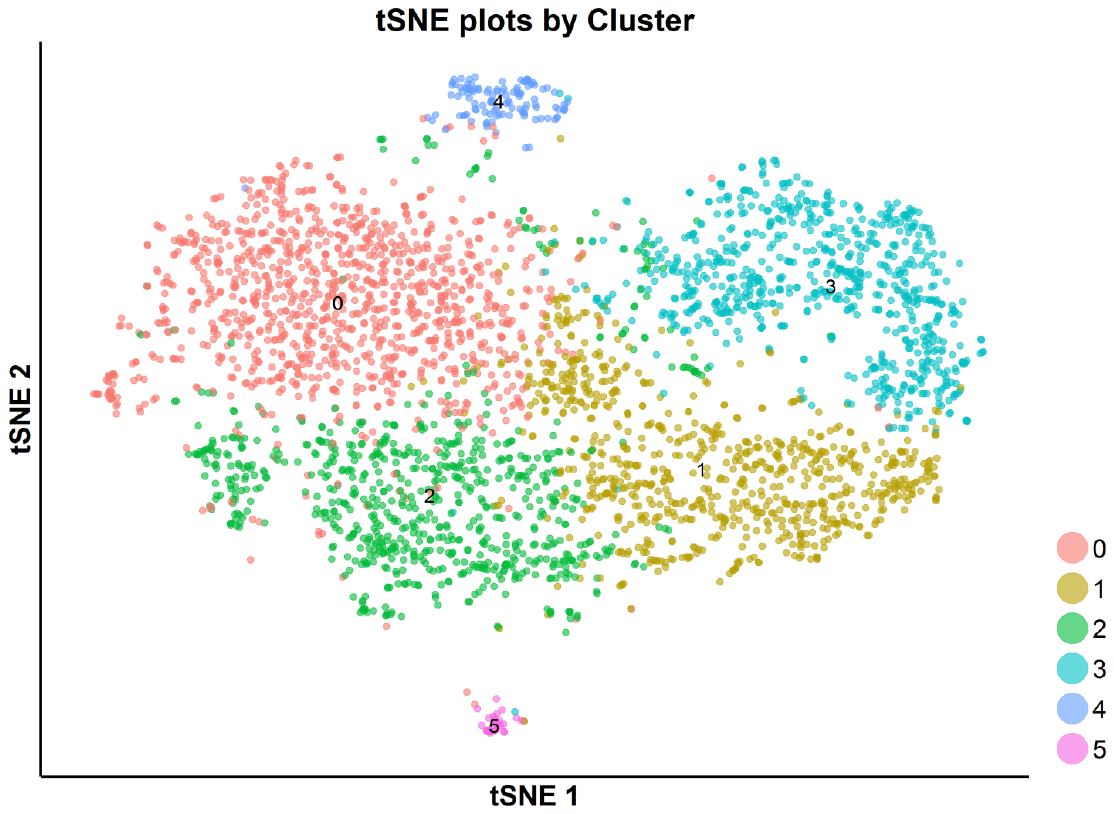

**Figure S24.** tSNE plot of the subpopulations of the **SKI-73N**-treated cells. The **SKI-73N-**treated cells were clustered into 6 subpopulations with the resolution granularity of 0.5.

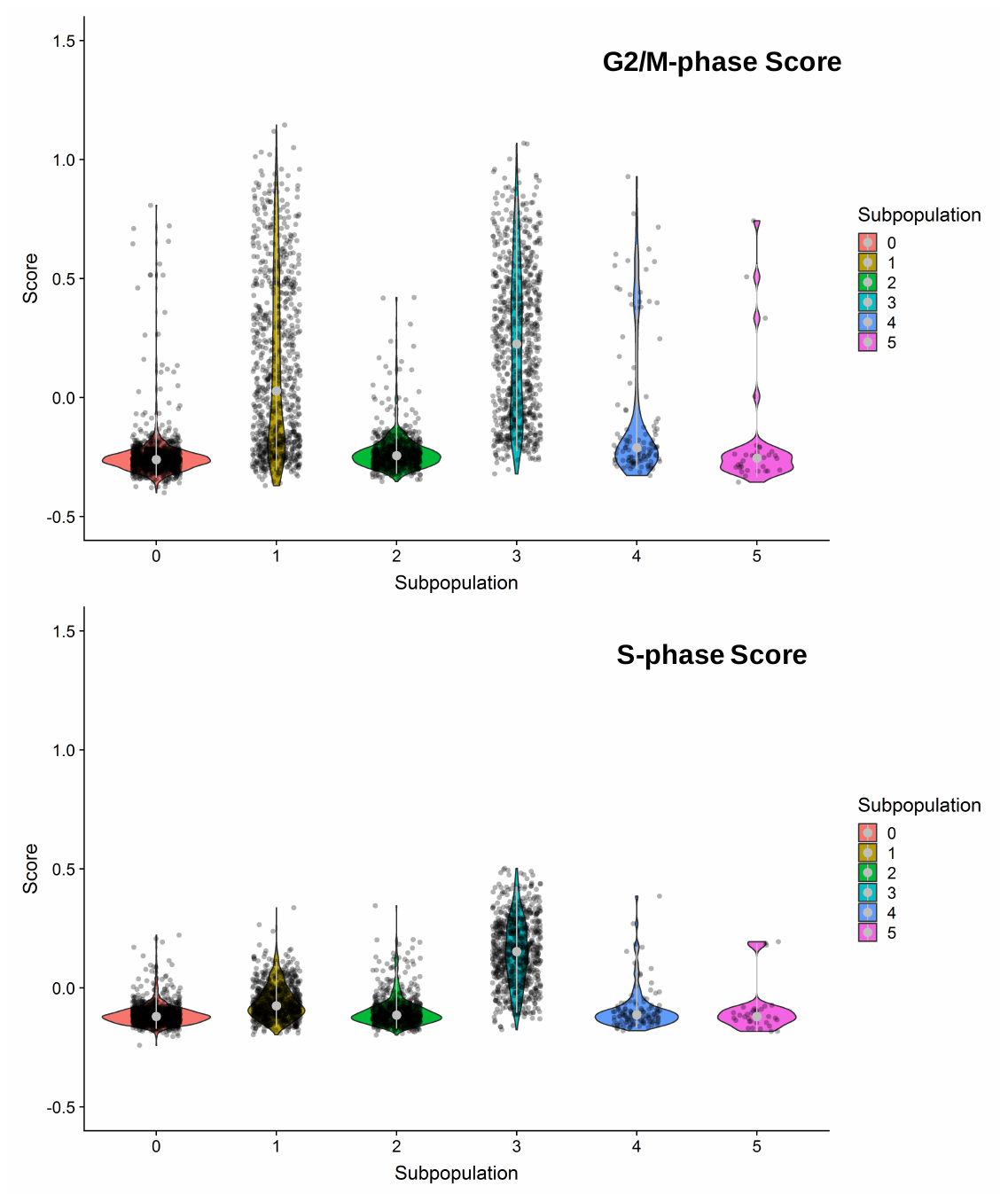

**Figure S25.** G2/M- and S-phase scores of the 6 clustered subpopulation of the **SKI-73N-**treated cells. Subpopulations 1 and 3 were dominated by G2/M- or S-phase scores.

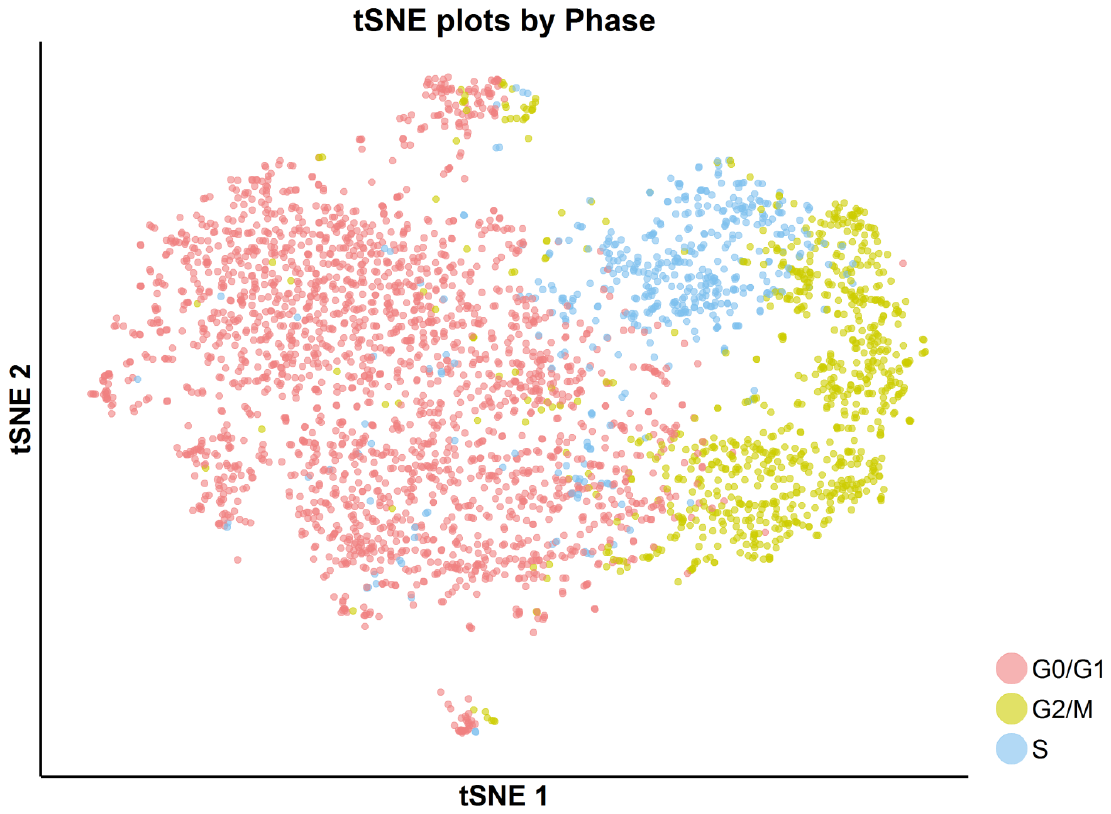

**Figure S26.** tSNE plot with cell-cycle awareness for the **SKI-73N-**treated cells. Individual cells were classified according to their cell cycle scores as described in Supplementary Methods.

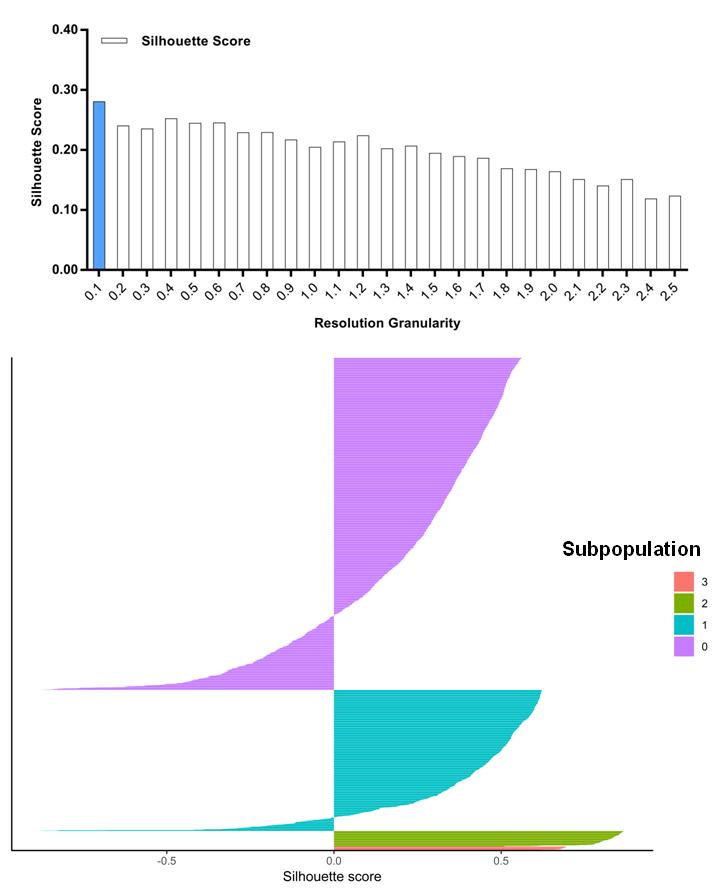

**Figure S27.** Silhouette analysis of the **SKI-73-**treated cells guided by the resolution granularity. The resolution granularity of 0.1 for the highest Silhouette score was used for clustering and led to 4 subpopulations of the **SKI-73-**treated cells.

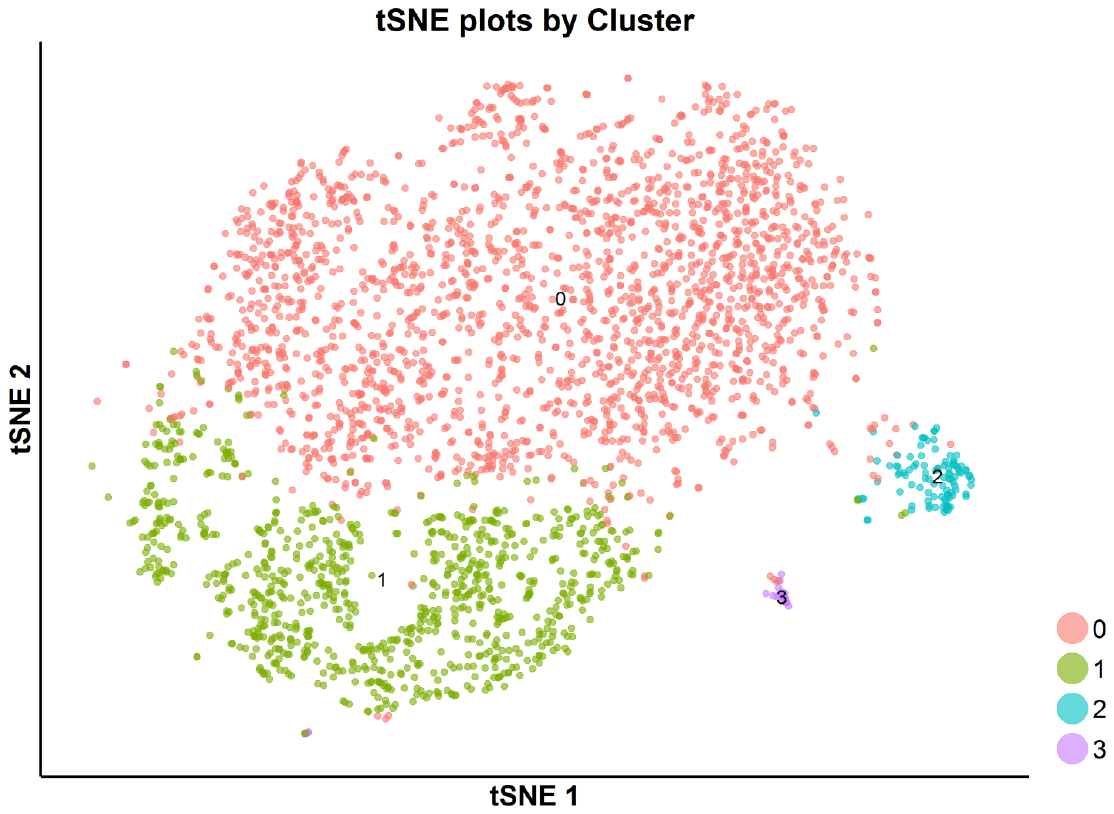

**Figure S28.** tSNE plot of the subpopulations of the **SKI-73-**treated cells. The **SKI-73-**treated cells were clustered into 4 subpopulations with the resolution granularity of 0.1.

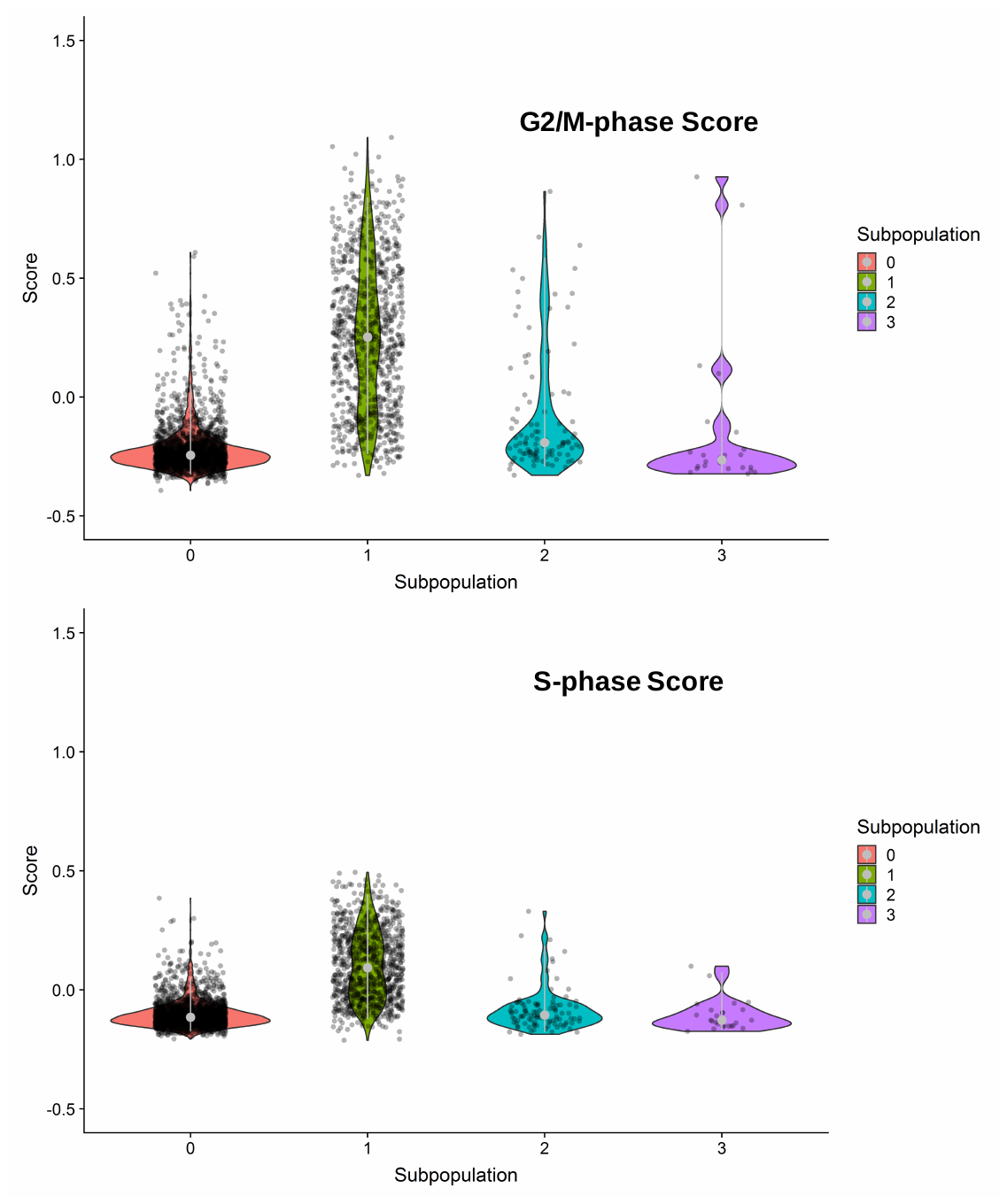

**Figure S29.** G2/M- and S-phase scores of the 4 clustered subpopulation of the **SKI-73-**treated cells. Subpopulation 1 was dominated by G2/M- or S-phase scores.

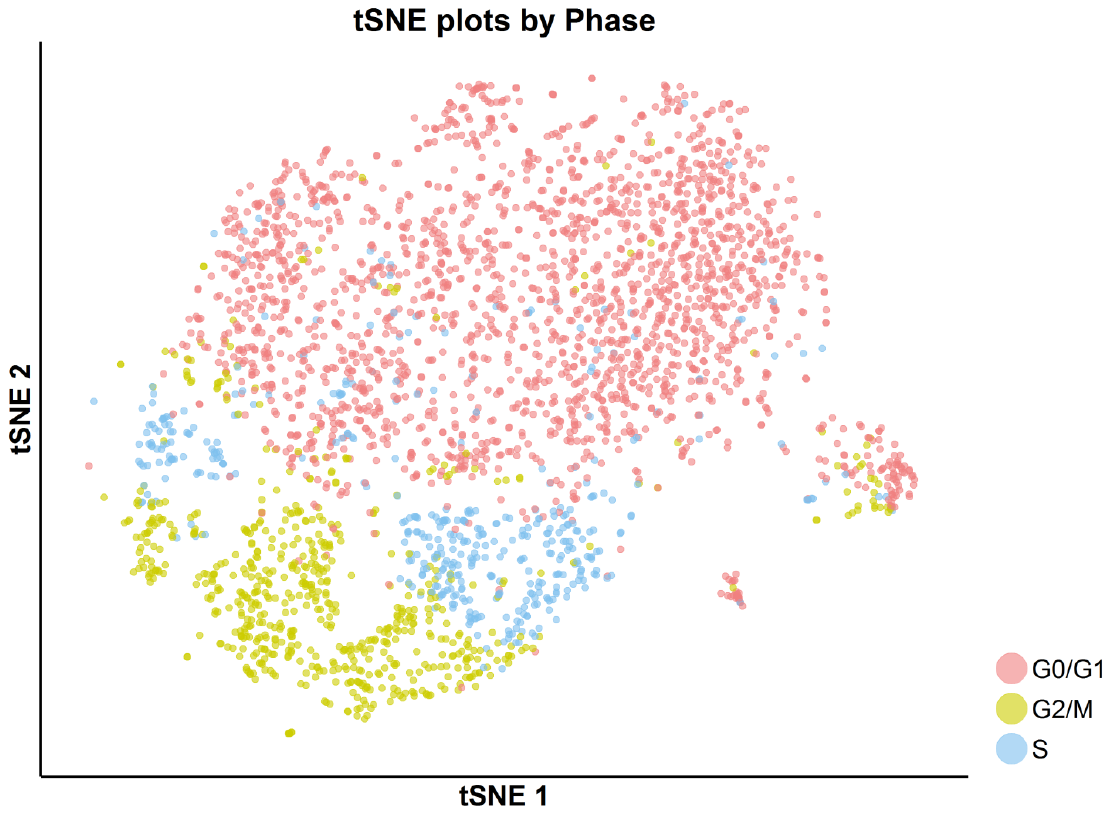

**Figure S30.** tSNE plot with cell-cycle awareness for the **SKI-73-**treated cells. Individual cells were classified according to their cell cycle scores as described in Supplementary Methods.

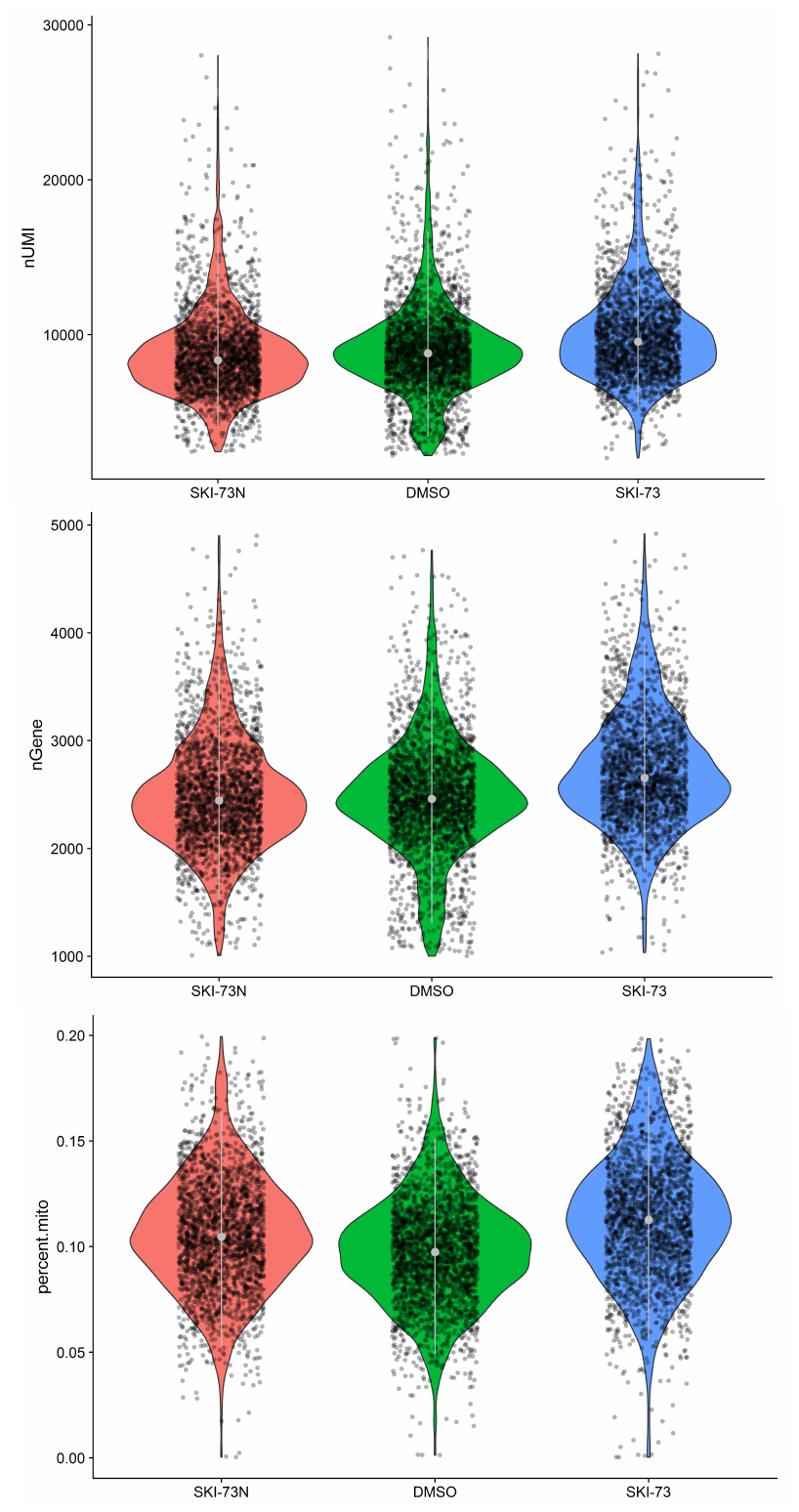

**Figure S31.** Quality control of the G0/G1-phase cells subject for scRNA-seq analysis. A subset of MDA-MB-231 cells after the treatment of **SKI-73N**, DMSO and **SKI-73** was assigned as G0/G1-phase cells as described in Supplementary Methods. The threshold with 1,000~5,000 genes and < 20% mitochondrial RNA transcripts were used to select the G0/G1-phase cells for scRNA-seq analysis with their UMIs (top), the number of genes per cell (middle) and the fraction of mitochondria genes (bottom) shown.

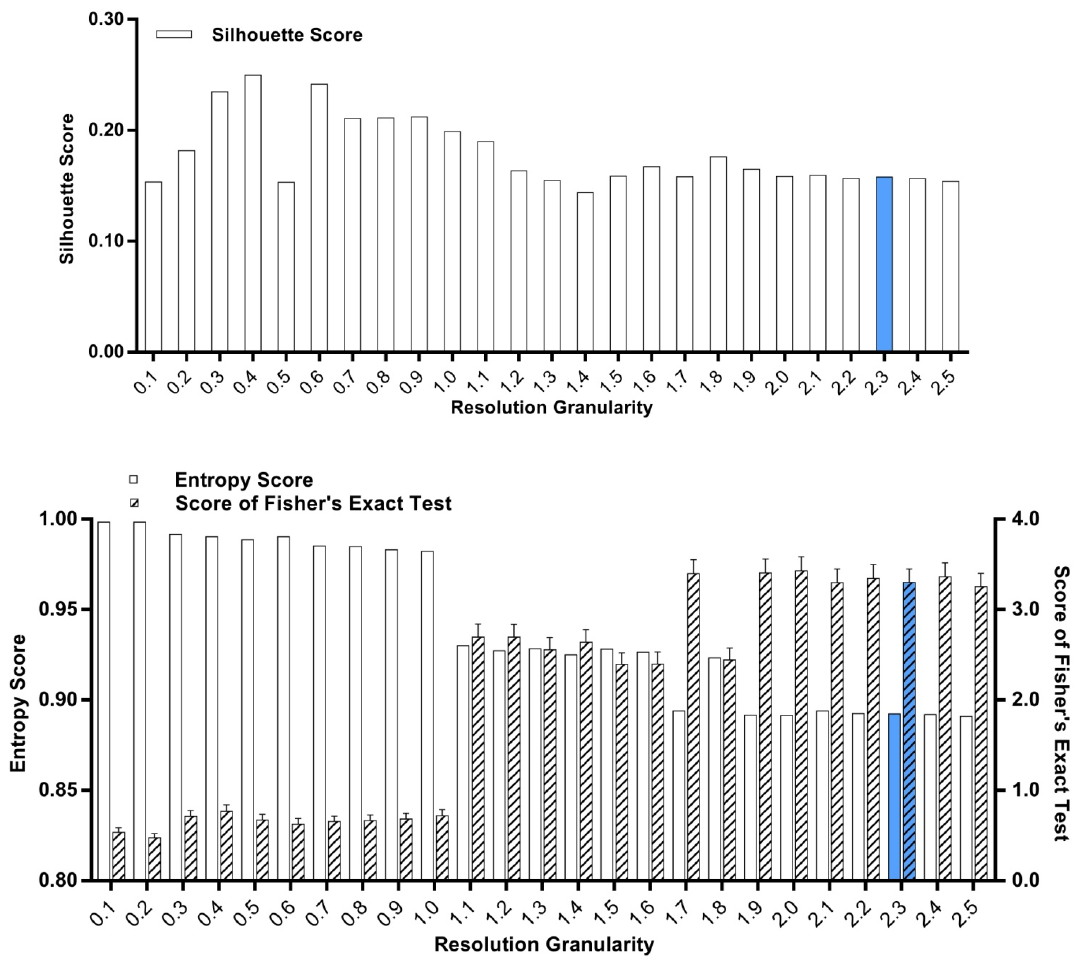

**Figure S32.** Silhouette analysis, entropy analysis and Fisher’s exact test of the classified G0/G1-phase cells after the treatment of **SKI-73N**, DMSO and **SKI-73**. The resolution granularity of 2.3 for a low entropy score and a high Fisher’s exact test score was used for clustering.

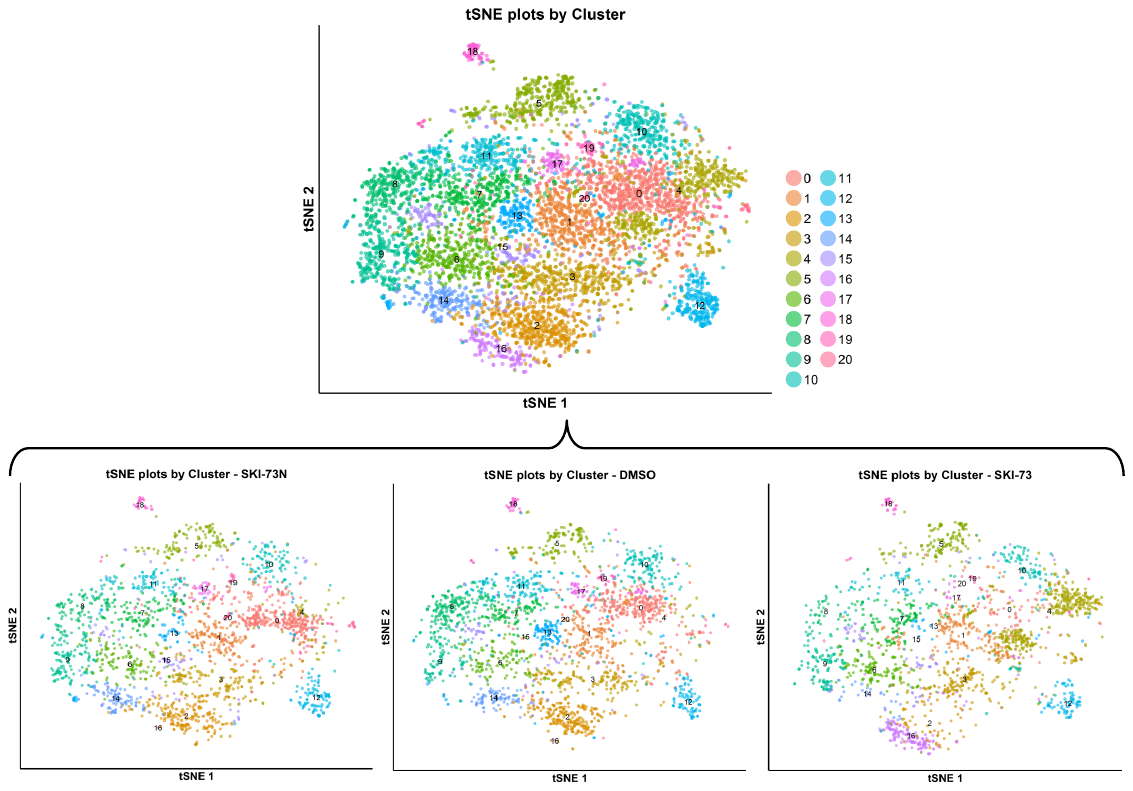

**Figure S33.** tSNE plot of the subpopulations of the classified G0/G1-phase cells after the treatment of **SKI-73N**, DMSO and **SKI-73**. The combined cell population under the three treatment conditions (the top panel) was clustered into 21 subpopulations with the resolution granularity of 2.3. The tSNE plot of the 21 clustered subpopulations was split according to their three treatment origins---**SKI-73N**, DMSO and **SKI-73**.

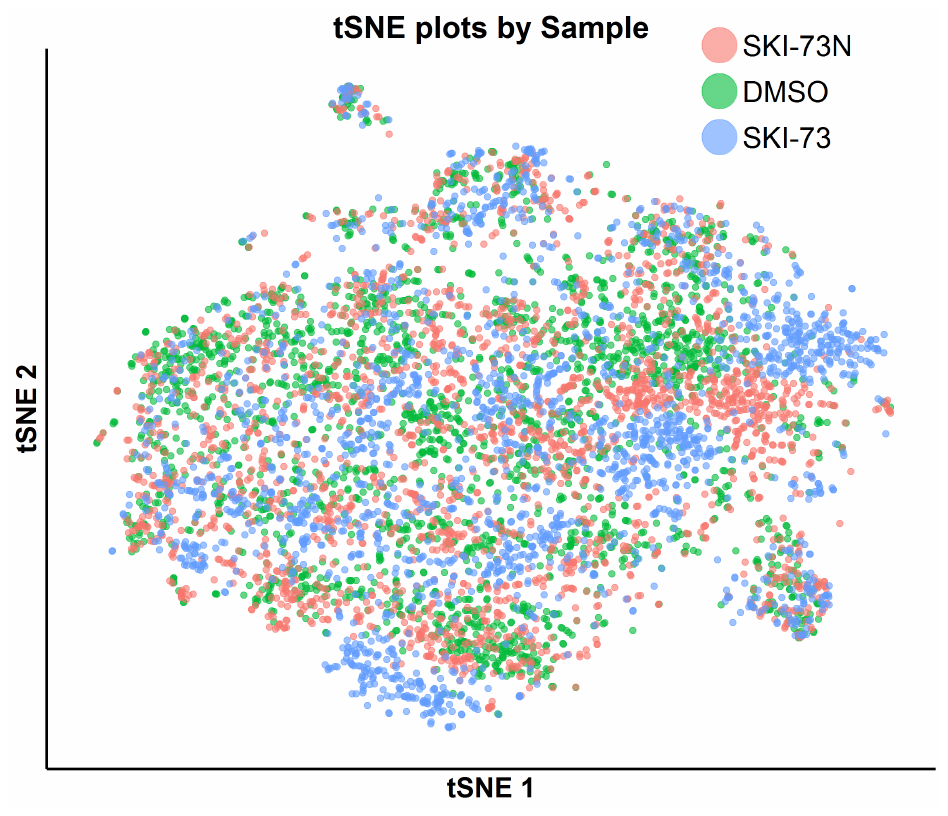

**Figure S34.** tSNE plot of the classified G0/G1-phase cells with awareness of their three treatment origins--- **SKI-73N**, DMSO and **SKI-73**.

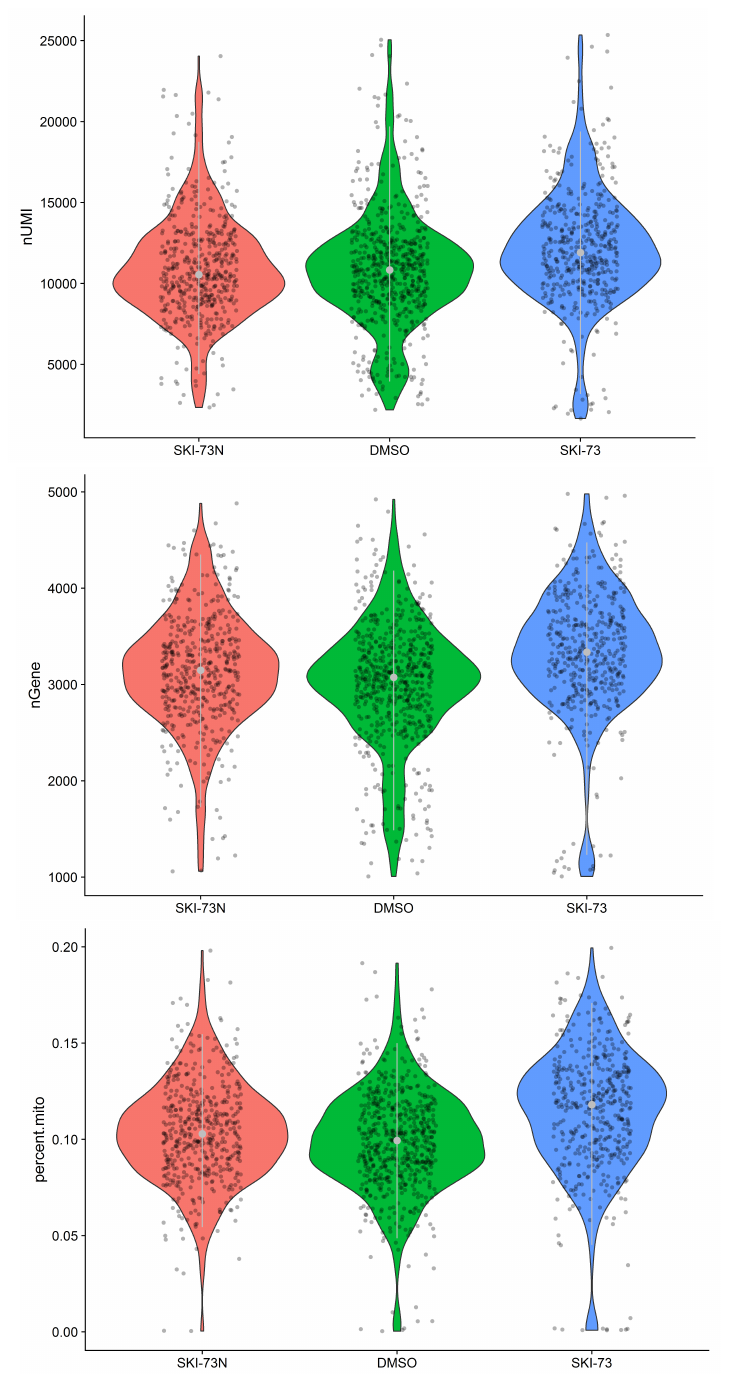

**Figure S35.** Quality control of the S-phase cells subject for scRNA-seq analysis. A subset of MDA-MB-231 cells after the treatment of **SKI-73N**, DMSO and **SKI-73** was assigned as S-phase cells as described in Supplementary Methods. The threshold with 1,000~5,000 genes and < 20% mitochondrial RNA transcripts were used to select the S-phase cells for scRNA-seq analysis with their UMIs (top), the number of genes per cell (middle) and the fraction of mitochondria genes (bottom) shown.

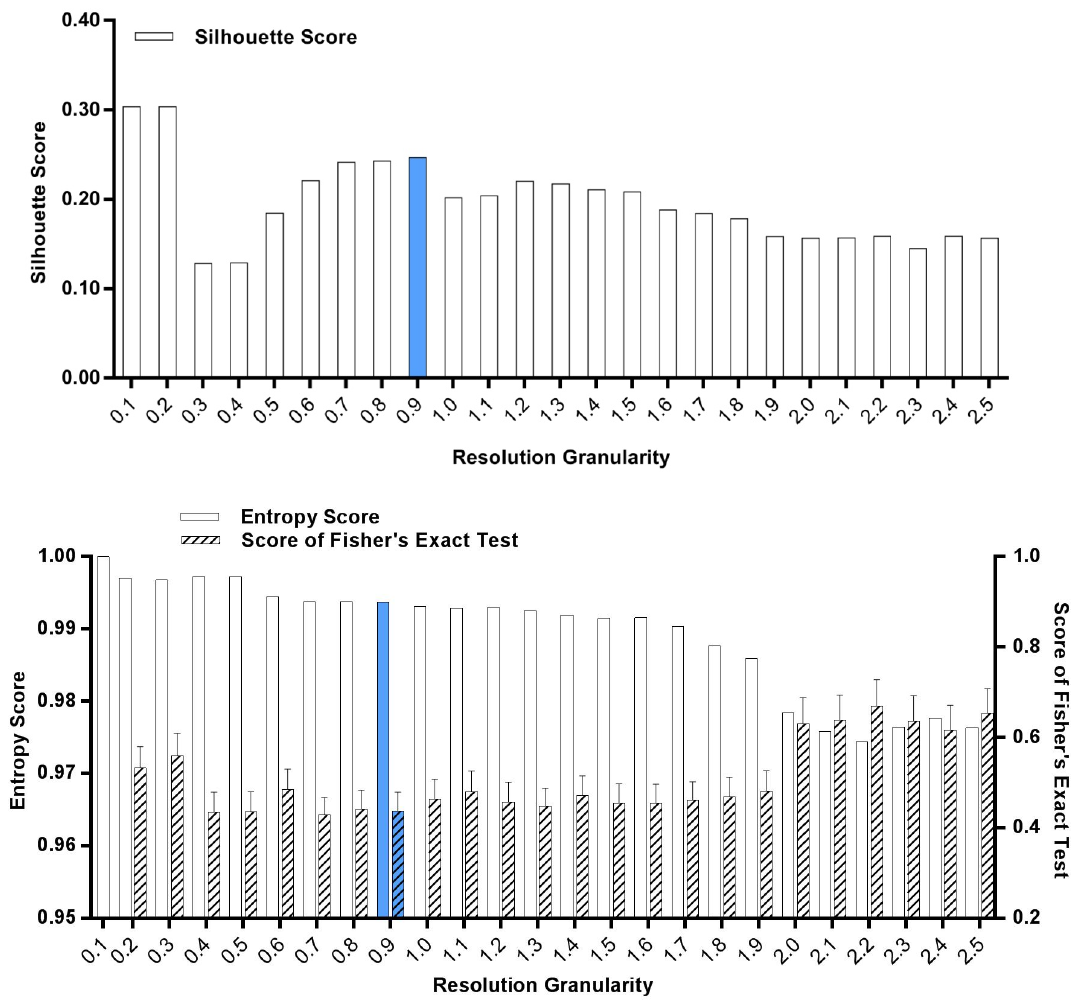

**Figure S36.** Silhouette analysis, entropy analysis and Fisher’s exact test of the classified S-phase cells after the treatment of **SKI-73N**, DMSO and **SKI-73**. The resolution granularity of 0.9 for a low entropy score and a high Fisher’s exact test score was used for clustering.

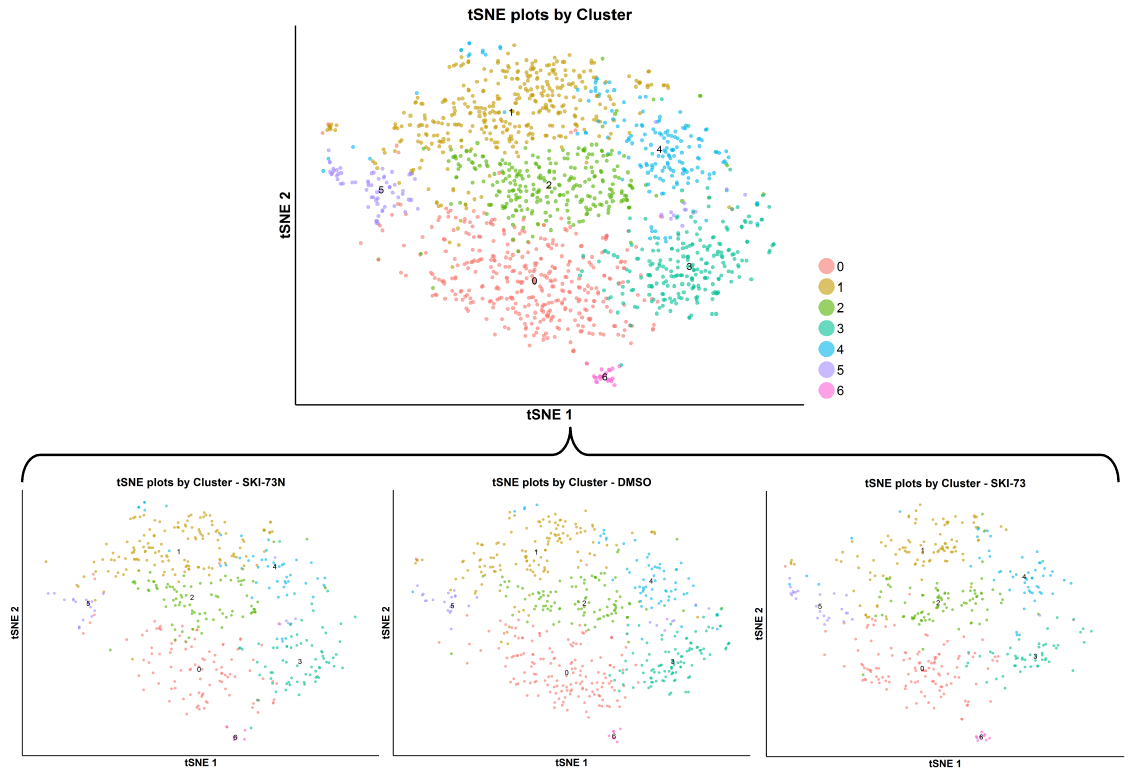

**Figure S37.** tSNE plot of the subpopulations of the classified S-phase cells after the treatment of **SKI-73N**, DMSO and **SKI-73**. The combined cell population under the three treatment conditions (the top panel) was clustered into7 subpopulations with the resolution granularity of 0.9. The tSNE plot of the 7 clustered subpopulations was split according to their three treatment origins---**SKI-73N**, DMSO and **SKI-73**.

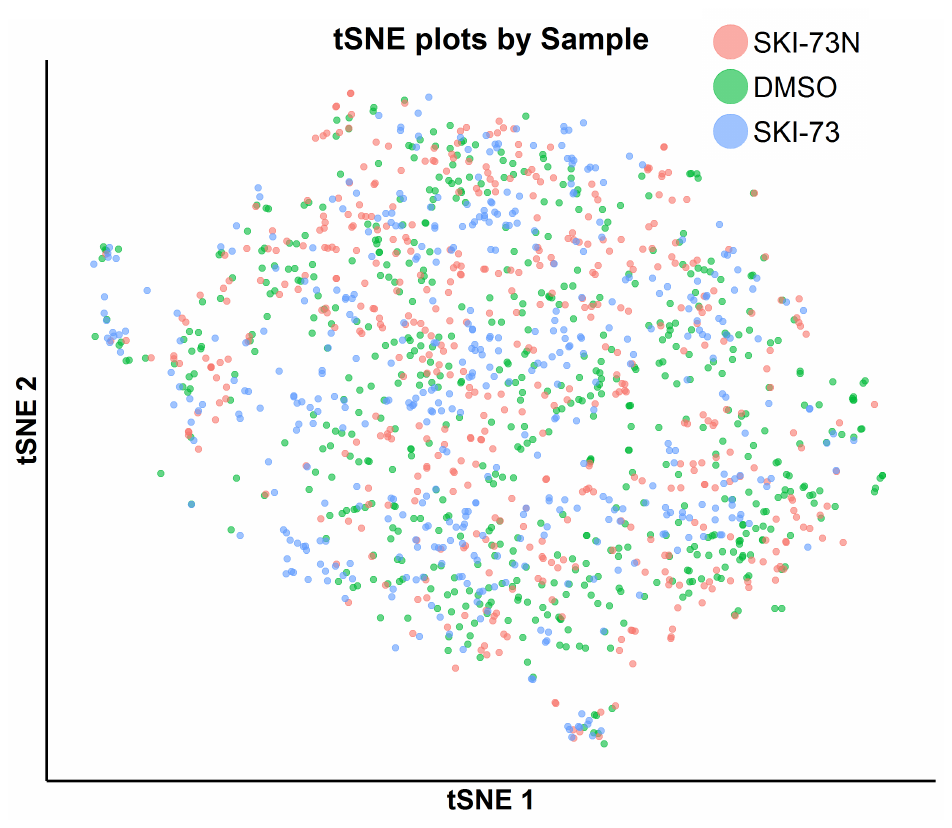

**Figure S38.** tSNE plot of the classified S-phase cells with awareness of their three treatment origins--- **SKI-73N**, DMSO and **SKI-73**.

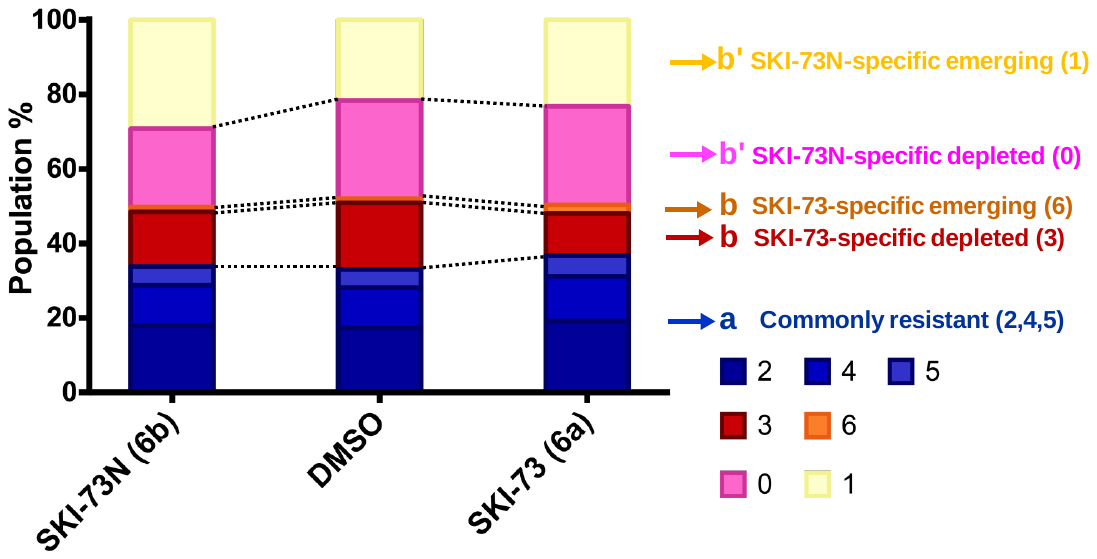

**Figure S39.** Population analysis of the S-phase cells after the treatment of **SKI-73** and **SKI-73N**. Depleted or emerging subpopulations were defined as the < 0.8 and > 1.2 ratios, respectively, for *d_(SKI-73N,i),total_*/*d_(DMSO,i),total_* or *d_(SKI-73,i),total_*/*d_(DMSO,i),total_* as described in Supplementary Methods. Resistant subpopulations were defined within the 0.8~1.2 ratio range for both *d_(SKI-73N,i),total_*/*d_(DMSO,i),total_* and *d_(SKI-73,i),total_*/*d_(DMSO,i),total_*. Here commonly resistant (2, 4 and 5), **SKI-73**-specific depleted/emerging (3 and 6), and **SKI-73N**-specific depleted/emerging (0 and 1) subpopulations are color-coded in blue, red/orange, and pink/yellow, respectively.

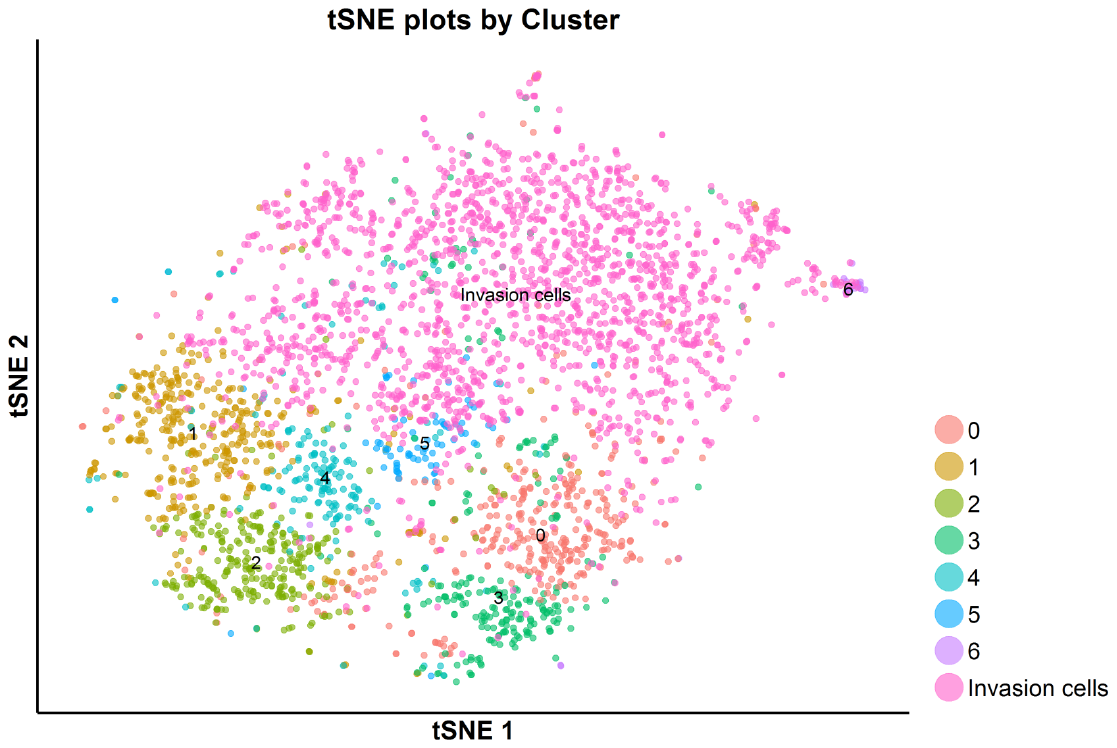

**Figure S40.** tSNE plot of the total S-phase cells analyzed by scRNA-seq---**SKI-73N-**, DMSO**-** and **SKI-73**-treated cells and invasion cells. The S-phase cells upon the treatment with **SKI-73N**, DMSO and **SKI-73** were clustered into 7 subpopulations as described in Figure 37.

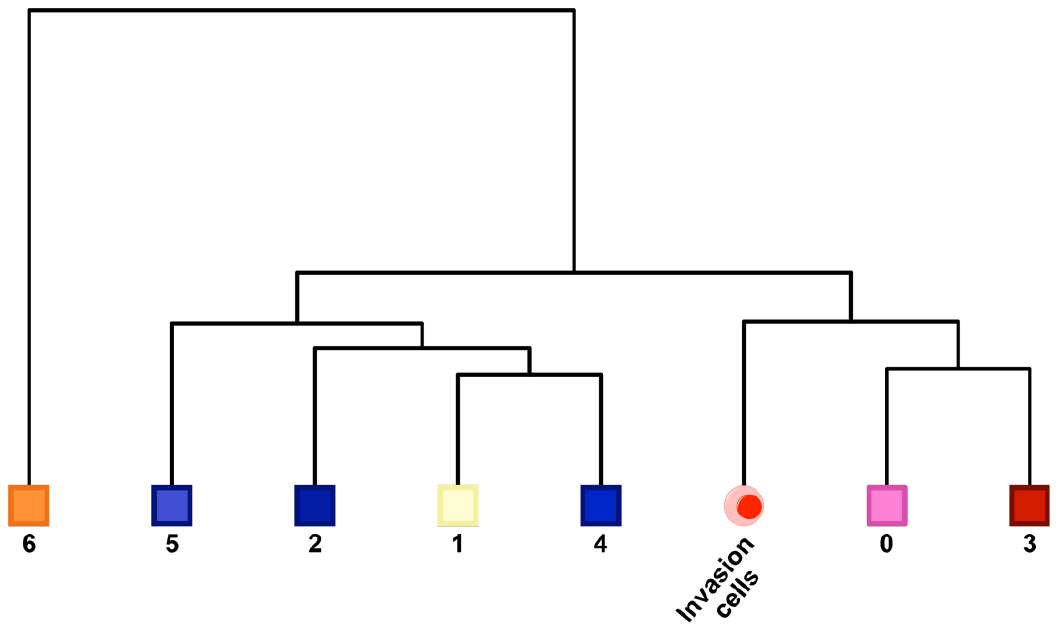

**Figure S41.** Unsupervised correlation analysis of the S-phase subpopulations. Within the **SKI-73N**-, DMSO- and **SKI-73**-treated S-phase cells (7 subpopulations) and the S-phase invasion cells, the correlation analysis of the 8 subsets was conducted with the algorithm in the Seurat package as described in Supplementary Methods. The subpopulations 0 and 3 are closely related to invasion cells.

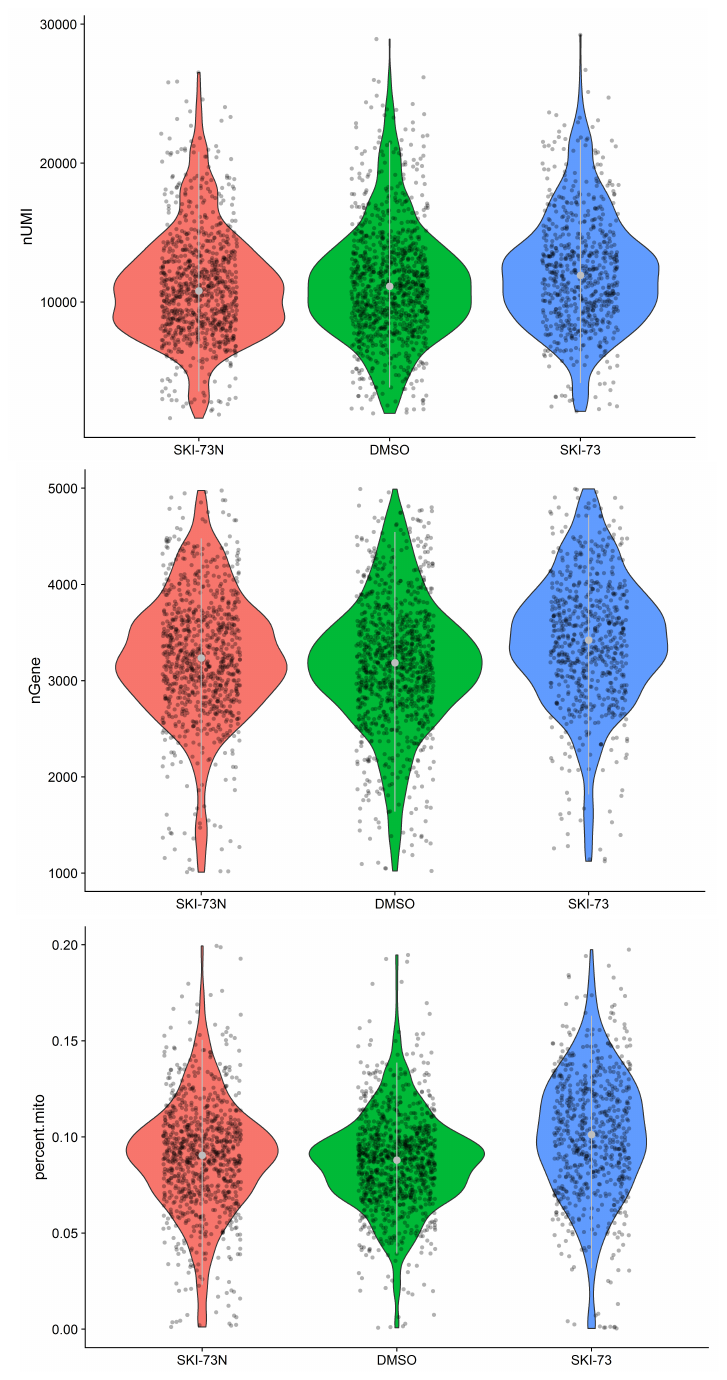

**Figure S42.** Quality control of the G2/M-phase cells subject for scRNA-seq analysis. A subset of MDA-MB-231 cells after the treatment of **SKI-73N**, DMSO and **SKI-73** was assigned into G2/M-phase cells as described in Supplementary Methods. The threshold with 1,000~5,000 genes and < 20% mitochondrial RNA transcripts were used to select the G2/M-phase cells for scRNA-seq analysis with their UMIs (top), the number of genes per cell (middle) and the fraction of mitochondria genes (bottom) shown.

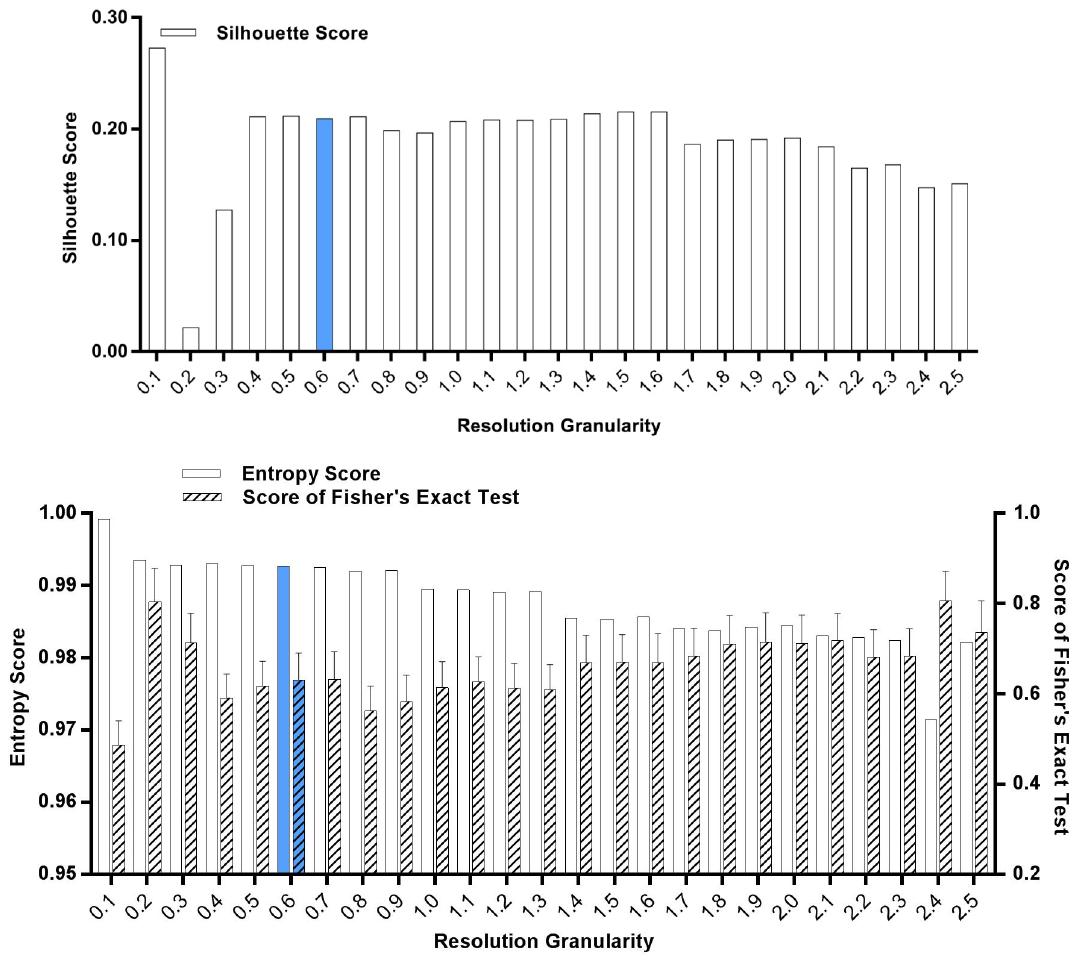
**Figure S43.** Silhouette analysis, entropy analysis and Fisher’s exact test of the classified G2/M-phase cells after the treatment of **SKI-73N**, DMSO and **SKI-73**. The resolution granularity of 0.6 for a low entropy score and a high Fisher’s exact test score was used for clustering.

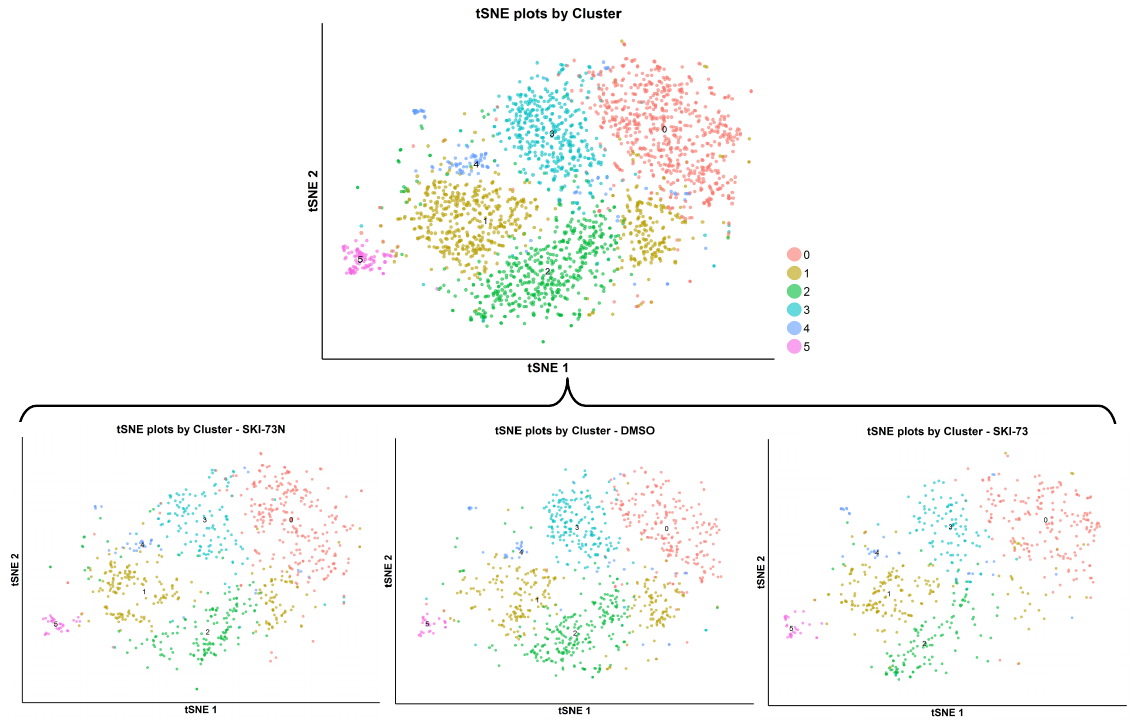

**Figure S44.** tSNE plot of the subpopulations of the classified G2/M-phase cells after the treatment of **SKI-73N**, DMSO and **SKI-73**. The combined cell population under the three treatment conditions (the top panel) was clustered into 6 subpopulations with the resolution granularity of 0.6. The tSNE plot of the 6 clustered subpopulations was split according to their three treatment origins---**SKI-73N**, DMSO and **SKI-73**.

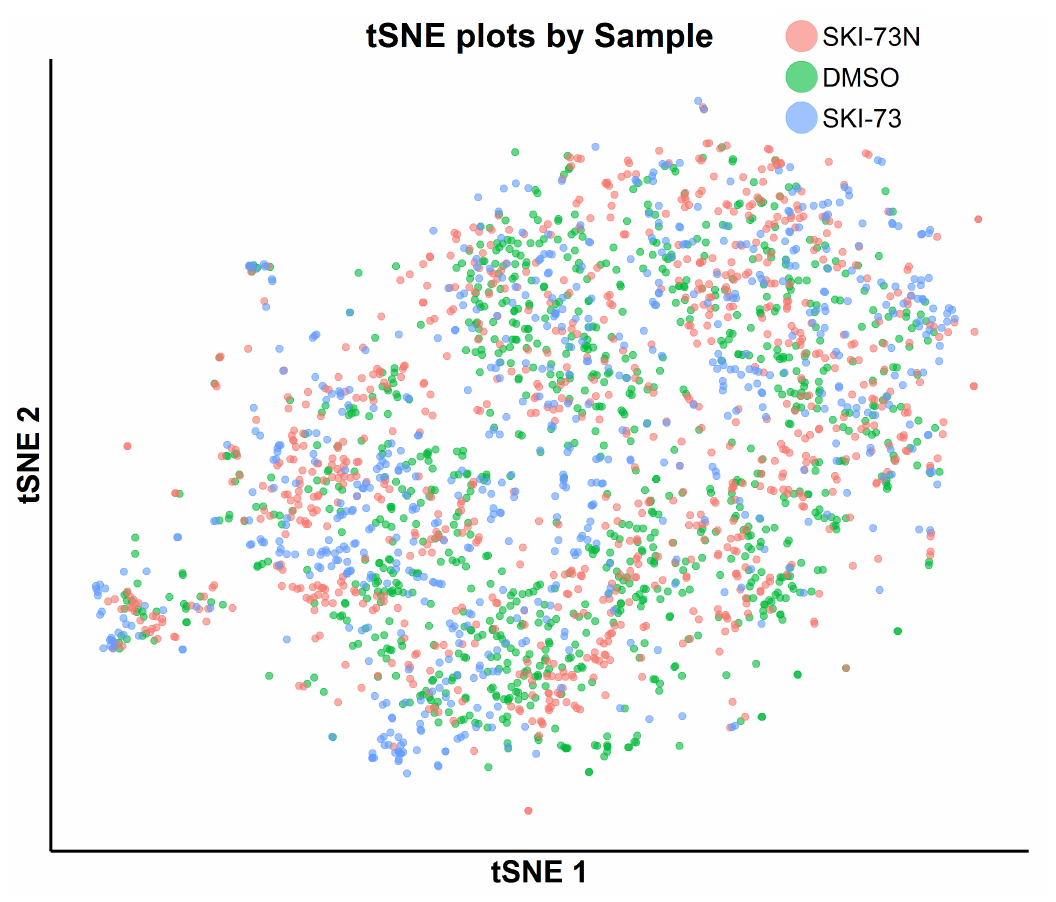

**Figure S45.** tSNE plot of the classified G2/M-phase cells with awareness of their three treatment origins--- **SKI-73N**, DMSO and **SKI-73**.

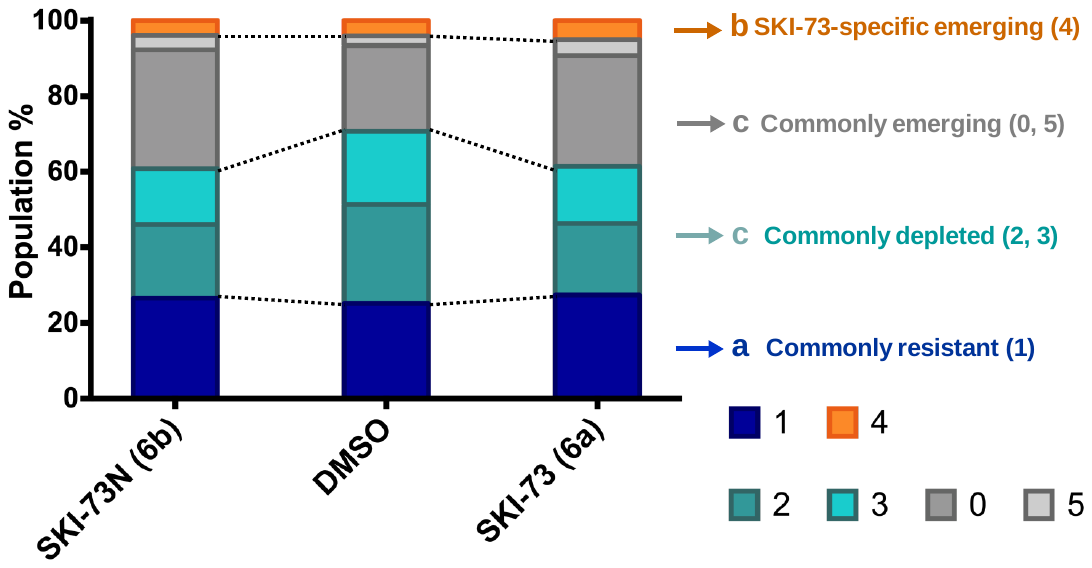

**Figure S46.** Population analysis of the G2/M-phase cells after the treatment of **SKI-73** and **SKI-73N**. Depleted or emerging subpopulations were defined as the < 0.8 and > 1.2 ratios, respectively, for *d_(SKI-73N,i),total_*/*d_(DMSO,i),total_* or *d_(SKI-73,i),total_*/*d_(DMSO,i),total_* as described in Supplementary Methods. Resistant subpopulations were defined within the 0.8~1.2 ratio range for both *d_(SKI-73N,i),total_*/*d_(DMSO,i),total_* and *d_(SKI-73,i),total_*/*d_(DMSO,i),total_*. Here commonly resistant/depleted/emerging (1/2, 3/0, 5) and **SKI-73**-specific emerging (4) subpopulations are color-coded in blue/cyan/grey and orange, respectively.

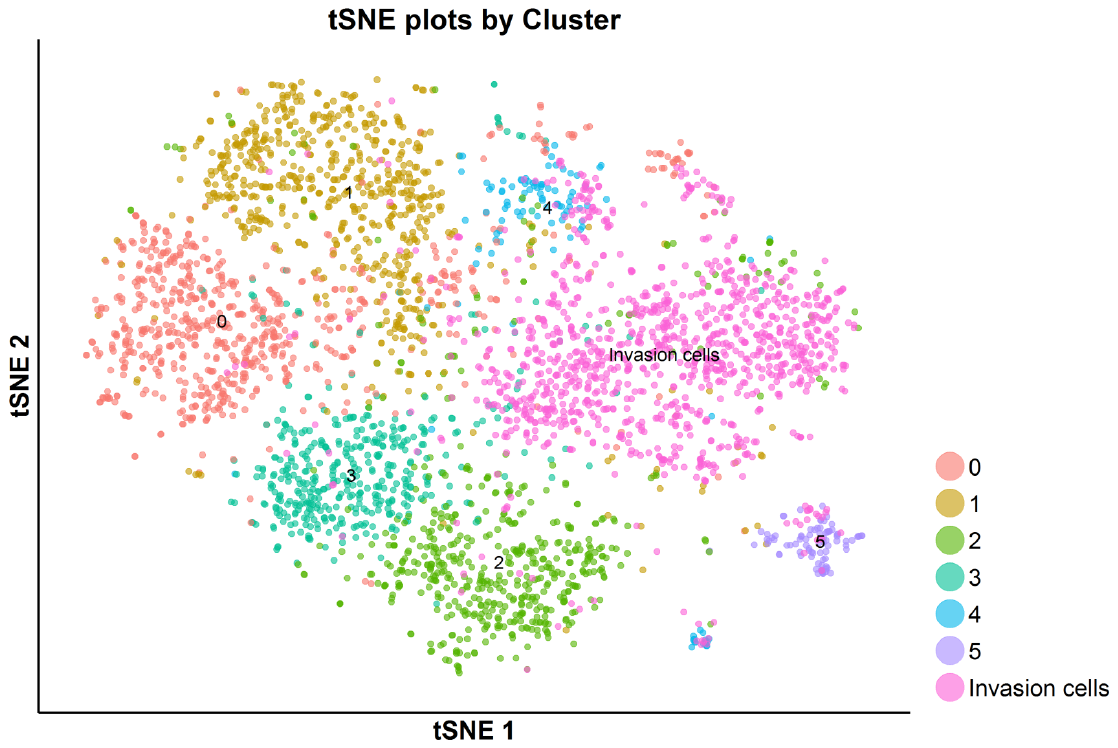

**Figure S47.** tSNE plot of the total G2/M-phase cells analyzed by scRNA-seq---**SKI-73N-**, DMSO**-** and **SKI-73**-treated cells and invasion cells. The G2/M-phase cells upon the treatment with **SKI-73N**, DMSO and **SKI-73** were clustered into 6 subpopulations as described in Figure 44.

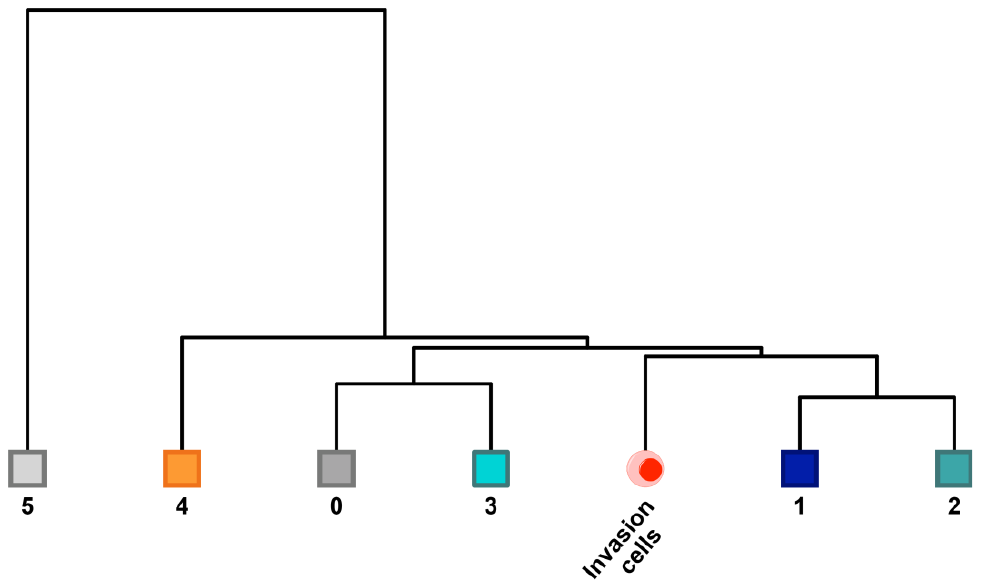

**Figure S48.** Unsupervised correlation analysis of the G2/M-phase subpopulations. Within the **SKI-73N**-, DMSO- and **SKI-73**-treated G2/M-phase cells (6 subpopulations) and the G2/M-phase invasion cells, the correlation analysis of the 7 subsets was conducted with the algorithm in the Seurat package as described in Supplementary Methods. The subpopulations 1 and 2 are closely related to invasion cells.

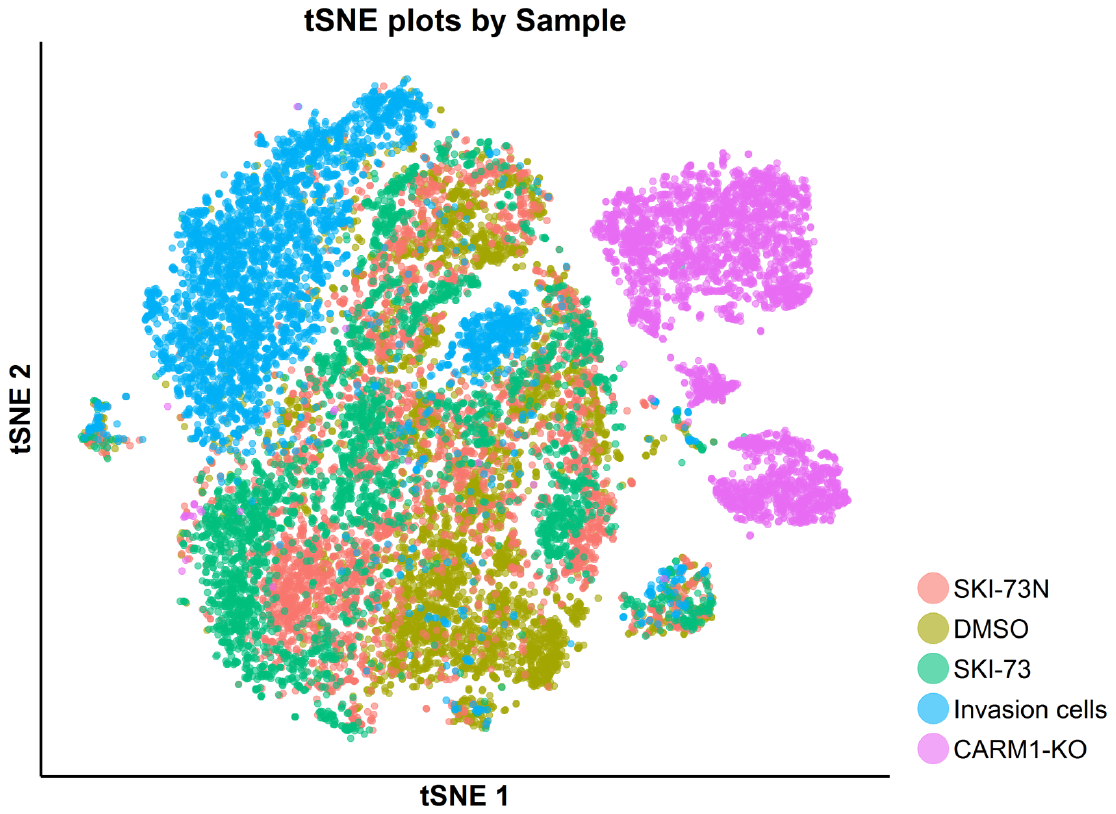

**Figure S49.** tSNE plot of the total cell population with five origins---**SKI-73N-**, DMSO**-** and **SKI-73**-treated cells, invasion cells and CARM1-*KO* cells.

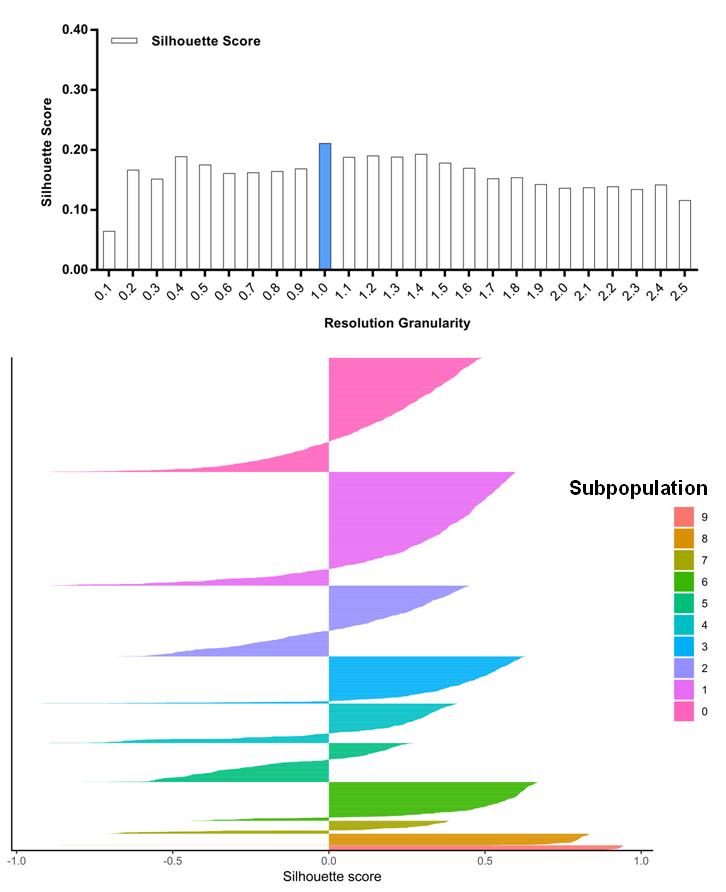

**Figure S50.** Silhouette analysis of invasion cells guided by the resolution granularity. The resolution granularity of 1.0 for the highest Silhouette score was used for clustering and led to 10 subpopulations of the invasion cells.

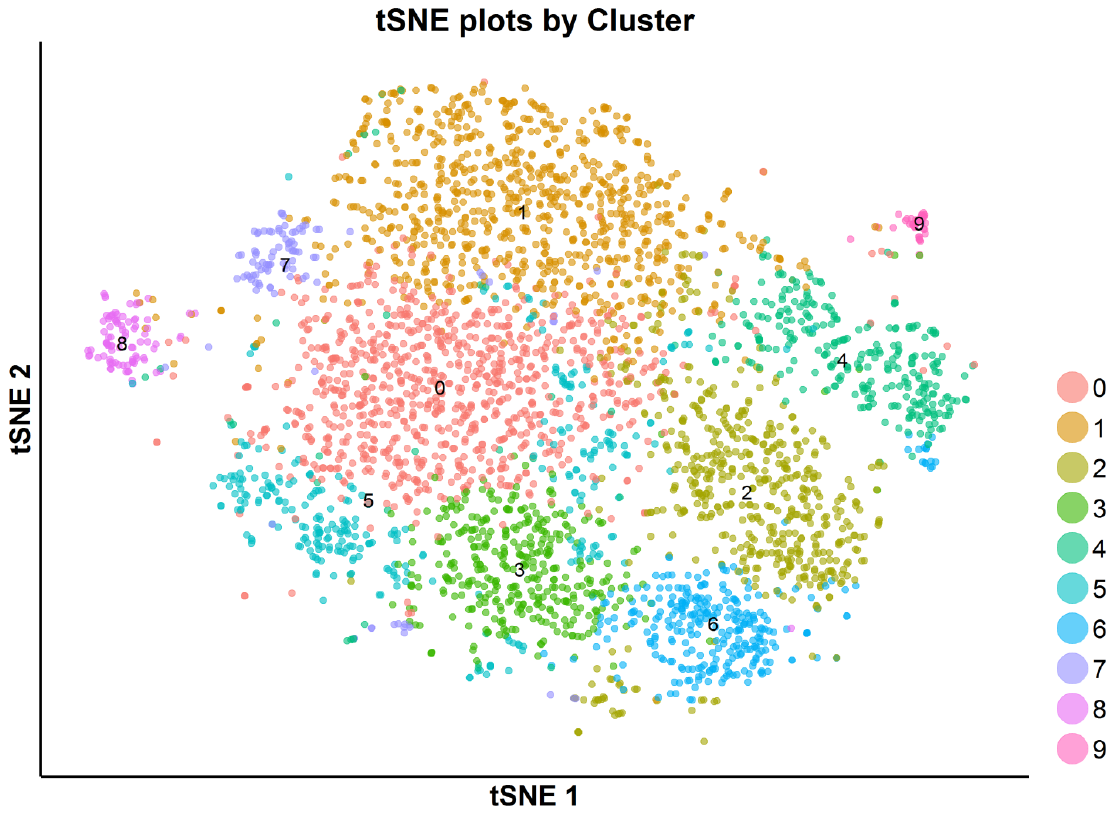

**Figure S51.** tSNE plot of the subpopulations of invasion cells. The invasion cells were clustered into 10 subpopulations with the resolution granularity of 1.0.

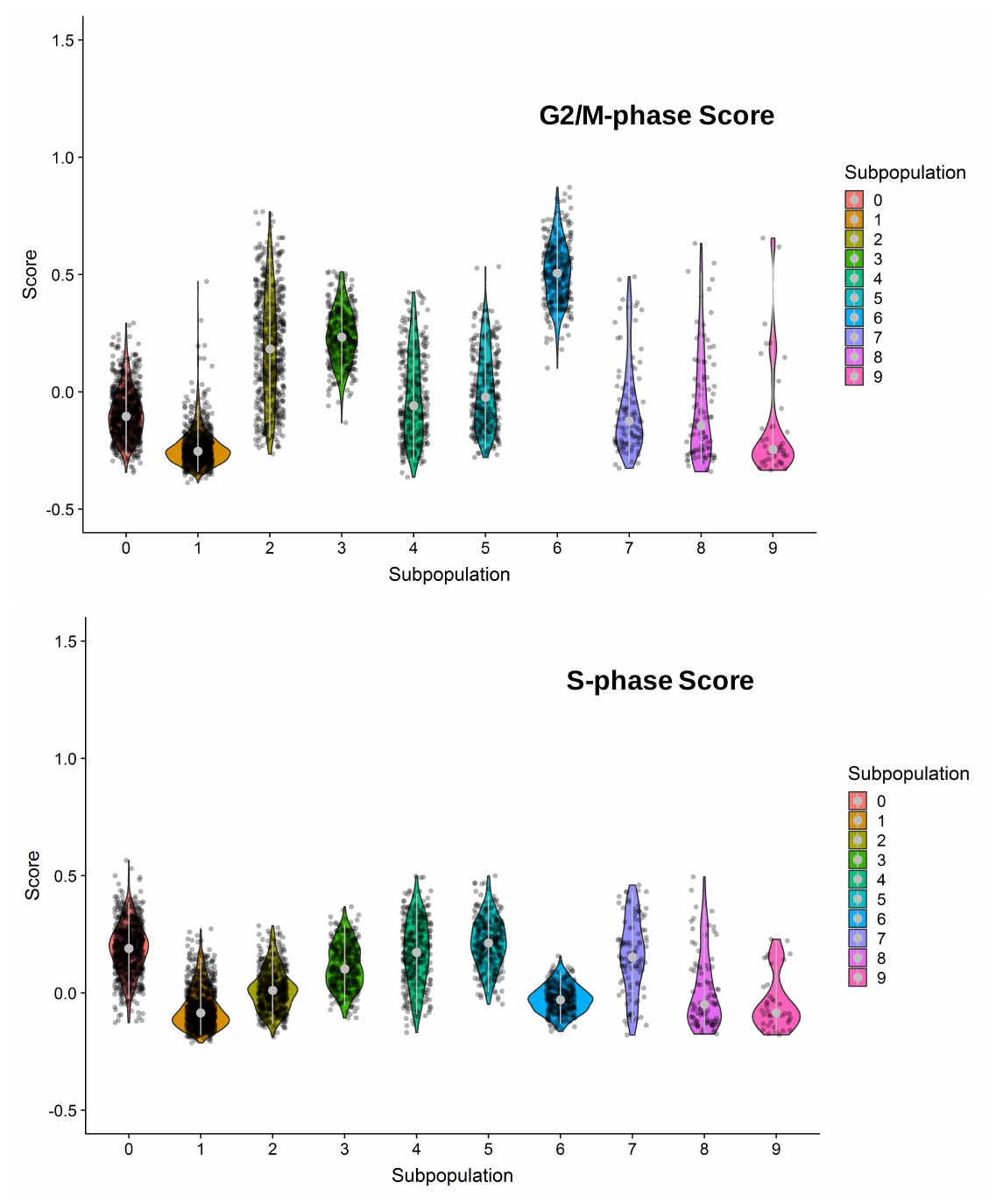

**Figure S52.** G2/M- and S-phase scores of the 10 clustered subpopulation of invasion cells. Subpopulation 0, 2, 3 and 6 were dominated by G2/M- or S-phase scores.

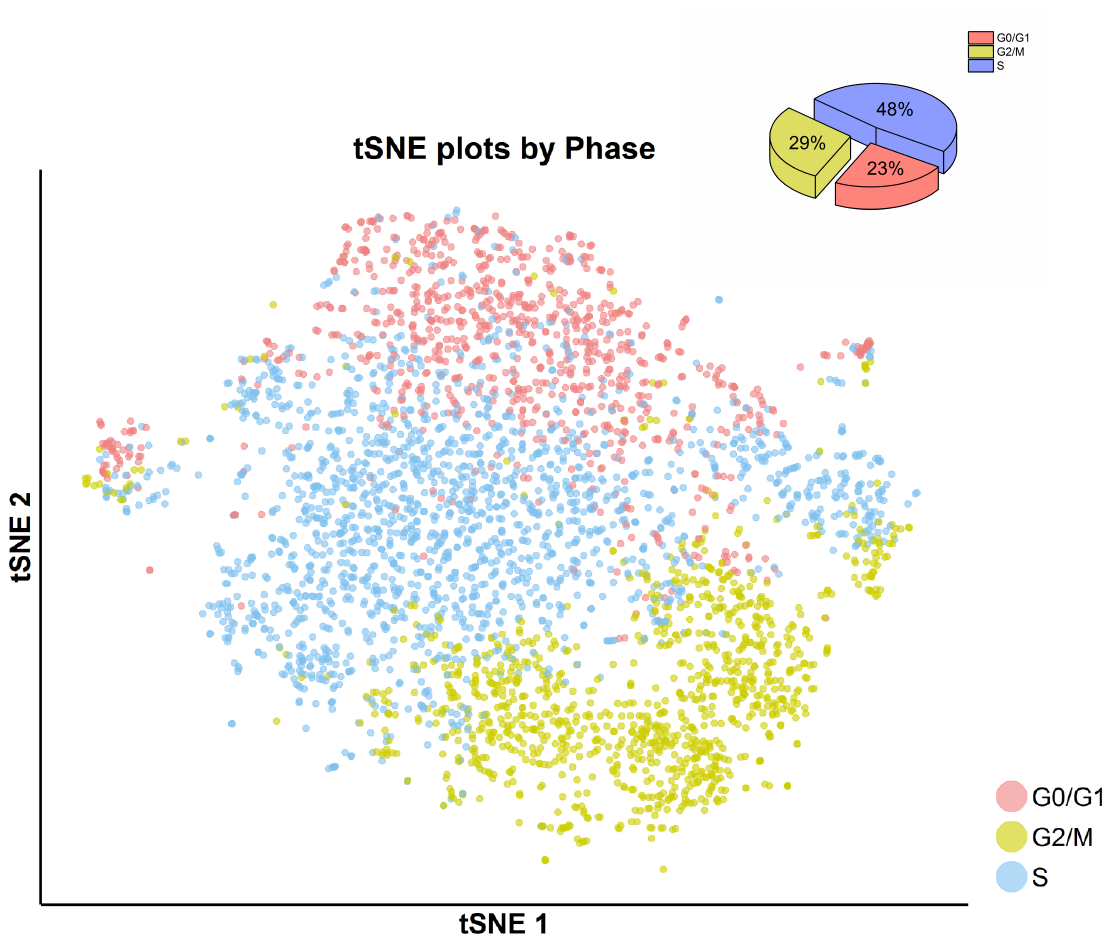

**Figure S53.** tSNE plot (center) with cell-cycle awareness and cell-cycle assignment (right upper corner) of invasion cells. Individual cells were classified according to their cell cycle scores as described in Supplementary Methods. Invasion cell showed more S- and G2/M-phase characters in comparison with the cells treated with DMSO, **SKI-73N** and **SKI-73**.

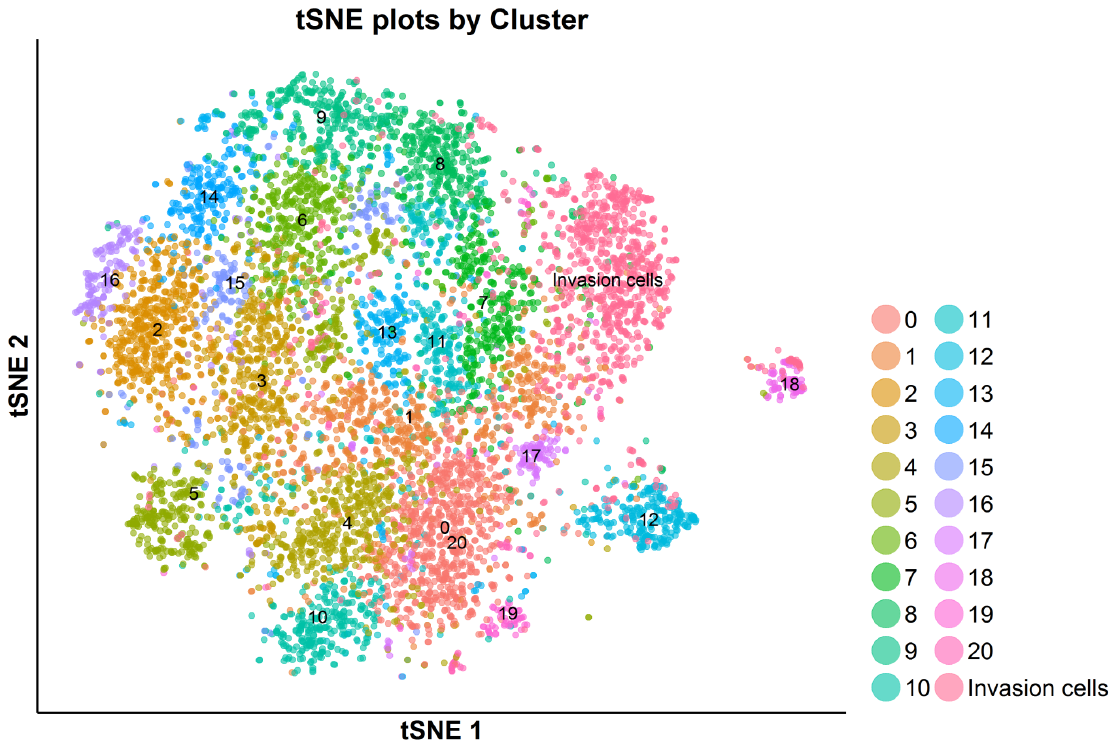

**Figure S54.** tSNE plot of the total G0/G1-phase cells analyzed by scRNA-seq---**SKI-73N-**, DMSO**-** and **SKI-73**-treated cells and invasion cells. The G0/G1-phase cells upon the treatment with **SKI-73N**, DMSO and **SKI-73** were clustered into 21 subpopulations as described in Figure 28.

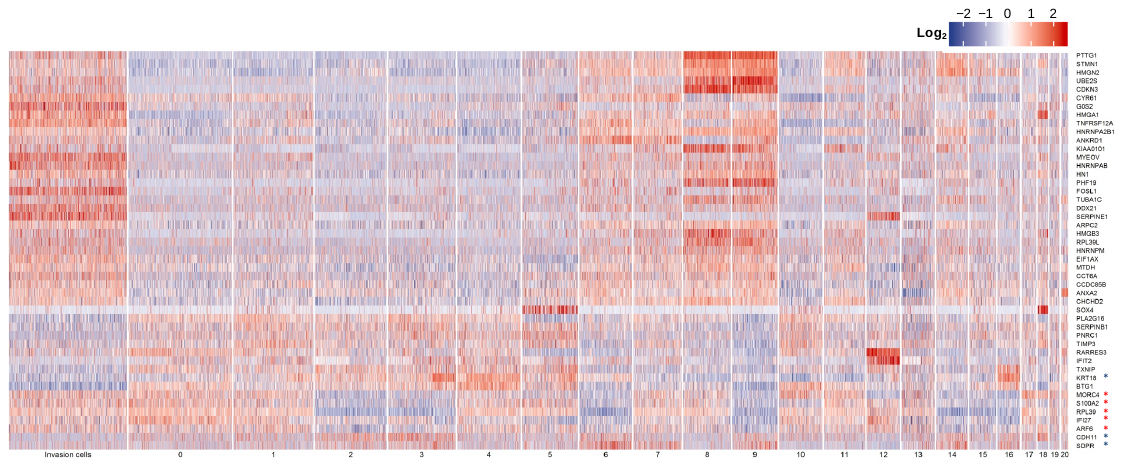

**Figure S55.** Heatmap of representative cancer-associated genes for comparison between invasion cells and the 21 subpopulations of G0/G1 cells. From top to bottom, the top 20 up-regulated and then top 10 down-regulated transcripts in invasion cells with implicated cancer relevance as revealed by differential analysis and listed in Table S8. From the bottom, the top 2 down-regulated (blue star) and the top 4 up-regulated transcripts (red star) in both invasion cells and the most invasion-prone cells (Subpopulation 8 of G0/G1 cells) with implicated cancer relevance as revealed by differential analysis and listed in Table S12. “KRT18” is the top down-regulated transcript shared in Table S8 and S12.

**Supplementary Tables S2-4, S13 (see the separate Excel files for S5-12)**

**Table S2.** Direct hydrogen bond interactions of **1** with the residues of CARM1 and their comparison with CARM1-**SNF** and CARM1-**SAH** complexes. The variations among the subunits of the CARM1-**1** complex (4IKP) were observed for the NZ of Chain A (III and IV) and Q160 in Chain C (´A and ´B). For the CARM1-**SNF** complex (2Y1W) and the CARM1-**SAH** complex (2Y1X), there is no significant difference among individual subunits. Bond distances of Å are displayed in parenthesis and the specific interacting atoms were shown as the following: N and O for backbone nitrogen and oxygen; OG for the side chain oxygen of Ser; OH for the side chain hydroxyl group of Tyr; OD1 and OD2 for the chain carbonyl oxygen and the side chain carboxylic oxygen of Asp; Gl and OE2 for the side chain carbonyl oxygen and the side chain carboxylic oxygen of Glu; NE2 for the side chain nitrogen of Asn or Gln; NH1/NH2 for side chain nitrogens of Arg.

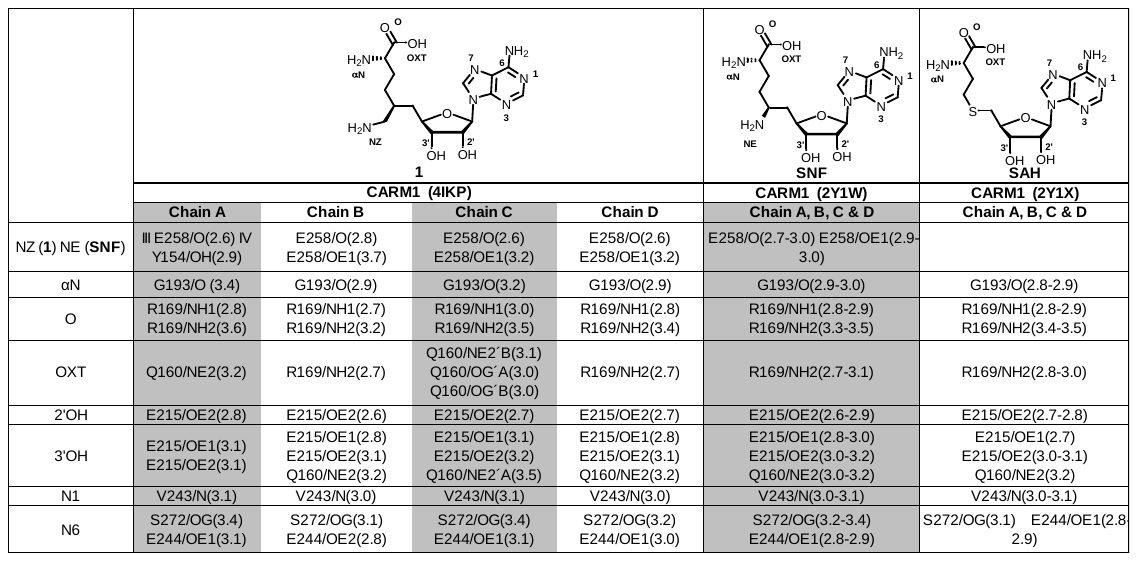

**Table S3.** Comparison of water hydrogen bonds of **1** and **SNF** in complex with human CARM1 (PDB 4IKP and 2Y1W). The key variations of the water hydrogen bonds with **1** in comparison with **SNF** were observed at the former’s NZ and OXT and CARM1’s Q160 region. Bond distances of Å are displayed in parenthesis and the specific interacting atoms were shown as the following: N and O for backbone nitrogen and oxygen; OG for the side chain oxygen of Ser; OH for the side chain hydroxyl group of Tyr; OD1 and OD2 for the chain carbonyl oxygen and the side chain carboxylic oxygen of Asp; Gl and OE2 for the side chain carbonyl oxygen and the side chain carboxylic oxygen of Glu; NE2 for the side chain nitrogen of Asn or Gln; NH1/NH2 for side chain nitrogen atoms of Arg.

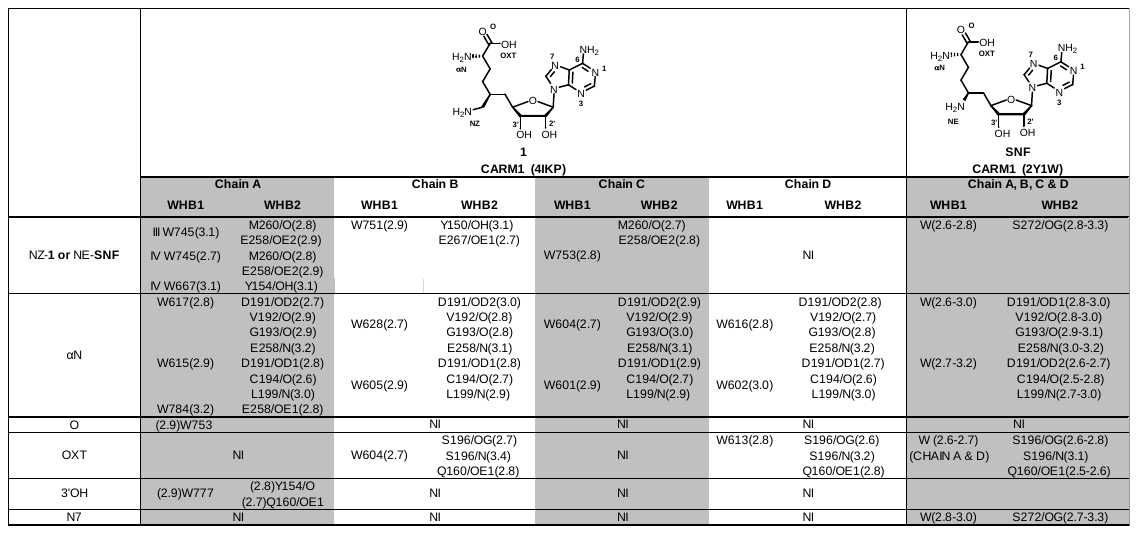

**Table S4.** Water hydrogen bonds of **SAH** in complex with CARM1 (PDB 2Y1X). The water hydrogen bonds of **SAH** in complex with CARM1 are shown with the bond distances of Å in parenthesis and the specific interacting atoms as the following: N and O for backbone nitrogen and oxygen; OG for the side chain oxygen of Ser; OH for the side chain hydroxyl group of Tyr; OD1 and OD2 for the chain carbonyl oxygen and the side chain carboxylic oxygen of Asp; Gl and OE2 for the side chain carbonyl oxygen and the side chain carboxylic oxygen of Glu; NE2 for the side chain nitrogen of Asn or Gln; NH1/NH2 for side chain nitrogen atoms of Arg.

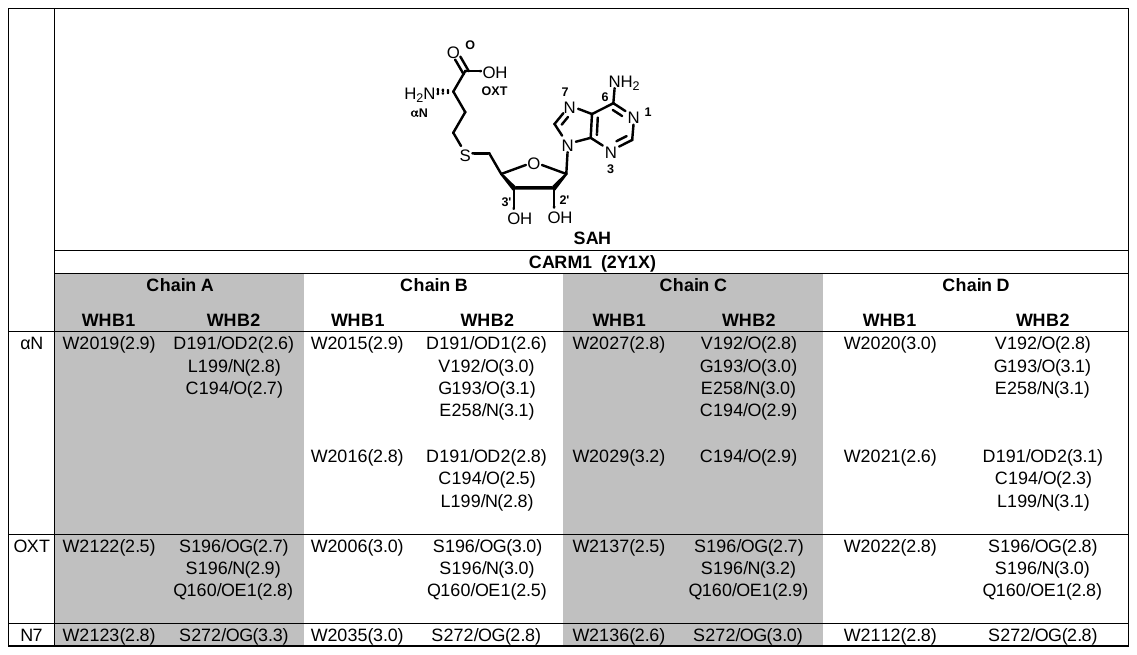

**Table S13.** Crystallography data and refinement statistics of the X-ray structures of CARM1 in complex with **1** and **5a**.

| **Ligands** | **1** | **5a** |
| --- | --- | --- |
| **PDB Code** | **4IKP** | **6D2L** |
| **Data collection** |  |  |
| Wavelength (Å) | 1.03321 | 0.97918 |
| Space group | P2_1_2_1_2_1_ | P2_1_ |
| **Cell dimensions** |  |  |
| *a*, *b*, *c* (Å) | 75.1,98.8,206.6 | 75.6,155.6,95.3 |
| α, β, γ (°) | 90.0,90.0,90.0 | 90.0,101.0,90.0 |
| Resolution (Å) | 50.0-2.00 | 50.0-2.00 |
| Unique reflections | 104,330 | 142,452 |
| Redundancy | 8.1 | 4.5 |
| Completeness (%) | 99.8 | 97.0 |
| I/σ(I) | 30.4 | 10.4 |
| R_sym_^a^ | 0.086 | 0.155 |
| R_pim_ | 0.032 | 0.081 |
| **Refinement** |  |  |
| No. protein molecules/ASU | 4 | 6 |
| Resolution (Å) | 48.1-2.00 | 50.0-2.00 |
| Reflections used or used/free | 103,958 | 139,748/1,400 |
| Rwork(%) | 20.3 | 18.7 |
| Rfree(%) | 23.1 | 23.6 |
| Average B value (Å^2^) | 33.9 | 30.8 |
| *Protein* | 33.4 | 30.5 |
| *Compound* | 29.4 | 34.8 |
| *Other* | n/a | n/a |
| *Water* | 42.1 | 35.5 |
| Number of Atoms | 11,635 | 16,868 |
| *Protein* | 10,770 | 15,819 |
| *Compound* | 117 | 253 |
| *Other* | n/a | n/a |
| *Water* | 748 | 766 |
| RMS Bonds (Å) | 0.007 | 0.008 |
| RMS Angles (°) | 1.127 | 1.273 |
| Wilson B (Å^2^) | 33.9 | 29.6 |
| **Ramachandran plot** |  |  |
| Most favoured (%) | 96.9 | 97.1 |
| Additional allowed (%) | 3.03 | 2.9 |
| Outliers (%) | 0.07 | 0.0 |
